# Lifetime brain atrophy estimated from a single MRI: measurement characteristics and genome-wide correlates

**DOI:** 10.1101/2024.11.06.622274

**Authors:** Anna E. Fürtjes, Isabelle F. Foote, Charley Xia, Gail Davies, Joanna Moodie, Adele Taylor, David C. Liewald, Paul Redmond, Janie Corley, Andrew M. McIntosh, Heather C. Whalley, Susana Muñoz Maniega, Maria Valdés Hernández, Ellen Backhouse, Karen Ferguson, Mark E. Bastin, Joanna Wardlaw, Javier de la Fuente, Andrew D. Grotzinger, Michelle Luciano, W. David Hill, Ian J. Deary, Elliot M. Tucker-Drob, Simon R. Cox

## Abstract

A measure of lifetime brain atrophy (LBA) obtained from a single magnetic resonance imaging (MRI) scan could be an attractive candidate to boost statistical power in uncovering novel genetic signals and mechanisms of neurodegeneration. We analysed data from five young and old adult cohorts (MRi-Share, Human Connectome Project, UK Biobank, Generation Scotland Subsample, and Lothian Birth Cohort 1936 [LBC1936]) to test the validity and utility of LBA inferred from cross-sectional MRI data, i.e., a single MRI scan per participant. LBA was simply calculated based on the relationship between total brain volume (TBV) and intracranial volume (ICV), using three computationally distinct approaches: the difference (*ICV-TBV*), ratio (*TBV*/*ICV*), and regression-residual method (TBV∼ICV). LBA derived with all three methods were substantially correlated with well-validated neuroradiological atrophy rating scales (*r* = 0.37-0.44). Compared with the difference or ratio method, LBA computed with the residual method most strongly captured phenotypic variance associated with cognitive decline (*r* = 0.36), frailty (*r* = 0.24), age-moderated brain shrinkage (*r* = 0.45), and longitudinally-measured atrophic changes (*r* = 0.36). LBA computed using a difference score was strongly correlated with baseline (i.e., ICV; *r* = 0.81) and yielded GWAS signal similar to ICV (*r_g_* = 0.75). We performed the largest genetic study of LBA to date (*N* = 43,110), which was highly heritable (*h^2^* _SNP GCTA_ = 41% [95% CI = 38-43%]) and had strong polygenic signal (LDSC *h^2^* = 26%; mean *χ2* = 1.23). The strongest association in our genome-wide association study (GWAS) implicated *WNT16*, a gene previously linked with neurodegenerative diseases such as Alzheimer, and Parkinson disease, and amyotrophic lateral sclerosis. This study is the first side-by-side evaluation of different computational approaches to estimate lifetime brain changes and their measurement characteristics. Careful assessment of methods for LBA computation had important implications for the interpretation of existing phenotypic and genetic results, and showed that relying on the residual method to estimate LBA from a single MRI scan captured brain shrinkage rather than current brain size. This makes this computationally-simple definition of LBA a strong candidate for more powerful analyses, promising accelerated genetic discoveries by maximising the use of available cross-sectional data.

## Introduction

The loss of brain matter due to ageing-related neurodegeneration, subsequently referred to as brain atrophy, is an important feature of non-pathological ageing (Scahill et al., 2003), late-life cognitive decline (e.g., Raz & Rodrigue, 2006), and accelerated neurodegenerative diseases such as dementia (Good et al., 2002). Brain atrophy can be observed in structural magnetic resonance imaging (MRI) scans, and is characterised by the widening of sulci, loss of gyral volume and the concomitant enlargement of ventricles as the brain shrinks away from the intracranial vault and cedes space to increasing cerebrospinal fluid volume. As a dynamic within-person phenomenon, brain atrophy may be most adequately inferred with repeated brain MRI scans, through which individual-level brain matter volume changes can be tracked over time (Lindenberger & Ghisletta, 2009; Raz et al., 2005; Salthouse, 2011), and phenotypic and genetic predictors can be identified. However, the high costs and inconvenience of repeated MRI scanning can be prohibitive for longitudinal MRI data collection at large scale. Whereas there are several ongoing initiatives that will result in large-scale collection of longitudinal scans with other data (including genetic information), a reliable method of estimating brain atrophy from a single-occasion MRI scan of a person’s brain would substantially boost sample sizes and statistical power for scientific discovery in the shorter-term.

This is particularly true in the context of genome-wide association studies (GWAS) – for which many thousands of participants are required (Abdellaoui et al., 2023) – a measure of total brain atrophy obtainable from a single cross-sectional MRI scan is an especially attractive candidate for boosting statistical power. It could help uncover novel genetic signals and molecular mechanisms of neurodegeneration. Current meticulously collected longitudinal brain GWAS efforts remain understandably small and lacking in statistical power (Brouwer et al., 2022; N<16,000), given the rarity of longitudinal brain MRI studies with genetic data. Here, we present a phenotypic and genetic analysis of multiple approaches to derive an estimate of lifetime brain atrophy (LBA) from a single MRI scan.

In neuroradiological settings, it is common practice for trained experts to manually rate brain atrophy from a single MRI scan using well-validated visual scales (Koedam et al., 2011; Pasquier et al., 1996; Scheltens et al., 1995). Computational approaches concur with visual rating scales, but offer greater statistical power and greater fidelity to index adulthood atrophic changes (Velickaite et al., 2020). LBA can be computed in middle-aged and older adults from a single-occasion brain MRI scan using measures of total intracranial volume (ICV) and total brain volume (TBV). Such an approach to LBA compares how much brain tissue an individual currently has (TBV) as some function of prior (or pre-neurodegenerative) brain volume – inferred from ICV (Miller et al., 2002; Rudick et al., 1999; Vågberg et al., 2017; Whitwell et al., 2001). Supported by the evidence that ICV remains broadly stable across the lifespan (Caspi et al., 2020; Holmes et al., 2015; Royle et al., 2013) – which is not true for ageing-related reductions in the volume of the brain – one can employ ICV as an ‘archaeological index’ of an individual’s maximum prior brain size. Using this as a baseline, we may estimate LBA by comparing TBV and ICV in middle-aged and older adults^1^.

However, there are different methods for comparing ICV and TBV to estimate a person’s LBA. Here, we test three commonly-used computational approaches to derive LBA from ICV and TBV: the difference, ratio, and regression-residual methods (*Box S1*). We show that these three methods produce estimates with starkly different statistical properties, which has substantial implications for the resulting atrophy estimate and its interpretation. All three computational approaches have previously been used to estimate brain changes (or other changes between two time points; e.g., Brouwer et al., 2022; Wang et al., 2024), but to our knowledge no study has provided a side-by-side evaluation of the resulting measurement characteristics. It is unclear which method most clearly reflects age-associated brain shrinkage or longitudinally-observed atrophic changes, and whether the methods differentially account for ageing-related health traits. Furthermore, it remains unknown whether the different approaches capture variance associated with distinct genetic signal, and distinct genetic overlap with other ageing-related traits. Specifically, there is a need to establish whether using the difference method mainly captures polygenic signal associated with ICV, and whether the inclusion of ICV (or another strongly related proxy of skull size) as a covariate in a TBV GWAS captures genetic variance associated with brain atrophy rather than just initial brain or overall head/skull size.

In this pre-registered study (https://osf.io/gydmw/) we leveraged five large-scale cohorts to conduct the largest genetic study to-date of LBA in middle- and older adulthood (*Fig.1*). First, we phenotypically validated LBA, showing it was substantially associated with neuroradiological atrophy ratings. Out of the three computational approaches, the residual method captured maximal associations with cognitive decline, frailty, age-moderated brain shrinkage, and longitudinally-measured atrophic changes. Second, we performed the largest LBA genome-wide association study (GWAS) to date (*N* = 43,110) and show that LBA as a residual exhibited substantial genetic signal independent of ICV; it was highly heritable and had strong polygenic signal. The most significant GWAS genomic risk locus implicated *WNT16*, a gene previously linked with neurodegeneration. In contrast, LBA as a difference score mostly indexed ICV, suggesting that it has far-reaching implications for future longitudinal and cross-sectional GWAS studies of brain ageing that modelling change with a difference score approach mainly captured baseline variance (i.e., skull size) rather than brain matter shrinkage.

**Fig. 1.**
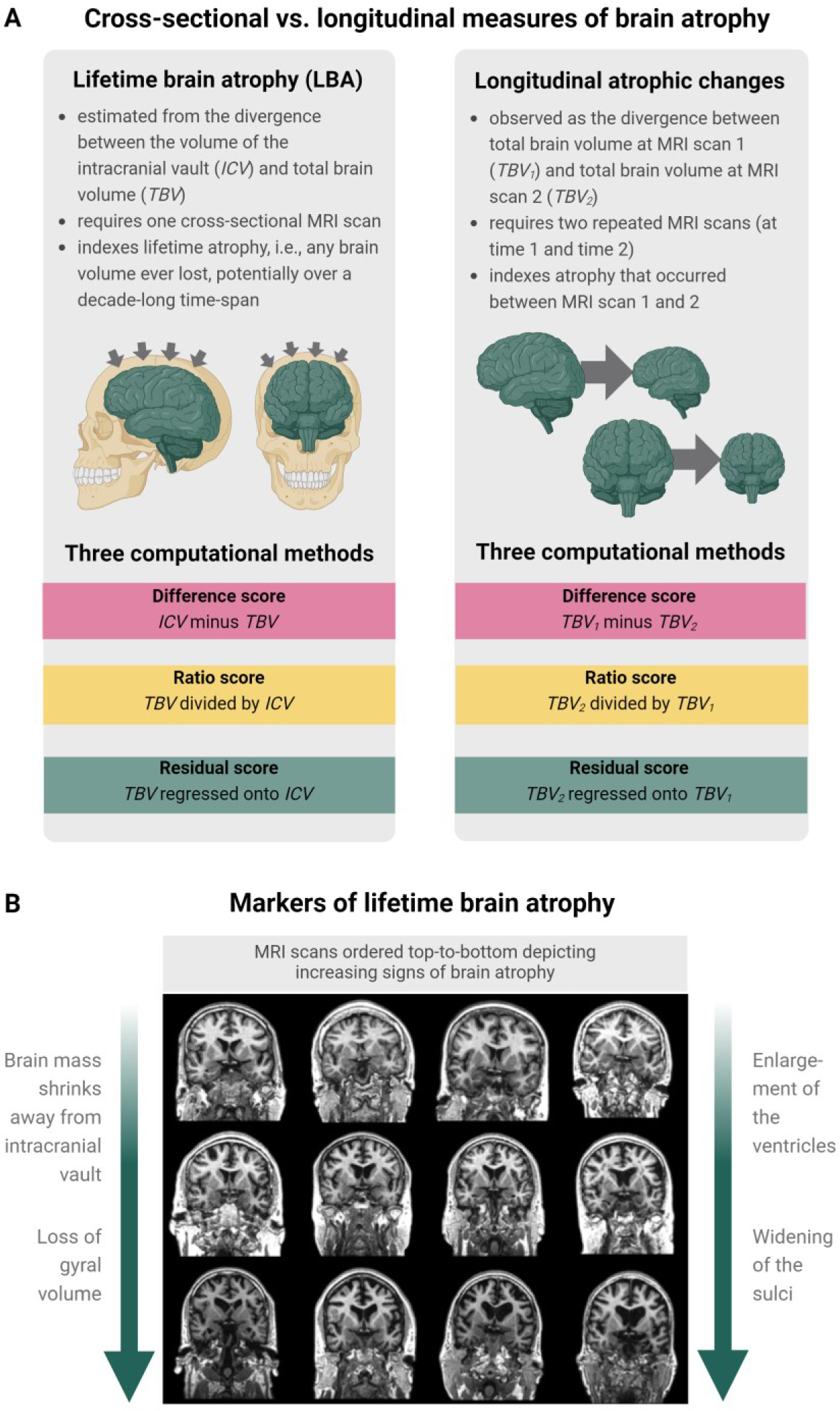
**A.** Illustration of the differences between cross-sectionally-estimated LBA and longitudinally-observed atrophic changes and the three distinct computational methods used to calculate brain atrophy (top panel). Illustration was produced in BioRender. **B.** The bottom panel shows MRI scans ordered top-to-bottom depicting increasingly strong signs of brain atrophy. Both visual rating scales and computational methods can be applied to estimate brain atrophy from a single neuroimaging scan such as this. Coronal and axial brain MRIs are T1-W slices of 73-year-old adults from the Lothian Birth Cohort 1936, adapted from Cox and Deary (2022) with permission under CC-BY 4.0..

## Results

### Description and characterisation of the LBA phenotype

We initially sought to validate measures of LBA estimated from a single MRI scan using the difference (*ICV-TBV*), ratio (*TBV*/*ICV*), and regression-residual method (TBV∼ICV). We reasoned that LBA should 1) correlate with atrophy rated by neuroradiologists, and with other brain- and ageing-related phenotypes; 2) be greater in cross-sectionally scanned older adults (on average), and increase longitudinally with age; and 3) capture observed atrophic changes in longitudinal data. First, we derived, characterised and phenotypically validated the three LBA measures estimated from a single MRI scan in middle- and older-aged adults. For basic measurement characteristics across five cohorts (MRi-Share, Human Connectome Project, UK Biobank, Generation Scotland, and LBC1936), *SMaterials* depict descriptive statistics (*Table S1*), variable distributions coloured by age (*SFig.3*), and Pearson’s correlations between TBV, ICV, cerebrospinal fluid (CSF) volume and LBA derived with the difference, ratio, and residual method (e.g., correlation between ICV and LBA in LBC1936: *r* _difference_ = 0.81 [95%CI = 0.76-0.84], *r* _ratio_ = 0.49 [95%CI = 0.40-0.57], *r* _residual_ = 0.00 [95%CI = −0.12-0.11]).

#### 1. Measures of LBA predict brain atrophy rated by neuroradiological experts, as well as other ageing-related health traits such as frailty and cognitive ability

Since the difference, ratio, and residual method produced LBA measures with quite different statistical properties, we compared their associations with visually-rated atrophy and other ageing-relevant phenotypes in two cohorts with repeated MRI measures: the LBC1936 (*N* = 286, 71-73 years of age) and the UK Biobank (UKB; *N* = 4,674, 46-81 years of age). *Fig.2* summarises associations in LBC1936. *Cognitive ability*, *frailty*, and *body mass index* (BMI) were modelled using growth curve models to extract individual-level intercept (e.g., *iCog*) and slope values (e.g., *sCog*), to capture variance associated with stable baseline characteristics and change, respectively (*Methods*).

**Fig. 2.**
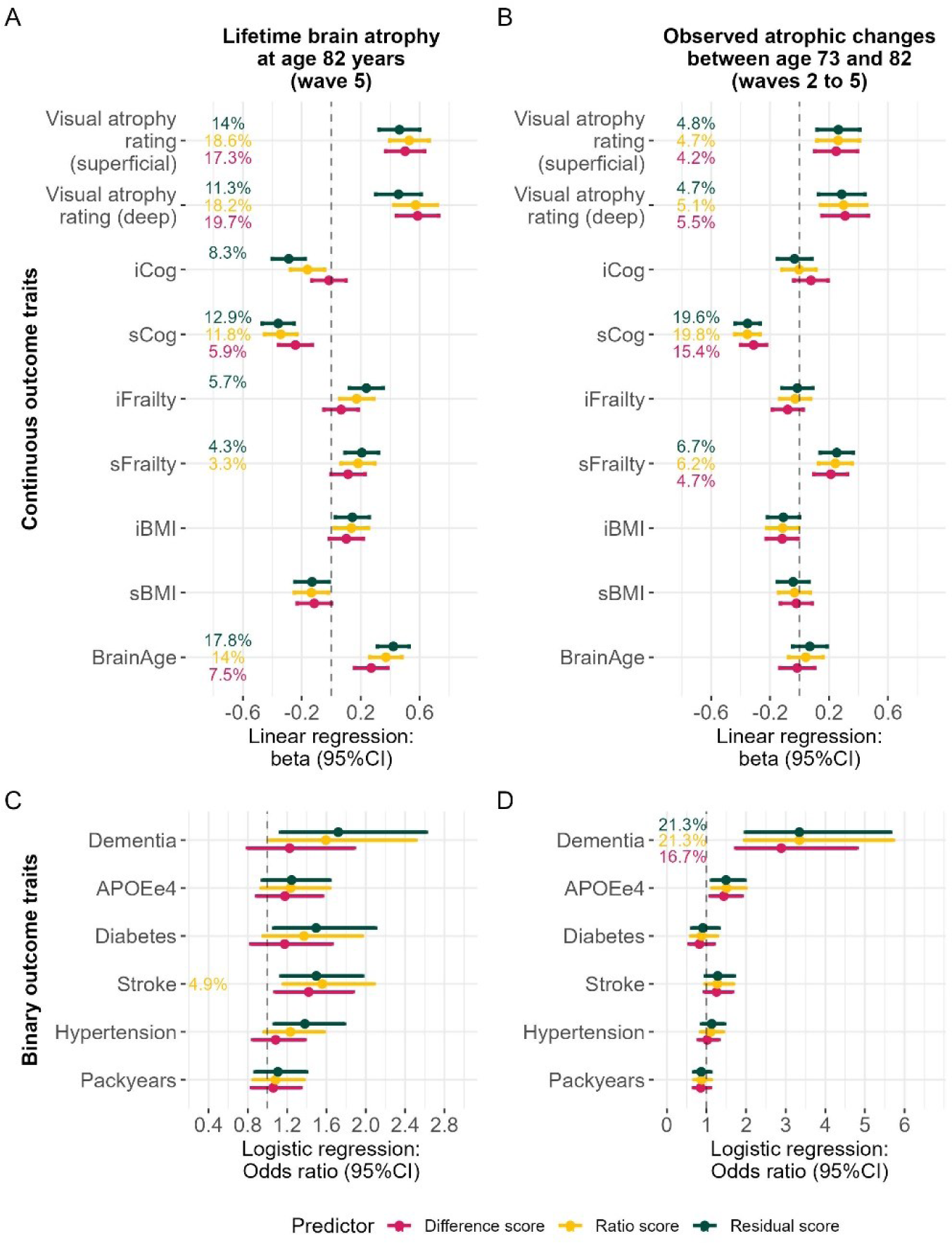
Associations with health-related phenotypes in LBC1936 for LBA and observed atrophic changes. Atrophy estimated with the ratio and residual method were flipped to match the difference score whereby larger values represent more brain atrophy. Associations with continuous traits in panels A-B were calculated with linear regressions, where the outcome was always one of the traits listed on the *y*-axes and the predictor was either LBA or longitudinal atrophic changes. It may be counterintuitive to include *APOE* as an outcome, but we make no claims of causality or directionality and prioritised treating all outcome traits equally. We outline in the *Methods* how outcome traits were derived. Beta effect sizes indicate change per SD in the health trait (e.g., cognitive ability). Percentages indicate variance explained (*R^2^*) in the health trait, and is only printed if the association is statistically significant (*p* < 0.05/ 15 traits). Associations with binary traits in panel C-D were calculated with logistic regressions; *R^2^* estimates were obtained with Nagelkerke’s *R^2^*. Odds ratios indicate the increased chances of having one of the diseases listed on the *y*-axis associated with one SD deviation in LBA (or observed atrophic changes). Only ‘packyears’ was analysed with hurdle regression where *R^2^* is inferred with maximum likelihood pseudo *R^2^*. Confidence intervals are at 95%. *R^2^* was only printed if the association was statistically significant (*p* < 0.05/ 15 traits). Note when interpreting the results that the *x*-axes on panel C-D are scaled differently. Variables for which we extracted intercepts and slopes (Cog, Frailty, BMI) were relative to the same baseline as observed atrophic changes but LBA represents loss since maximum brain size many years earlier.

First, we externally validated LBA by demonstrating substantial associations between LBA and brain atrophy rated by well-validated (Farrell et al., 2009) neuroradiological rating scales (*beta* _residual_ = 0.46 [95%CI = 0.32-0.60], *beta* _ratio_ = 0.53 [95%CI = 0.39-0.66], *beta* _difference_ = 0.50 [95%CI = 0.37-0.63]). Such visual rating scales were used by trained experts to manually rate brain atrophy from a single MRI scan in LBC1936 (description in *SMaterials*). Associations between longitudinally-observed atrophic changes and visually-rated atrophy were smaller (*beta* _residual_ = 0.26 [95%CI = 0.12-0.41], *beta* _ratio_ = 0.26 [95%CI = 0.11-0.41], *beta* _difference_ = 0.25 [95%CI = 0.10-0.39]), and there was no association between TBV and visually-rated atrophy (*beta* _TBV_ = −0.07 [95%CI = - 0.22-0.07]).

Second, associations with other ageing-related traits were similarly strong for LBA (*Fig.2A&C*) as they were for longitudinally-observed atrophic changes (*Fig.2B&D*) in LBC1936. The degree to which LBA predicted cognitive function, frailty, and brain age, was dependent upon the LBA computation method, and below we only report LBA results from the residual method because it consistently outperformed the difference and ratio method. Greater LBA explained sizable percentages of variances in, and was associated with lower cognitive ability (i.e., intercept, *iCog* in *Fig.2*; *beta* _residual_ = −0.29 [95%CI = −0.40 – −0.18]) as well as steeper rates of cognitive decline (i.e., slope, *sCog*; *beta* _residual_ = −0.36 [95%CI = −0.47 – −0.25]), greater baseline frailty (i.e., intercept; *beta* _residual_ = 0.24 [95%CI = 0.13-0.35]), greater longitudinal increases in frailty (i.e., slope; *beta* _residual_ = 0.21 [95%CI = 0.09-0.32]), and a larger brain age gap (i.e., brains appearing older given chronological age; *beta* _residual_ = 0.42 [95%CI = 0.31-0.53]). Given participants in the LBC1936 were assessed at the age of 73 years, the cognitive intercept variable (*iCog*) likely captured not only baseline levels of cognitive function, but also decades-worth of cognitive decline. It validates LBA _residual_ as a marker of cognitive and brain *ageing* that it predicted the cognitive intercept significantly (*beta* _residual_ = −0.29 [95%CI = −0.40 – −0.18]) when LBA _difference_ (*beta* _difference_ = −0.02 [95%CI = −0.13 – 0.10]; *ns.*) and LBA _ratio_ did not (*beta* _ratio_ = −0.16 [95%CI = −0.28 – −0.05]; *ns.*).

In the same LBC1936 participants, those associations differed when the predictor variable (i.e., LBA) was replaced with longitudinal atrophic changes (observed between ages 73 to 82 years) in that observed atrophic changes only indexed inter-individual *changes* across this same 9-year period (i.e., slopes rather than intercepts): Those 9-year atrophic changes were associated with steeper rates of cognitive decline (*beta* _residual_ = −0.35 [95%CI = −0.44 – −0.27]) and steeper rates of worsened frailty (*beta* _residual_ = 0.25 [95%CI = 0.14-0.37]), but not baseline levels of cognitive ability or frailty (i.e., intercepts). Clinically-ascertained all-cause dementia (Mullin et al., 2023) was less strongly associated with LBA (*p* = 0.011 did not survive correction for multiple testing; *OR* _residual_ = 1.72 [95%CI = 1.13-2.61]) than it was associated with the longitudinal 9-year atrophic changes (*OR* _residual_ = 3.34 [95%CI = 1.98-5.65]). This may reflect that dementia patients exhibit the steepest rates of disease-caused brain shrinkage in years proximal to diagnosis– which we capture more directly with longitudinal MRI measures across the 9-year period. By contrast, the broader LBA phenotype might index longer-term changes of ageing that are not specifically disease-related and therefore less precisely distinguish participants with dementia diagnoses.

In the UKB sample, LBA was significantly associated with greater cognitive ability (*beta* _residual_ = 0.23 [95%CI = 0.20-0.26]), older brain age gap (*beta* _residual_ = 0.27 [95%CI = 0.24-0.30]), more severe frailty (*OR* _residual_ = 1.24 [95%CI = 1.16-1.32]), increased likelihood to have diabetes (*OR* _residual_ = 1.51 [95%CI = 1.33-1.71]), and hypertension (*OR* _residual_ = 1.46 [95%CI = 1.37-1.56]; *SFig.7*)^2^. Associations were significant despite the extremely healthy nature of UKB neuroimaging participants: raw LBA estimates suggested UKB participants were similarly healthy to those in the MRi-Share and HCP cohorts who were chronologically four decades younger, on average (*SFig.1-2*).

#### 2. Measures of LBA indicate age-associated brain shrinkage

##### Single time-point MRI measures correlation with age

When considering only one single time-point, LBA estimated with all three methods was substantially correlated with age in the UKB, which is a mid-to-late life age-heterogeneous sample (46-81 years). We show that the strength of this correlation is moderated by sample age. That is, we created subsamples of successively older upper-age limits within a cohort, which caused the correlation between age and LBA to increase towards older adulthood (*Fig.3C*). For example, LBA _residual_ was correlated with age at *r* = 0.45 [95%CI 0.44-0.46] in the full UKB sample (*N* = 43,389; age range = 45-81 years). However, considering a subsample aged <62-years produced an age correlation of *r* = 0.1 [95%CI 0.08-0.11] (*N* = 15,461), and the same correlation was zero [95%CI −0.001-0.05] when considering only below 55-years-olds (*N* = 5,475). Results remained stable but yielded fewer significant age correlations when this analysis was repeated for males and females separately (*SFig.13*).

**Fig. 3.**
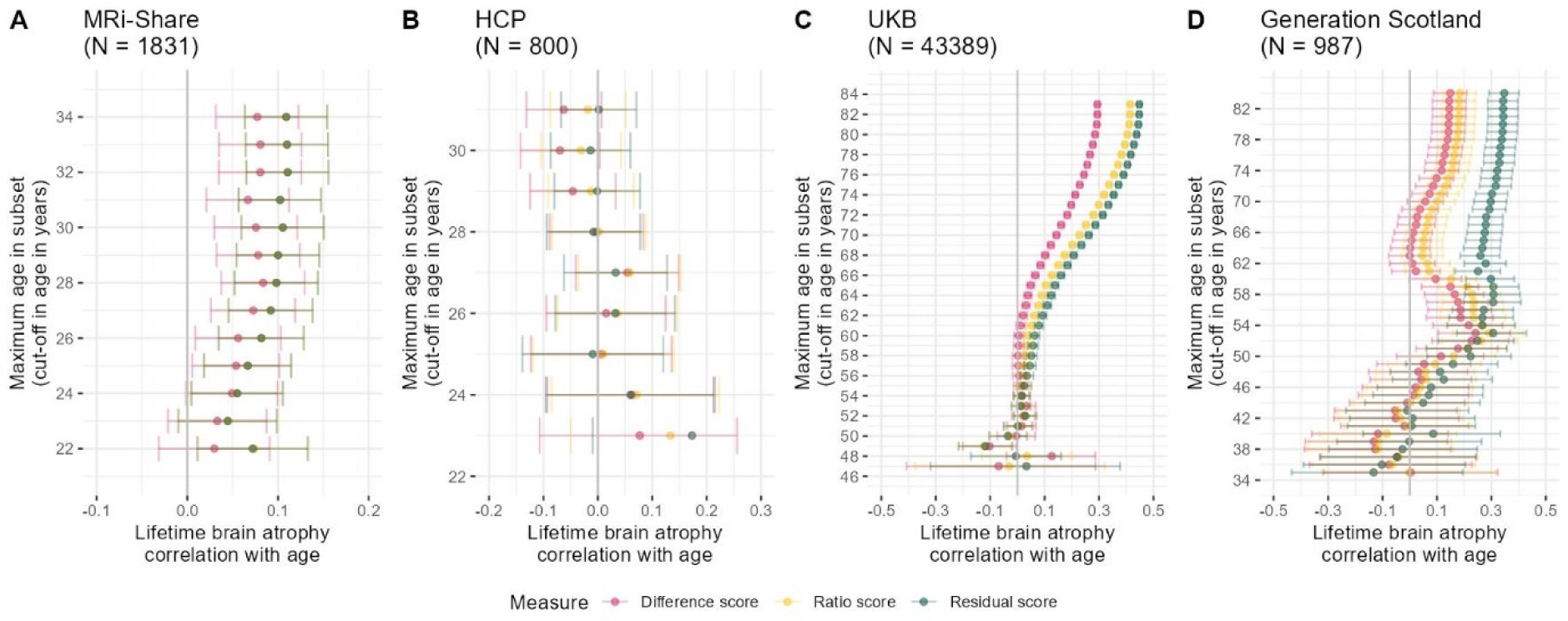
LBA is moderated by sample age across four earlier- and later-life cohorts. Age moderation of LBA estimated with the difference, ratio, and residual method. The *y*-axes indicate the maximum age in sample subsets taken to calculate the respective plotted age correlation. For example, a value of 35 on the *y*-axis means that the sample subset used to calculate the age correlation contains participants 35 years or younger. The ratio and residual scores have been flipped (i.e., multiplied by - 1) so a larger value corresponds to more brain atrophy. Panel A displays age correlations in the MRi-Share cohort (*N* = 1,831; age range 18-34 years), Panel B the HCP (*N* = 800; age range 22-31 years), Panel C the UKB (*N* = 43,389; age range = 45-83), and Panel D the Generation Scotland sample (*N* = 987; age range 26-84). The kink in the Generation Scotland diagram is likely a reflection of the bimodal distribution of the LBA phenotypes, which least strongly affected the residual score (*SFig.3*). This kink was likely driven by the males in the sample (*SFig.13*). Note that the mismatch in the reported UKB sample size in this Figure and *Table S1* is due to this Figure requiring one MRI scan only which is available from many more participants than two repeated measures as required by analyses of longitudinally-observed atrophic changes. Age correlations were non-significant when repeated in an unrelated HCP sample (*n* = 326; *SFig.14*).

This same trend was evident in the young-to-old-age Generation Scotland sample (*N* = 987; age range 26-84 years; *Fig.3D*) in that age correlations were strongest including older participants. This trend even existed in young adults, namely the MRi-Share cohort including mainly university students (*Fig.3A*, *N* = 1,831; age range 18-34 years), but not the HCP cohort (*Fig.3B, N* = 800; age range 22-31 years). Note that the raw HCP data unexpectedly showed strong age correlations for ICV (i.e., smaller skulls at older ages contradicting our pre-registered expectations; *r* = −0.20, *p* = 4.2×10^-11^; *SFig.13*) which seems to have been driven by 303 participants >31-years with small skulls that were excluded in the HCP data we analysed (e.g., in *Fig.3B*; *SMethods*). Additional analyses in the age-homogeneous and repeatedly MRI-scanned LBC1936 cohort showed that LBA increased with advancing chronological age, even when MRI measures at each visit were processed independently of each other (i.e., in this analysis MRI data was processed with the FreeSurfer cross-sectional rather than the longitudinal processing stream; *SFig.12* & description in *SMaterials*).

#### 3. LBA moderately captures longitudinally-observed atrophic changes over nine years

To quantify the extent to which LBA approximated longitudinally-observed atrophic changes, we derived atrophy scores from both cross-sectional MRI data measured at a second of two occasions (i.e., LBA), and from two longitudinal MRI scans measured nine years apart in LBC1936. We term the latter *observed atrophic changes*, capturing a time window shorter than, but overlapping with the one captured by *lifetime* BA (*Fig.1*). For all computational methods, the Pearson’s correlation was substantial, but numerically highest when atrophy was inferred with the residual method (LBC1936; *N* = 277): *r* _residual_ = 0.36 [95% CI = 0.25-0.46], *r* _ratio_ = 0.29 [95% CI = 0.18-0.40], *r* _difference_ = 0.30 [95% CI = 0.19-0.40]; *SFig.4*. This suggests that LBA moderately captured variation indicated by longitudinal atrophic changes, despite the fact that the two measures cover time windows of different lengths (i.e., lifetime vs. 9 years in LBC1936). The correlations appeared substantial given that LBC1936 participants demonstrated, on average, limited atrophic changes across the measured 9-year window (mean *TBV _time 1_* = 1,028 *mm^3^*, mean *TBV _time 2_* = 959 *mm^3^*; *SFig.10-11*) which likely limited the potential for a strong correlation between LBA and observed atrophic changes.

**Fig. 4.**
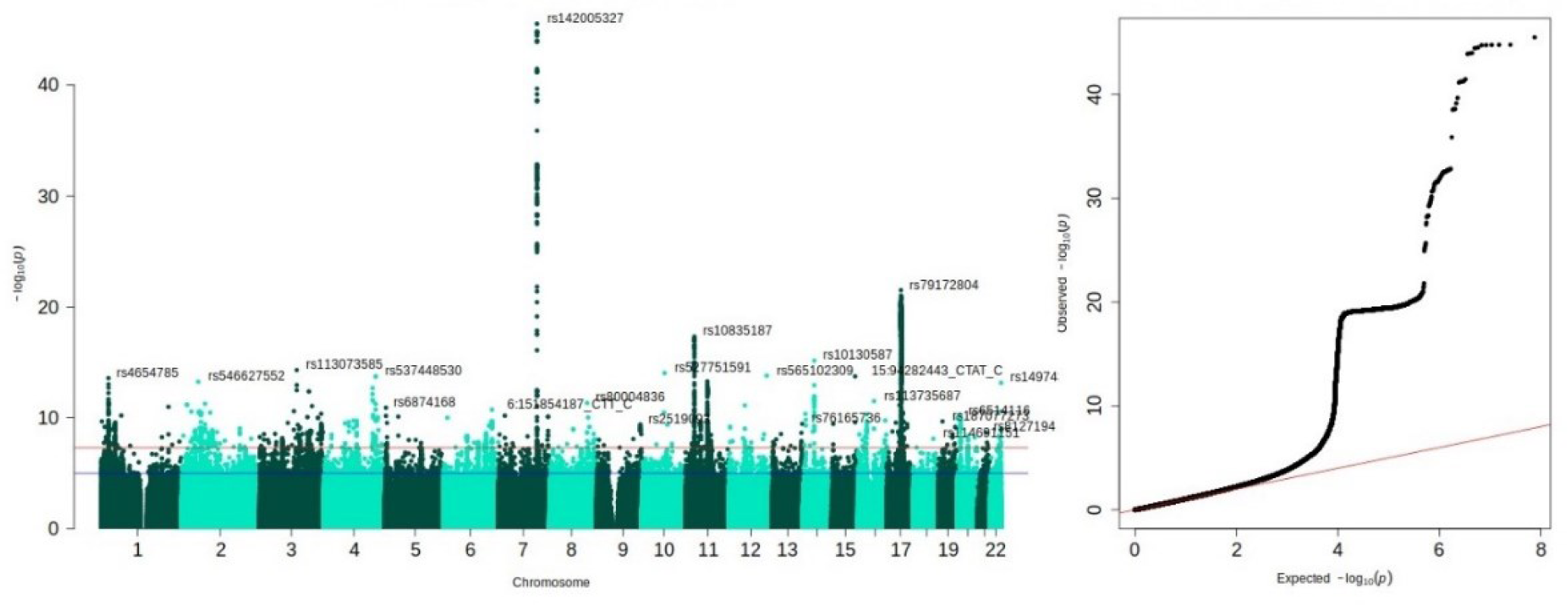
Left: Manhattan plot for LBA inferred with the residual method indicating top GWAS hits from the 28 associated loci. Right: *QQ* plot indicating deviation of polygenic signal from the expected null signal on the red line. The genomic inflation factor was 1.045. Both panels were plotted with the *qqman* package in R (Turner, 2018). Further evaluation and visualisation of the *QQ* plot in *SFig.33* which shows that the step in the *QQ*-plot shown here disappeared when considering HapMap3 SNPs only.

**Fig. 5.**
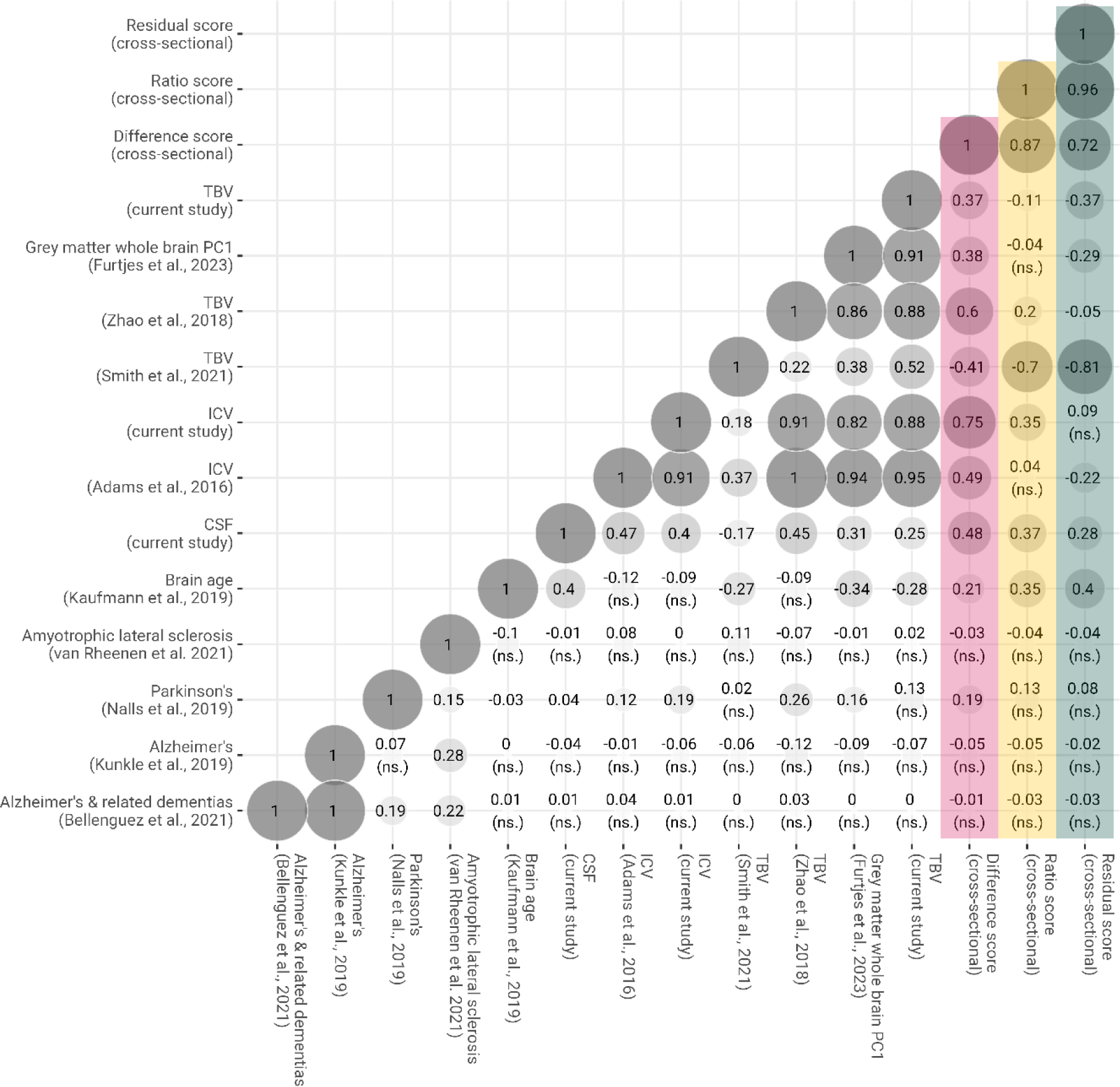
Genetic correlations inferred via LDSC between LBA calculated by three computational approaches and other structural neuroimaging and neurodegenerative phenotypes. Genetic correlations with the difference score are coloured pink, those with the ratio score are coloured yellow, and those with the residual score are coloured green. The notion ‘cross-sectional’ in the figure ‘indicates that this measure was calculated based on a single, cross-sectional MRI scan. To match procedures in phenotypic investigations, genetic correlates associated with LBA inferred with the ratio and residual method were flipped (i.e., multiplied by −1) whereby positive effect sizes indicate SNPs that increase the risk of greater LBA, which is the same direction of effects tagged by LBA inferred with the difference method. Unless marked with *ns.* genetic correlations were statistically significantly from zero at 99% confidence intervals (critical z score = 2.807).

The UKB data confirmed that LBA moderately captured longitudinal atrophic changes: Their correlations were numerically lower than in LBC1936, but still substantial considering atrophic changes in UKB only covered a 4-year time window (*N* = 4,674; *r* _residual_ = 0.29 [95% CI = 0.27-0.32], *r* _ratio_ = 0.24 [95% CI = 0.22-0.27], *r* _difference_ = 0.21 [95% CI = 0.18-0.24]; *SFig.5*). Given the shorter 4-year time window, mean atrophic changes were even more limited in UKB (mean *TBV _time 1_* at age 73 = 1,185 *mm^3^*, mean *TBV _time 2_* at age 82 = 1,171 *mm^3^*; *SFig.8-9*) than they were in LBC1936.

These correlations were very similar when using the T1-scaling factor instead of ICV to estimate LBA _residual_ (*SFig.6*). Correlations across all cross-sectional and longitudinal measures in LBC1936 and UKB are displayed in *SFig.4-5*, additionally illustrating that CSF volume does not correspond to LBA although CSF and LBA _difference_ may seem intuitively corollary because TBV equals ICV minus CSF volume. Our pre-registered efforts (https://osf.io/gydmw/) to artificially equate timelines for LBA and longitudinal atrophic changes were not safely interpretable due to the extremely healthy nature of UKB participant’s brains compared to the much younger cohorts MRi-Share and HCP (*SFig.1-2*; *Methods*).

Analyses across younger and older cohorts indicated that TBV and ICV were more strongly correlated (i.e., more similar measures) in the younger cohorts: *r* = 0.96 [95%CI 0.95-0.96] in MRi-Share [age: M (range) = 22 (18-35) years]; *r* = 0.92 [95%CI 0.90-0.93] in HCP [age: M (range) = 27 (22-31) years]), as compared to the older cohorts: *r* = 0.90 [95%CI 0.89-0.90] in UKB [age: M (range) = 62 (46-82) years]; *r* = 0.77 [95%CI 0.74-0.80] in Generation Scotland [age: M (range) = 62 (26-84) years]; *r* = 0.81 [95%CI 0.78-0.83] in LBC1936). Although confidence intervals overlap for estimates from different cohorts, this was consistent with our pre-registered expectations and is compatible with the assumption of our study that ICV is a measure of prior brain size whereby the brain approximately fills the entire intracranial vault in young adulthood – prior to the occurrence of any atrophy that may be detectable in older adulthood.

### Genome-wide association study of LBA

The second part of this paper explores the genetic bases of LBA by reporting its single nucleotide polymorphism (SNP)-heritability, and calculating its genome-wide associations in the UKB (*N* = 43,110) and genetic overlap with other structural neuroimaging and neurodegenerative traits.

#### SNP heritability

SNP-heritability was calculated using Genome-wide Complex Trait Analysis (GCTA; Yang et al., 2011) in European-only UKB genotype data with a cryptic relatedness cut-off at 0.025 (*N* = 38,624; *Table 1*). Although the SNP-heritability of the residual score was slightly lower (41% [95% CI = 38-43%]) than for either the ratio (42% [95% CI = 40-44%]) or difference score (47% [95% CI = 45-49%]), it is notable that the residual score was entirely independent of ICV – which has an even higher heritability of 58% [95% CI = 55-61%]. That is, even when the residual score was derived from the residuals of two highly correlated and highly heritable phenotypes (*Methods S2.2*), it carried substantial systematic genetic signal (*amplifier effect*). It supports our phenotypic observations that LBA _difference_ is a less specific, broader phenotype that it yielded larger heritability estimates (47% vs. 41%), more GWAS loci (37 vs. 28 loci), but fewer GWAS hits (3557 hits vs. 6016 hits) than LBA _residual_.

**Table 1.** Genetic characteristics of LBA.

| Traits | GCTA SNP-heritability (SE) | Mean $\chi^2$† | LDSC intercept (SE) | LDSC heritability (SE) | Significant* GWAS hits | Independent significant* GWAS hits | GWAS loci |
| --- | --- | --- | --- | --- | --- | --- | --- |
| **LBA (residual score)** | 0.41 (0.01) | 1.23 | 1.01 (0.008) | 0.26 (0.02) | 6016 | 93 | 28 |
| **LBA (ratio score)** | 0.42 (0.01) | 1.24 | 1.02 (0.009) | 0.27 (0.02) | 6371 | 112 | 30 |
| **LBA (difference score)** | 0.47 (0.01) | 1.27 | 1.02 (0.010) | 0.30 (0.02) | 3557 | 128 | 37 |
| **Cerebrospinal fluid (CSF)** | 0.39 (0.01) | 1.24 | 1.01 (0.009) | 0.27 (0.02) | 2812 | 48 | 21 |
| **Total brain volume (TBV)** | 0.56 (0.01) | 1.34 | 1.03 (0.011) | 0.36 (0.03) | 8576 | 158 | 56 |
| **Intracranial volume (ICV)** | 0.58 (0.01) | 1.35 | 1.03 (0.011) | 0.37 (0.03) | 7361 | 138 | 45 |

† Mean $\chi^2$ calculated in HapMap3 SNPs; \* Genome-wide significance threshold = $p < 5 \times 10^{-8}$ , independence determined at $r^2 < 0.6$ calculated from UKB release2b 10k European reference panel

#### GWAS associations

We performed European-only GWAS analysis in REGENIE (Mbatchou et al., 2021). FUMA v1.5.2 (Watanabe et al., 2017; default settings) identified 28 independent genomic risk loci associated with LBA _residual_ (*Fig.4*). The most significant SNP, *rs142005327* (*p* = 3.16 x10^-46^) was part of a locus spanning from 120777961 to 121033191 on chromosome 7. Gene prioritisation in FUMA using expression quantitative trait loci (*eQTL*) mapping [PsychENCODE (Wang et al., 2018), BRAINEAC (Ramasamy et al., 2014), GTEx v8 Brain (Aguet et al., 2020)] indicated that our most significant risk locus contained SNPs that are known *eQTLs* of the *WNT16* gene expressed in the brain. The *WNT* gene family encodes signalling proteins involved in oncogenesis and developmental processes including regulation of cell fate (Staal et al., 2008). According to the GWAS Catalog, SNP rs142005327 was previously linked with brain morphometry (van der Meer et al., 2021; van der Meer et al., 2022), brain age (Kim et al., 2023), but also bone mineral density (Greenbaum et al., 2022) and osteoporosis (Guindo-Martínez et al., 2021) which are both age-related diseases and were linked to *WNT* signalling. Both *APOE* SNPs rs7412 and rs429358 were not associated with either of the LBA phenotypes (LBA _residual_; *p _rs7412_* = 0.431; *p _rs429358_* = 0.735).

LBA _ratio_ identified 30 genomic risk loci, 23 of which overlapped with the 28 genomic loci identified by LBA _residual_ (*Table 1*). LBA _difference score_ identified 37 genomic risk loci, 16 of which overlapped with the 28 LBA _residual_ genomic loci. This suggests LBA _ratio_ and LBA _difference_ captured broader and less precise genetic signal compared to LBA _residual_. All three LBA phenotypes produced similar looking Manhattan plots (*SFig.15-16*). The top hit in LBA _ratio_ was rs142005327 and the top hit in LBA _difference_ was rs10668066, both located in chromosome 7. The nearest gene for both these SNPs was the *WNT16* gene. Manhattan plots for TBV, ICV, and CSF volume calculated in the same sample as the LBA phenotypes are in *SFig.17-19*. SNP-by-age interaction analyses indicated no evidence for significant SNP-by-age interactions on any of the LBA phenotypes (*p* < 5 x10^-8^; *SFig.20-32*), which included both *APOE* SNPs rs7412 and rs429358 (LBA _residual_: *p _rs7412_* = 0.195; *p _rs429358_* = 0.660; detailed outline in *SMaterials*).

#### Genetic correlations

To characterise LBA, we calculated genetic correlations using bivariate LDSC (Bulik-Sullivan et al., 2015) with different structural neuroimaging traits (*Fig.5*). LBA _residual_ had a weak negative genetic correlation with our TBV GWAS (*r_g residual_* = −0.37; *SE* = 0.05) and was uncorrelated with the TBV GWAS by Zhao et al. (2019) (*r_g residual_* = −0.05; *SE* = 0.05), both of which did *not* include ICV as a covariate. The same measure of LBA _residual_ was strongly associated with the TBV GWAS by Smith et al. (2021) (*r_g residual_* = −0.81; *SE* = 0.06) where the TBV-associated SNP effects were adjusted for a close proxy of ICV (i.e., T1 scaling factor) as a covariate. The large magnitude of this genetic correlation suggests that the ICV-adjustment in Smith et al. mirrors our definition of LBA _residual_, indicating that this T1 scaling factor-corrected phenotype connotes genetic correlates of brain change rather than total brain volume alone, which the name would suggest.

Some genetic correlations underline the desirable properties of the residual method compared to the difference or ratio method (*r_g_* across three LBA scores = 0.72-0.96). Only LBA _residual_ was non-significantly genetically correlated with ICV (*r_g_* = −0.09; *SE* = 0.05). The other computational methods produced substantial genetic correlations with ICV (*r_g_* _ratio_ = 0.35, *SE* = 0.05; *r_g_* _difference_ = 0.75; *SE* = 0.06), which confirms phenotypic observations that only the residual method captures variance un-contaminated by ICV baseline levels. Furthermore, the difference method appears to have induced the counter-intuitive genetic correlation between greater LBA and larger TBV (*r_g_* = 0.37; *SE* = 0.04); as opposed to the opposite direction of effects indicated by the ratio (*r_g_* = −0.11; *SE* = 0.04) and the residual method (*r_g_* = −0.37; *SE* = 0.05). We suggest a negative genetic correlation between LBA and TBV is theoretically more intuitive because a simple measure of TBV will likely capture, among other factors, both early-life neurodevelopment as well as later-life neurodegeneration (i.e., decline which is compatible with a negative direction of effects). A GWAS-by-subtraction model (Demange et al., 2021) confirmed that the genome-wide TBV-associated residuals of ICV (*Methods*) near perfectly genetically overlapped with LBA _residual_ (*r_g_* = 0.95 between LBA _residual_ and TBV residualised for ICV on a genome-wide level). Heritability in males (LBA _difference_ = 0.37 [0.03], LBA _ratio_ = 0.30 [0.03], LBA _residual_ = 0.27 [0.03]) was similar but numerically larger than heritability in females (LBA _difference_ = 0.27 [0.03], LBA _ratio_ = 0.24 [0.03], LBA _residual_ = 0.25 [0.03]). Genetic correlations across LBA phenotypes calculated in males (*n* = 20,453) and females (*n* = 22,657) separately were large (*r_g_* = 0.91-1.00; *SFig.34*) indicating that genetic signal in the GWAS including all participants was largely unrelated to sex.

We also calculated genetic correlations between LBA and an array of neurogenerative diseases for which we identified well-powered GWAS summary statistics. Alzheimer disease (Kunkle et al., 2019) and related dementias (Bellenguez et al., 2022), and amyotrophic lateral sclerosis (ALS; van Rheenen et al., 2021) were genetically uncorrelated with LBA (*Fig.5*). Parkinson disease (Nalls et al., 2019) yielded significant genetic correlations with estimates of LBA _difference & ratio_, but not LBA _residual_. Because the difference and ratio scores substantially captured ICV-associated genetic variance (*r_g_* _LBA ratio & ICV_ = 0.35, *SE* = 0.04; *r_g_* _LBA difference & ICV_ = 0.75, *SE* = 0.05) and the residual score did not (*r_g_* _LBA residual & ICV_ = −0.09, *SE* = 0.05; *ns*.), we suggest that the significant associations between Parkinson disease and the difference and ratio scores were likely driven by ICV baseline differences (*r_g_* _Parkinson disease & ICV_ = 0.19; *SE* = 0.05) rather than brain changes (Krabbe et al., 2005).

Our analyses do not include a genetic correlation with longitudinal atrophic changes from Brouwer et al. (2022) – the largest multi-cohort GWAS of longitudinal lifespan brain changes (*N* ∼ 16,000) – because this set of GWAS summary statistics produced a negative LDSC heritability estimate. This means we were unable to detect systematic polygenic signal across a subset of HapMap3 SNPs, despite this trait having been reported to carry small but extant genetic signal (LDSC *h^2^*Z statistic = 1.24 reported in Brouwer et al., 2022). Unlike LBA, Brouwer et al. indicated two top SNPs in chromosomes 13 and 16, and identified no correlates in chromosome 7, or any SNP-correlates associated with *WNT16* signalling.

## Discussion

### Phenotypic characteristics of LBA

It was the aim of this study to externally validate and explicitly characterise a cross-sectional measure of LBA using three computational modelling techniques [difference (*ICV-TBV*), ratio (*TBV*/*ICV*), and regression-residual method (TBV∼ICV)], which were then carried forward to conduct the largest GWAS of LBA to date (*N* = 43,110). Our phenotypic investigations support that this straightforward definition of LBA (derived from a single MRI scan) yielded meaningful variance that sensibly correlated with brain atrophy estimated from neuroradiological rating scales, cognitive functioning, and frailty. It was also moderated by sample age, and LBA moderately captured longitudinal atrophic changes measured over a short time period (<9years). Given the much greater availability of samples with a single MRI scan and genetic information, LBA represents a cost-effective GWAS phenotype to boost statistical power for discovery of the molecular genetic bases of lifetime changes in brain atrophy. We demonstrated that these phenotypic properties were maximally captured when LBA was estimated with the residual rather than the difference or ratio method. Estimating LBA _residual_ exploited an amplifier effect whereby it remained statistically independent of neurodevelopmental baseline differences in head size (i.e., uncorrelated with ICV), at the same time as maximally focusing our analyses on age-related neurodegenerative brain volume *changes*.

### Summary of LBA genetic correlates

Building on this carefully described measure of LBA, we performed genome-wide investigations in over 2.5-times more participants (*N* = 43,110) than had contributed to previous efforts based on meticulously collected longitudinal MRI scans (*N* ∼ 16,000; Brouwer et al., 2022). LBA was substantially heritable (*h^2^*_SNP GCTA_ = 41% [95% CI = 38-43%]), and our GWAS summary statistics yielded strong polygenic signal in the LDSC framework (*h^2^*Z statistic = 13). These characteristics qualify our GWAS summary statistics to form a useful contribution to future investigations of neurodegenerative diseases (e.g., using polygenic scores, or Genomic Structural Equation Models; Grotzinger et al., 2019).

The most significant genomic risk locus of LBA indicated the *WNT16* gene in chromosome 7 which has previously been linked to reduced cortical volume in Alzheimer patients (Dong et al., 2023) and has also been implicated in other age-related traits like brain age (Kim et al., 2023) and osteoporosis (Guindo-Martínez et al., 2021). *WNT* signalling has been linked to neurodegenerative diseases such as Parkinson disease and ALS (Liu et al., 2022), as well as amyloid-beta-related pathogenesis in Alzheimer disease where *WNT* was suggested to drive synaptic loss (Elliott et al., 2018).

LBA did not demonstrate significant genetic correlations with existing GWAS of neurodegenerative diseases such as Alzheimer and Parkinson disease (or produced significant genetic correlations we suggest are driven by ICV), but their genetic correlation with LBA may be subject to insufficiently powered neurodegenerative sample sizes. Future, more detailed genetic follow-up analyses (for example, performing gene prioritisation or Mendelian Randomization) should be conducted in larger, better-powered, and multi-ancestral investigations. This paper is a pre-cursor for a larger consortium-level meta-analytic GWAS of this single-time-point definition of LBA which promises much larger sample sizes.

### Significance of carefully considered methodological approaches

The work presented here highlights the importance of carefully considering and interpreting different methodological approaches used to estimate lifetime changes (i.e., difference, ratio, residual method) which includes adjusting for the appropriate covariates in the GWAS analysis. Only change calculated with the residual method will, by definition, be uncorrelated with ICV baseline differences (*r_g_* = |0.09| *ns.*), and change calculated using the difference or ratio method highly likely comprises results driven by ICV baseline differences. Hence, the difference and ratio methods tag genetic signal that amalgamates late-life neurodegeneration (or any longitudinal changes), as well as developmental processes indicated by ICV (*r_g_* between ratio score and ICV = |0.35|; *r_g_* between difference score and ICV = |0.75|).

This consideration also applies to interpreting GWAS results from previous studies of TBV, where adjusting for ICV re-directed the tagged variance away from broad differences in brain size, towards lifetime neurodegenerative changes more specifically. For example, consider the *big40* TBV GWAS by Smith et al. (2021) who followed the well-established practice of adjusting GWAS analyses for a close proxy variable of ICV (i.e., T1 scaling factor). This ICV-adjusted TBV GWAS was strongly genetically correlated with LBA _residual_ which we postulate should capture change (*r_g_* = |0.81|), and had much weaker genetic correlations with other TBV GWAS that were unadjusted for ICV (*r_g_* = |0.22-0.52|) (Fürtjes et al., 2023; Zhao et al., 2019; and current study). By contrast, the unadjusted TBV GWAS yielded much smaller genetic correlations with LBA _residual_ (*r_g_* = |0.05-0.37|) because unadjusted TBV tags a broader and less specific phenotype underpinned by both early-life neurodevelopment and late-life neurodegeneration. This reasoning only holds in later-life cohorts, and does not extend to young, pre-neurodegenerative cohorts where TBV is much more similar to ICV. LBA would hence not index neurodegeneration but instead healthy variability in earlier life development which is not the focus of this study. Our work highlights the requirement for transparent characterisations, not only of the GWAS phenotype, but also of the residuals that remain after adjusting for covariates.

### Limitations of a broad LBA phenotype

We focused on LBA across the whole brain – rather than individual regions or tissue types – because precision and accuracy of a volume estimated in MRI is maximal, the larger the volume (Azimbagirad et al., 2021). The disadvantage to maximising measurement reliability is that LBA is a very broad phenotype biologically underpinned by a collection of many different tissue types and biological mechanisms. For example, it is unclear to what extent LBA captures cellular disintegration within brain regions (i.e., grey matter), or between brain regions (i.e., white matter). Gross morphometry from structural MRI does not indicate whether neurons, microglia, astrocytes, or other cell types are affected, or whether neurodegeneration resulted from ischemic damage, though this applies to regional as well as global MRI measurements. It would also not allow the detection of specific signatures of frontal and anterior temporal atrophy observed in fronto-temporal dementia (Young et al., 2018), or other regional neurodegenerative patterns in other diseases. Future studies are required to develop reliable ways to record those more specific and fine-grained changes and how they may relate to the genetics of brain-wide atrophy analysed here.

### Conclusion

This study provided a comprehensive phenotypic characterisation of LBA estimated from a single MRI scan, using five cohorts across adulthood, which allowed boosting statistical power for better genetic discovery. LBA was strongly heritable, had strong polygenic signal, and the strongest genetic correlates highlighted the *WNT16* pathway as one potential mechanism underpinning broad, late-life neurodegeneration. Our work underlined the importance of using appropriate computational approaches, where we demonstrated that only the residual method can focus analyses of TBV towards late-life neurodegeneration independent of earlier-life neurodevelopmental processes. Finally, we showed that it is imperative for neuroimaging genetics studies to transparently characterise not only their GWAS phenotype, but also the remaining residuals after covariate adjustment because the inclusion of ICV (or a close proxy variable such as the T1 scaling factor) as a covariate has important implications for the interpretation of phenotypic and genetic results.

## Reproducibility statement

Analyses presented here were pre-registered at https://osf.io/gydmw/. The analysis only relied on open-source software and the code is displayed at https://annafurtjes.github.io/BrainAtrophy_Genetics/.

## Acknowledgements

We thank the participants who took part in LBC1936, UK Biobank, Human Connectome Project, MRi-Share and Generation Scotland, and the team members and radiographers who collected, entered, processed and disseminated the data used in this paper.LBC1936 MRI acquisition and initial analyses were conducted at the Brain Research Imaging Centre, Neuroimaging Sciences, University of Edinburgh (www.bric.ed.ac.uk), which is part of the SINAPSE (Scottish Imaging Network—A Platform for Scientific Excellence) collaboration (www.sinapse.ac.uk), funded by the Scottish Funding Council and the Chief Scientist Office. The LBC1936 is supported by the Biotechnology and Biological Sciences Research Council, and the Economic and Social Research Council [BB/W008793/1], Age UK (Disconnected Mind project), the Medical Research Council [G0701120, G1001245, MR/M013111/1, MR/R024065/1], the Milton Damerel Trust, and the University of Edinburgh. We acknowledge the work of the LBC1936 radiographers, particularly Gayle Barclay, Donna MacIntyre, Iona Hamilton, Charlotte Jardine who have enabled the high retention of the MRI component. The 1.5T research MRI was funded by the Scottish Funding Council. We acknowledge the work of the LBC1936 radiographers Gayle Barclay, Donna MacIntyre, Iona Hamilton, Charlotte Jardine, who have enabled the high retention of the MRI component. We also thank Professor Stephen M. Smith for discussion on UK Biobank inter-site scanning differences. This research was conducted, using the UK Biobank Resource under approved project 10279. HCP data were provided by the Human Connectome Project, WU-Minn Consortium (Principal Investigators: David Van Essen and Kamil Ugurbil; 1U54MH091657) funded by the 16 NIH Institutes and Centers that support the NIH Blueprint for Neuroscience Research; and by the McDonnell Center for Systems Neuroscience at Washington University. Generation Scotland received core support from the Chief Scientist Office of the Scottish Government Health Directorates (CZD/16/6) and the Scottish Funding Council (HR03006) and is currently supported by the Wellcome Trust [216767/Z/19/Z]. This study was also supported and funded by the Wellcome Trust Strategic Award ‘Stratifying Resilience and Depression Longitudinally’ (STRADL) (Reference 525 104036/Z/14/Z).

AF, IF & GD are supported by National Institutes of Health (NIH) grant [R01AG073593]. SRC is supported by a Sir Henry Dale Fellowship jointly funded by the Wellcome Trust and the Royal Society (221890/Z/20/Z). EMTD was additionally supported by R01MH120219. EMTD is member of the Population Research Center (PRC) and the Center on Aging and Population Sciences (CAPS) at The University of Texas at Austin, which are supported by NIH grants P2CHD042849 and P30AG066614, respectively. W.D.H., and C.X. are supported by a Career Development Award from the Medical Research Council (MRC) [MR/T030852/1] for the project titled “From genetic sequence to phenotypic consequence: genetic and environmental links between cognitive ability, socioeconomic position, and health”. JW is funded by the UK Dementia Research Institute which receives its funding from DRI Ltd, funded by the UK Medical Research Council, Alzheimer’s Society and Alzheimer’s Research UK. ADG is supported by NIH Grants R01MH120219 and R01AG073593. EB & KF are funded by the Stroke Association/BHF/Alzheimer’s Society ‘Rates Risks and Routes to Reduce Vascular Dementia’ (R4VaD) Priority Programme Award in Vascular Dementia (16 VAD 07) and the Row Fogo Centre for Research into Ageing and the Brain (Ref No: AD.ROW4.35. BRO-D.FID3668413).

## Footnotes

1 Note that here we focus on brain atrophy in older participants because their brains tend to have experienced some degree of brain shrinkage as compared to their brains’ prior size as indicated by ICV. Modelling change by considering TBV relative to ICV in older participants is conceptually distinct from performing ICV correction in young/ pre-neurodegenerative participants where age-related atrophy is *not* the target outcome. As we cannot reasonably expect brain atrophy in young participants where TBV will be equal to (or very similar to) ICV, ICV-correction will not capture brain shrinkage but instead normal healthy biological variability with roots in development (e.g., Dhalama et al., 2022). By contrast, when ICV correction is performed in populations with measurable brain atrophy (e.g., due to old age or pathology), it focuses the analysis on brain matter *shrinkage* because TBV is assessed relative to baseline head size which can help amplify associations with ageing-related traits (e.g., Lyall et al., 2013) as is also our objective here.

2 Note that some of the ageing-related measures in UKB used here were recorded prior to the MRI scan. We analyse all these measures as outcome variables to be predicted by LBA. Results describe their relationship, but we do not imply any directionality or causality.

