## SupplementaryPlots for "Lifetime brain atrophy estimated from a single MRI: measurement characteristics and genome-wide correlates"

**Supplementary plots**

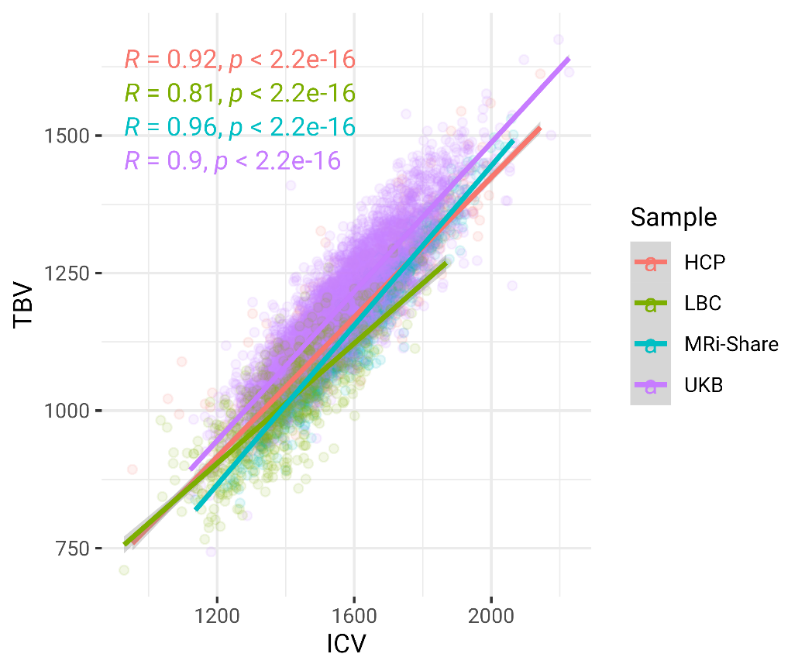

*SFig.1.* Relationship between TBV and ICV in all four considered cohorts. All three cohorts produced widely differing intercept and slope estimates: HCP (intercept = 161.77, slope = 0.63), Share (intercept = -4.42, slope = 0.72), UKB (intercept = 132.56, slope = 0.68), LBC (intercept = 249.58, slope = 0.54)

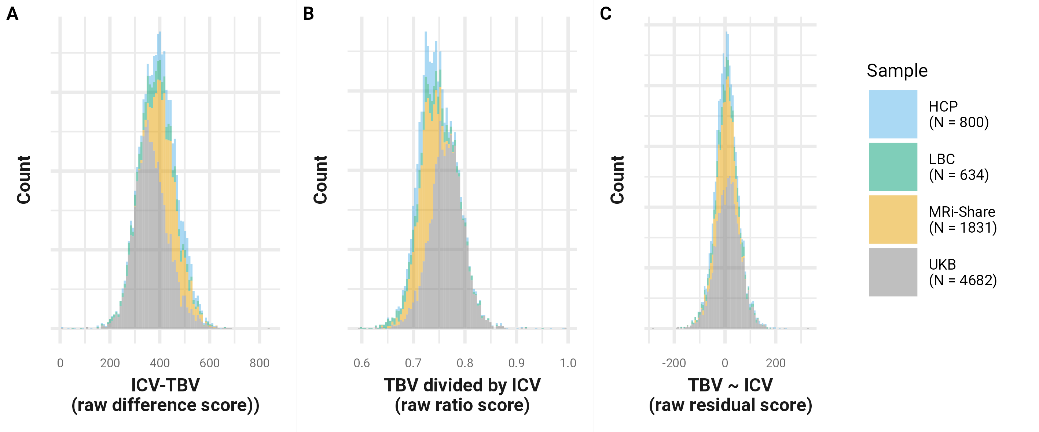

*SFig.2.* Distribution of lifetime atrophy scores in all considered samples. Regardless of their average sample age, those distributions look very similar, misleadingly indicating that brains scanned in the LBC and UKB cohorts appear just as healthy as those in the MRi-Share and HCP cohorts.

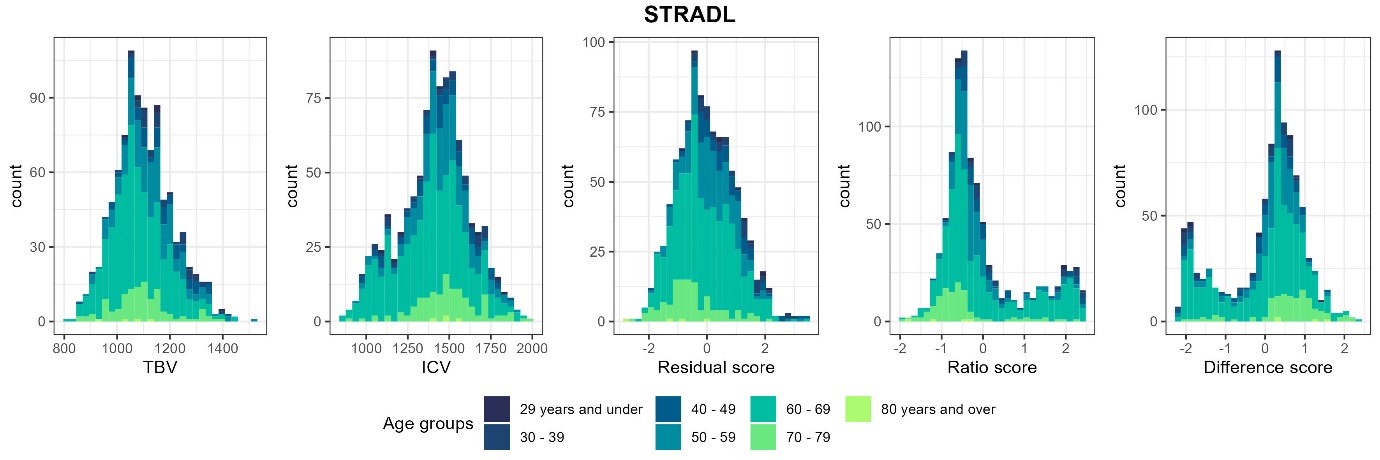

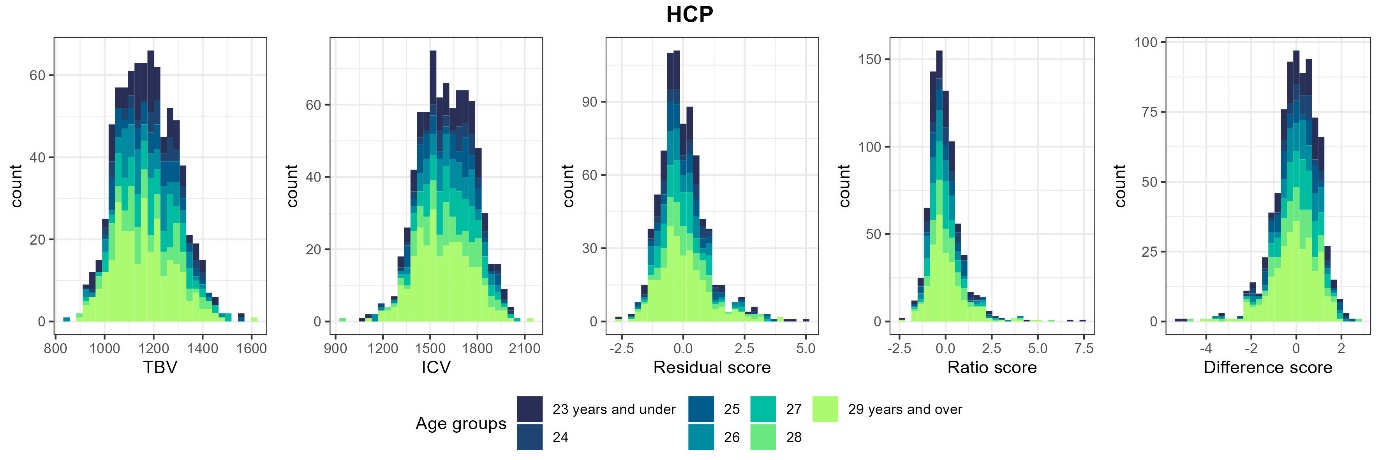

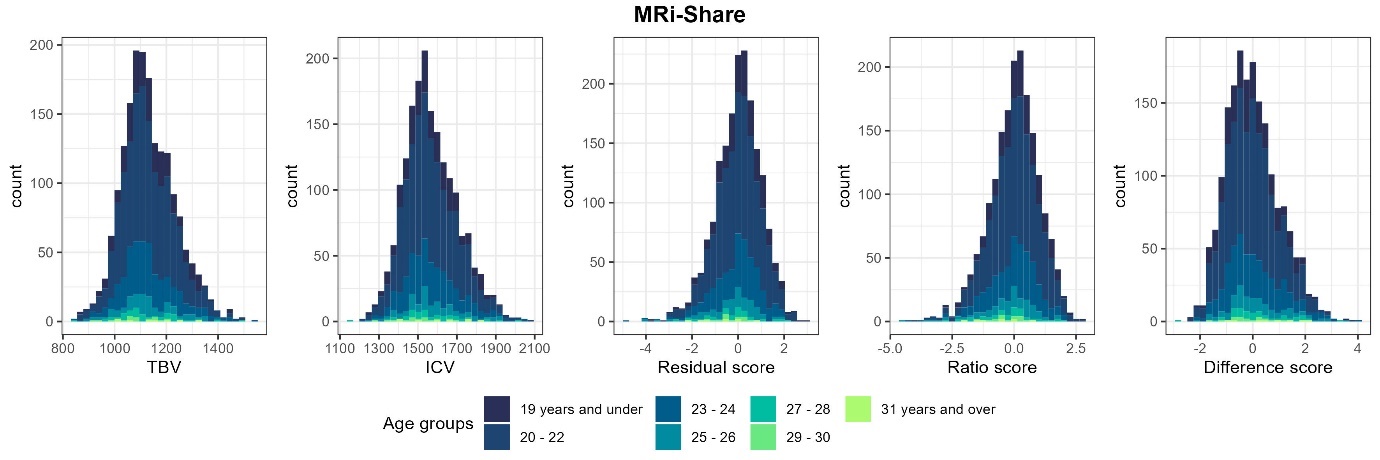

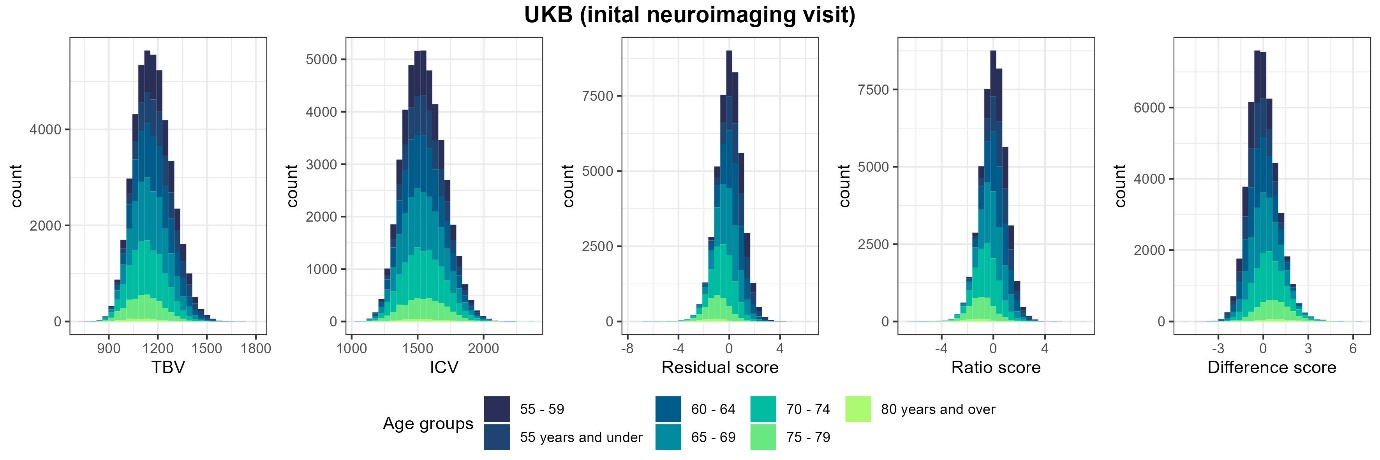

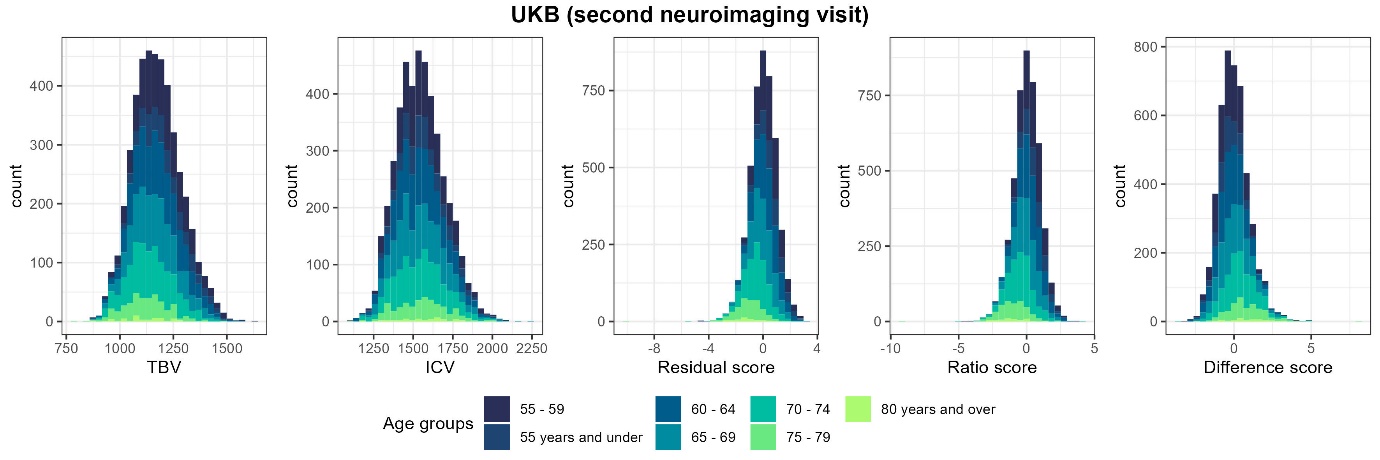

*SFig.3.* Distributions of TBV, ICV, and lifetime brain atrophy estimated with the residual, ratio, and difference method. Histograms are coloured by age groups.

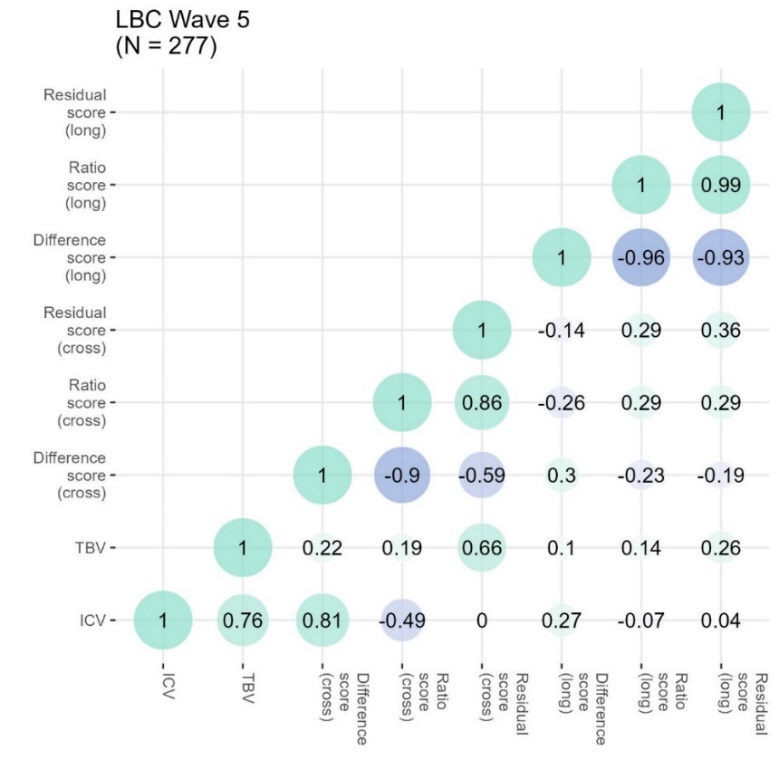

*SFig.4.* Pearson’s correlations in the LBC1936 cohort (wave 5) among TBV, ICV, CSF, lifetime atrophy scores inferred with three computational methods (‘cross’), and longitudinally-observed atrophic changes inferred with three computational methods (‘long’)

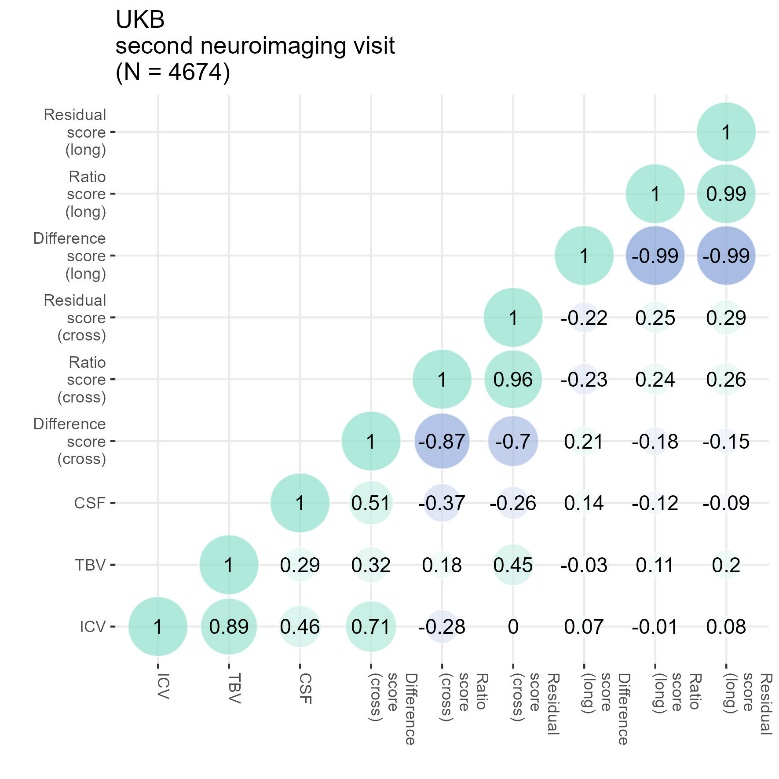

*SFig.5.* Pearson’s correlations in the UKB cohort among TBV, ICV, CSF, lifetime atrophy scores inferred with three computational methods (‘cross’), and longitudinally-observed atrophic changes inferred with three computational methods (‘long’).

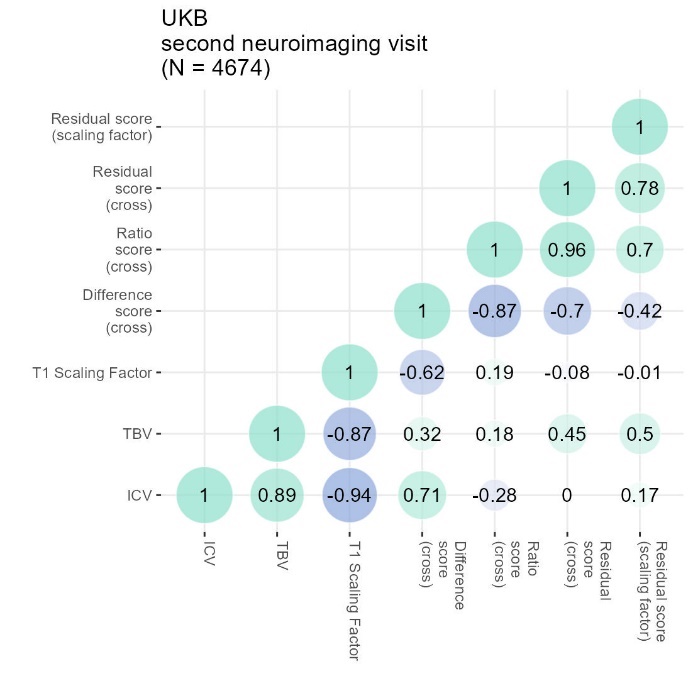

*SFig.6.* Pearson’s correlations in the UKB cohort among TBV, ICV, T1 Scaling Factor, and lifetime atrophy scores inferred with three computational methods (‘cross’), as well as lifetime atrophy inferred with a residual score using the T1 scaling factor instead of ICV (‘scaling factor’)

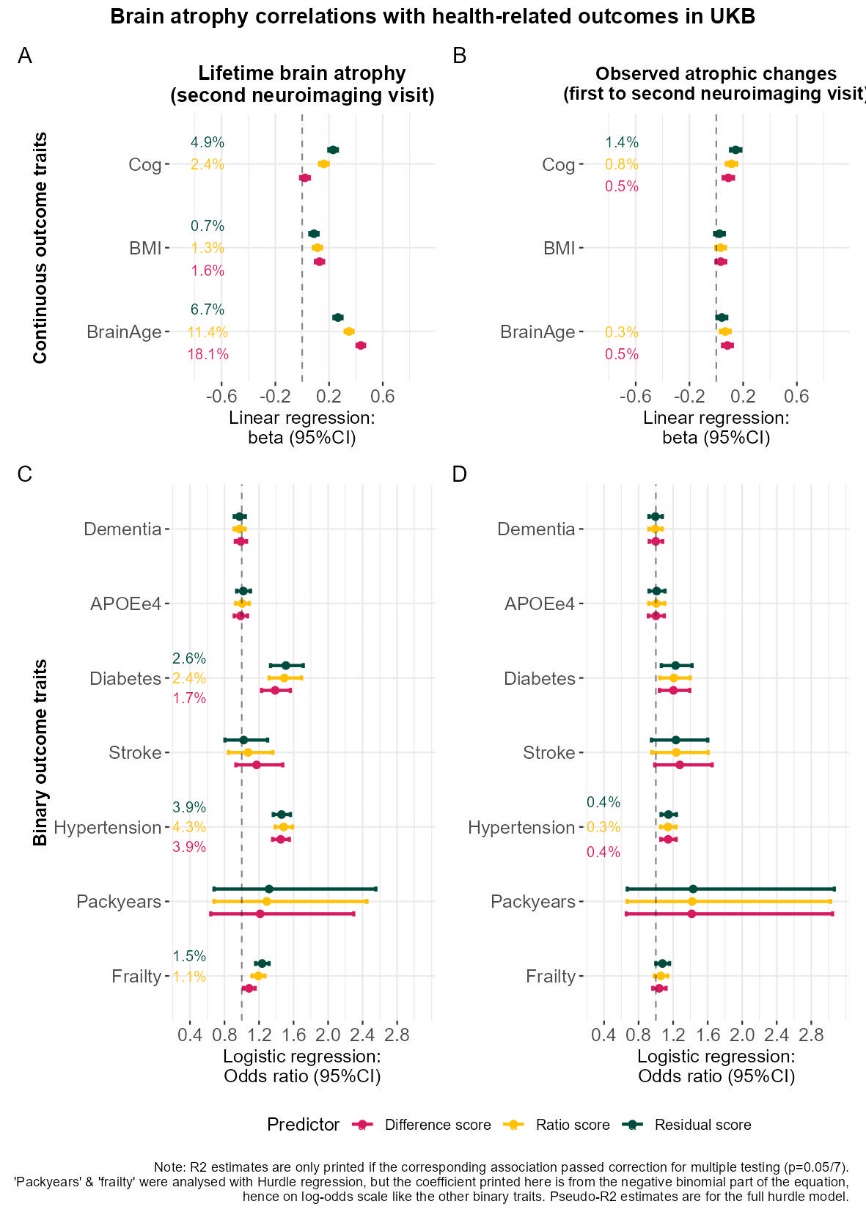

*SFig.7.* Associations with health-related phenotypes in UKB for ‘estimated’ LBA and longitudinally-‘observed’ atrophic changes. Atrophy estimated with the ratio and residual method were flipped to match the difference score whereby larger values represent more brain atrophy. Note that different variables were extracted for the UKB sample in a way that maximised information across available time points, which means that the MRI scan and the ageing-related information is not necessarily assessed at the same time (see Supplementary Methods).

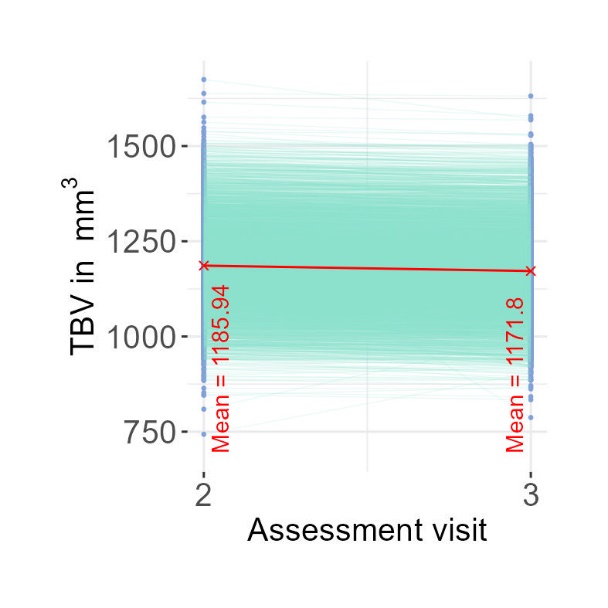

*SFig.8.* TBV estimates (*mm^3^*) in UKB participants with two MRI scans; *N* = 4682 at initial and repeated neuroimaging visit, mean lag = 4 years, age range at baseline = 46-81 years.

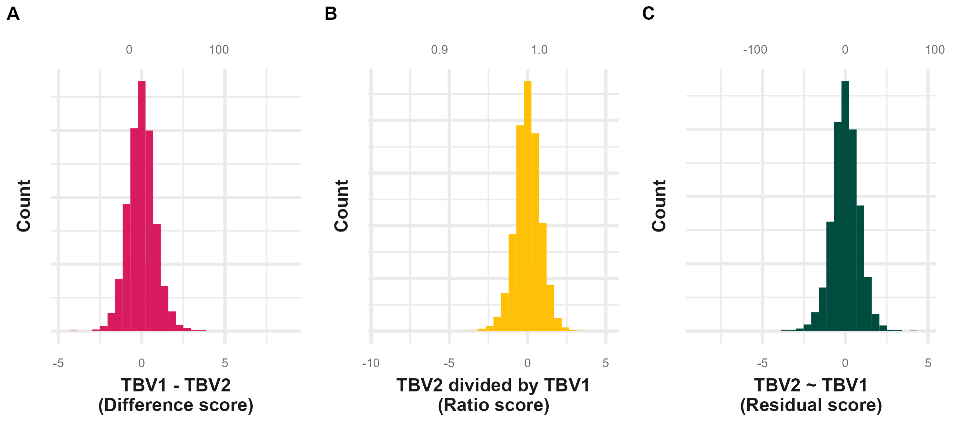

*SFig.9.* Distribution of atrophy scores derived from repeated TBV estimates in UKB (*N* = 4674). LBA _difference score_: minimum = -4.6, maximum = 8.55; LBA _ratio score_: minimum = -9.03, maximum = 5.02; LBA _residual score_: minimum = -8.72, maximum = 4.56.

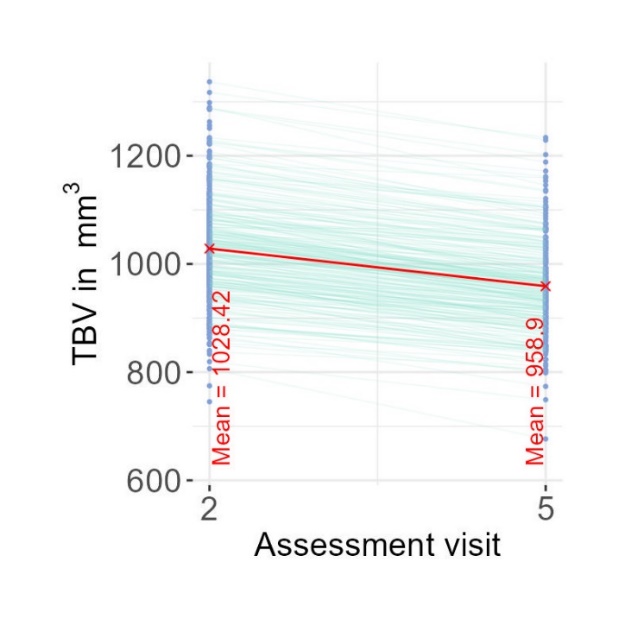

*SFig.10.* TBV estimates (*mm^3^*) in LBC1936 (*N* = 286) at first scan (wave 2) and fourth scan (wave 5), mean lag = 9 years, age range at baseline = 71-74 years.

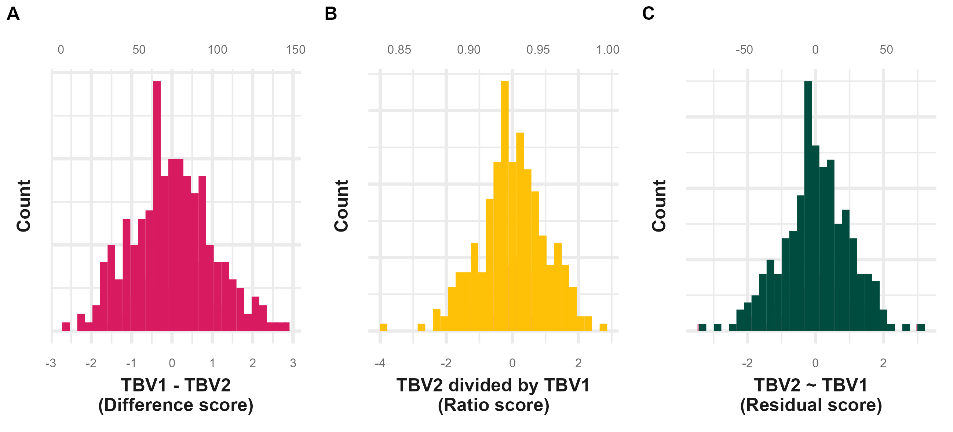

*SFig.11.* Distribution of atrophy scores derived from repeated TBV estimates in LBC1936 (*N* = 4674). LBA _difference score_: minimum = -2.63, maximum = 2.81; LBA _ratio score_: minimum = -3.88, maximum = 2.77; LBA _residual score_: minimum = -3.43, maximum = 3.00.

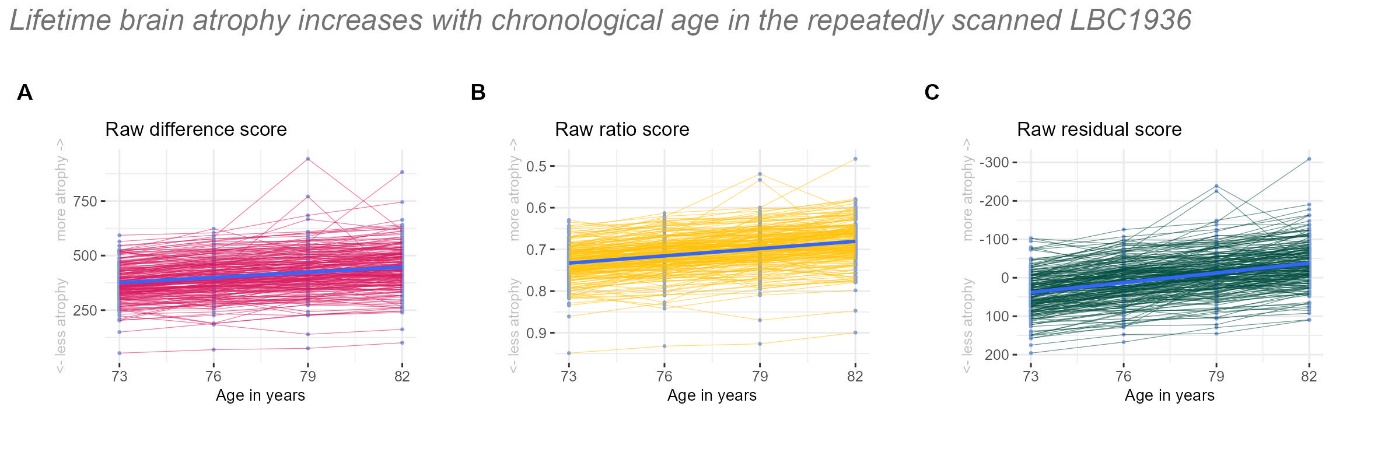

*SFig.12.* Repeated MRI measures age-correlation. LBA inferred at four assessments in the LBC1936. LBA was inferred at each time point from TBV and ICV estimates processed with the cross-sectional FS stream, which was meant to illustrate the tendency of LBA to increase with age, even when ICV is not specifically held constant across time points (which is what the FS longitudinal processing stream would have achieved). This figure displays only participants with estimates available across all waves. In this figure the residual score was calculated based on measurements at 4 time points so that atrophy estimates across all time points were derived relative to the same average value (this was required to illustrate time-dependent increases). In contrast, all other analyses across the manuscript derived the residual score based on one individual visit with only one entry for each participant. This figure displays raw LBA scores where ratio and residual scores were *not* flipped (i.e., multiplied with -1), but their *y*-axes are reversed. The scores were, however, flipped in all other figures and analyses to represent more LBA with larger values, and less LBA with smaller values.

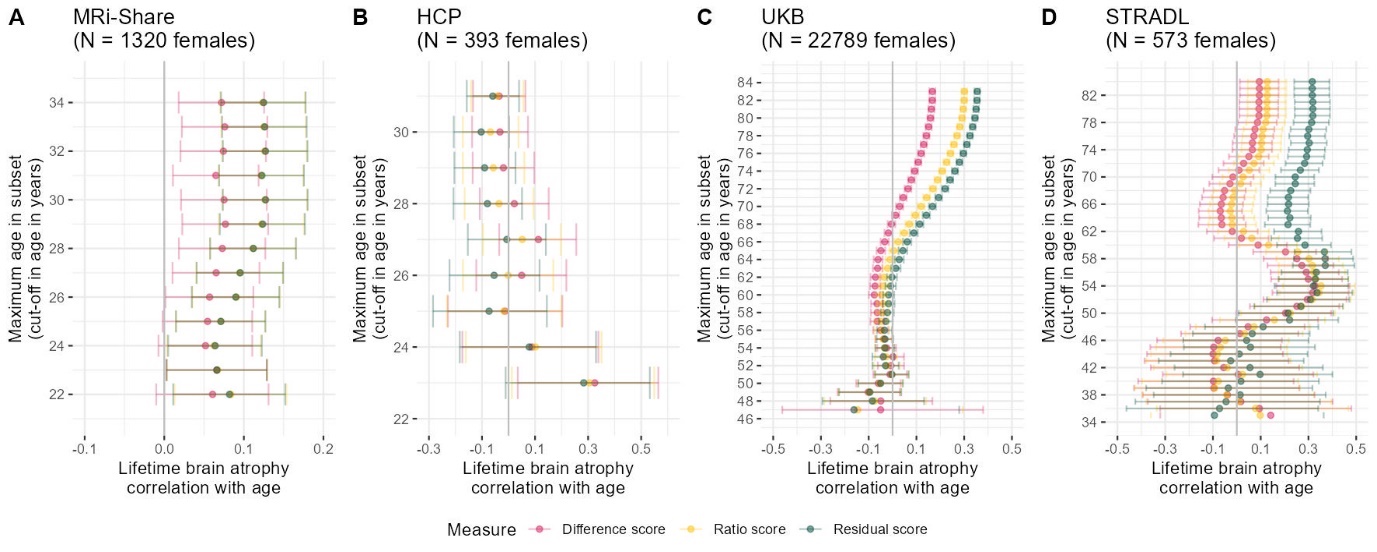

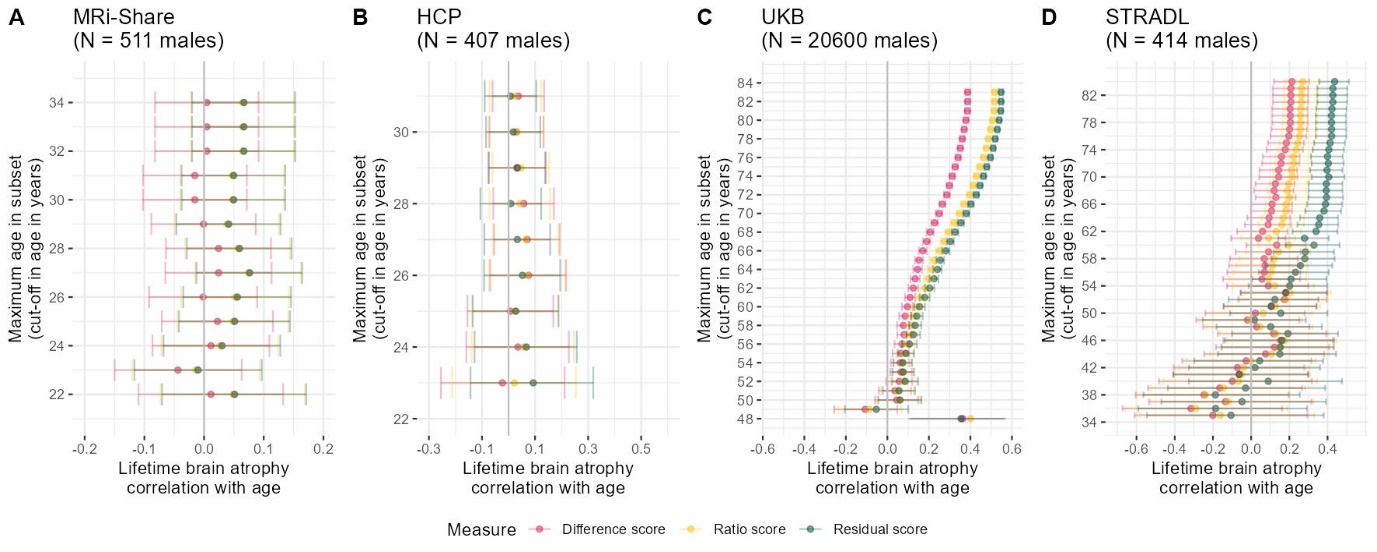

*SFig.13.* LBA is moderated by sample age across four earlier- and later-life cohorts. Re-analysis of data presented in Fig.3 split into males and females separately

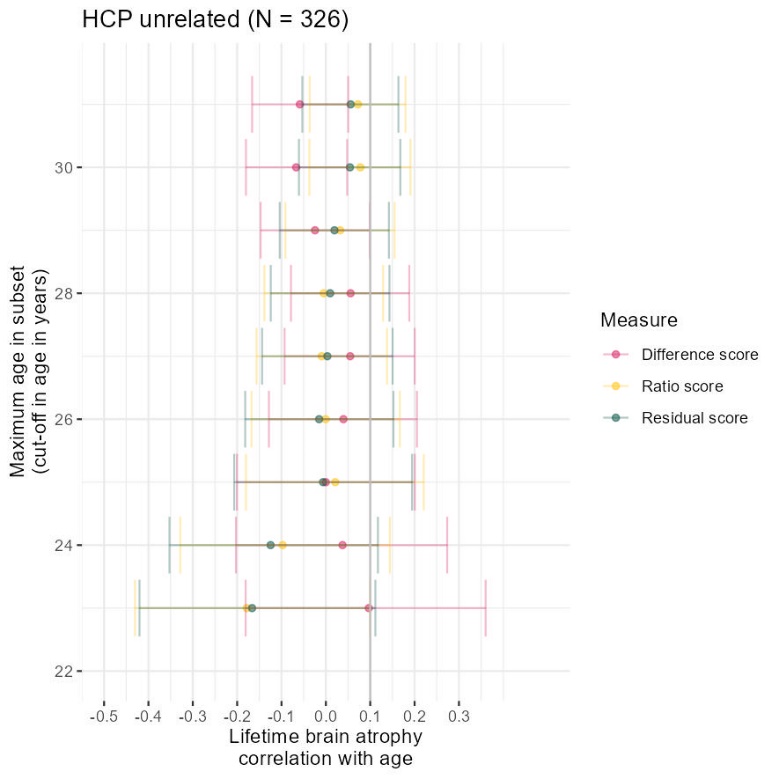

*SFig.14.* Age correlations in unrelated HCP sample (same as *Fig.2*, only in reduced unrelated sample)

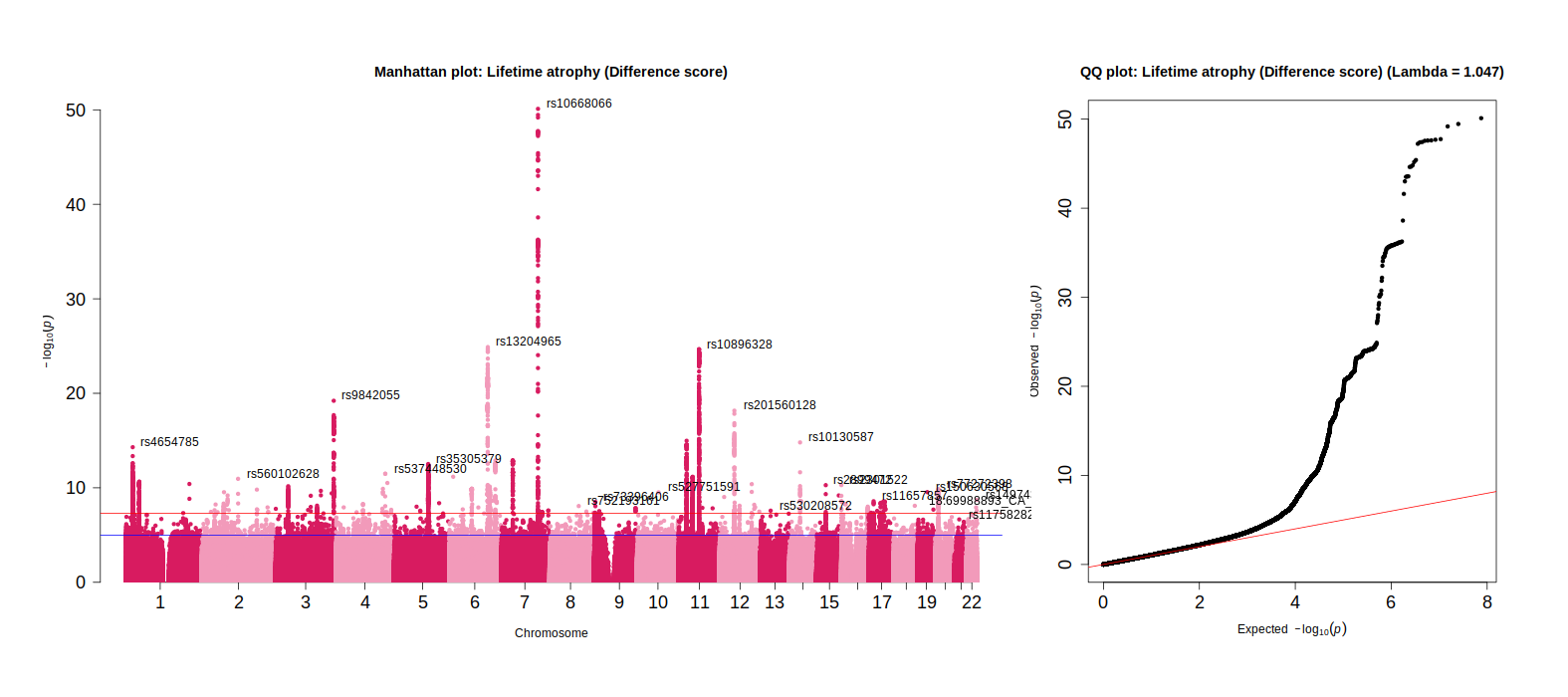

*SFig.15.* Manhattan plot for lifetime brain atrophy inferred with the difference method indicating top GWAS hits

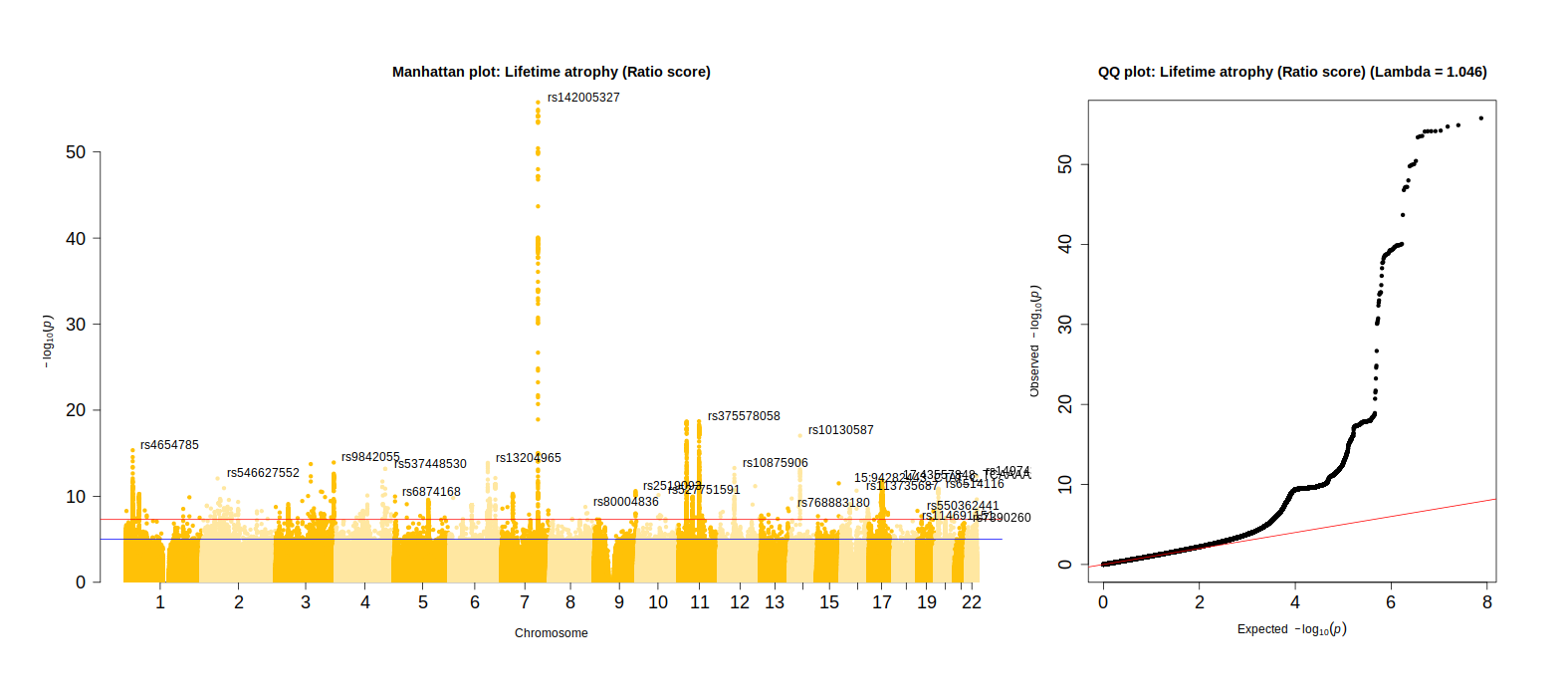

*SFig.16.* Manhattan plot for lifetime brain atrophy inferred with the ratio method indicating top GWAS hits

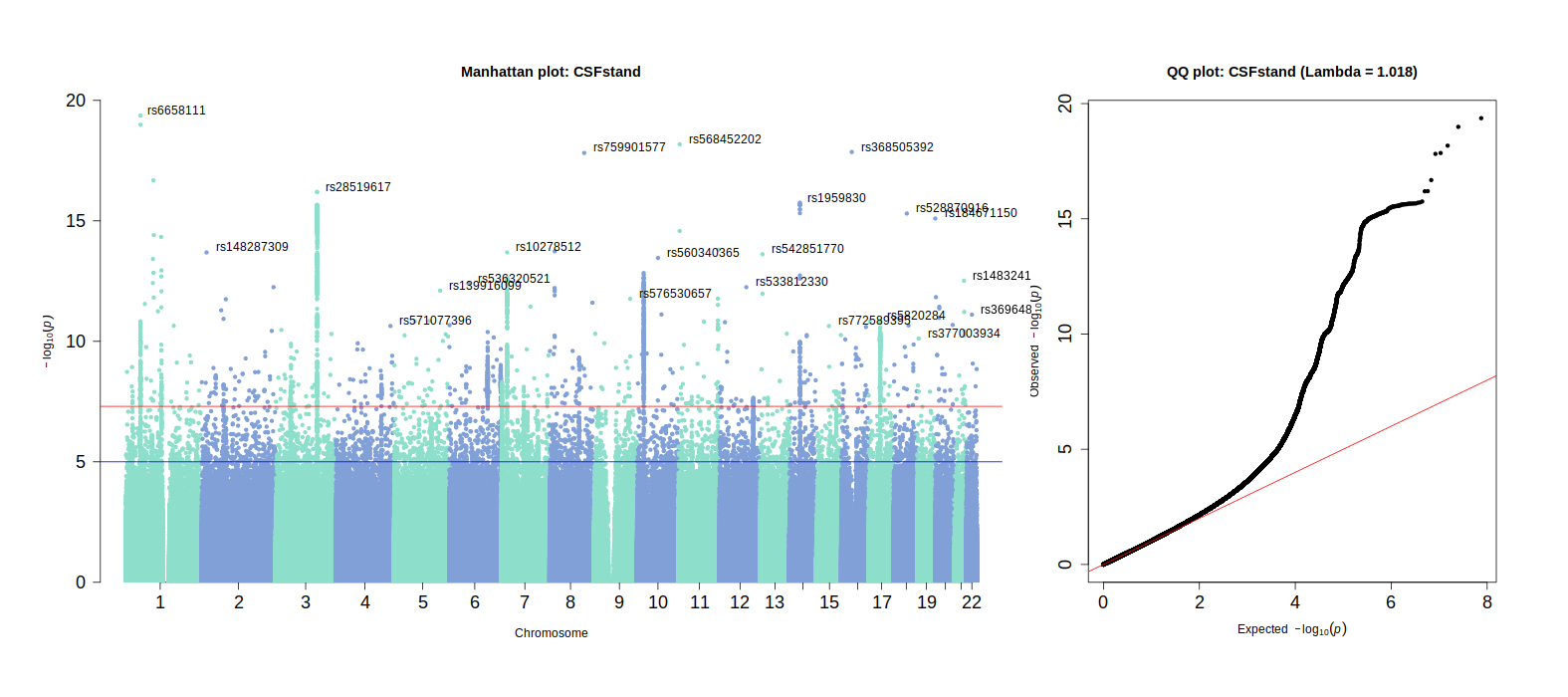

*SFig.17.* Manhattan plot for cerebrospinal fluid (CSF) volume indicating top GWAS hits

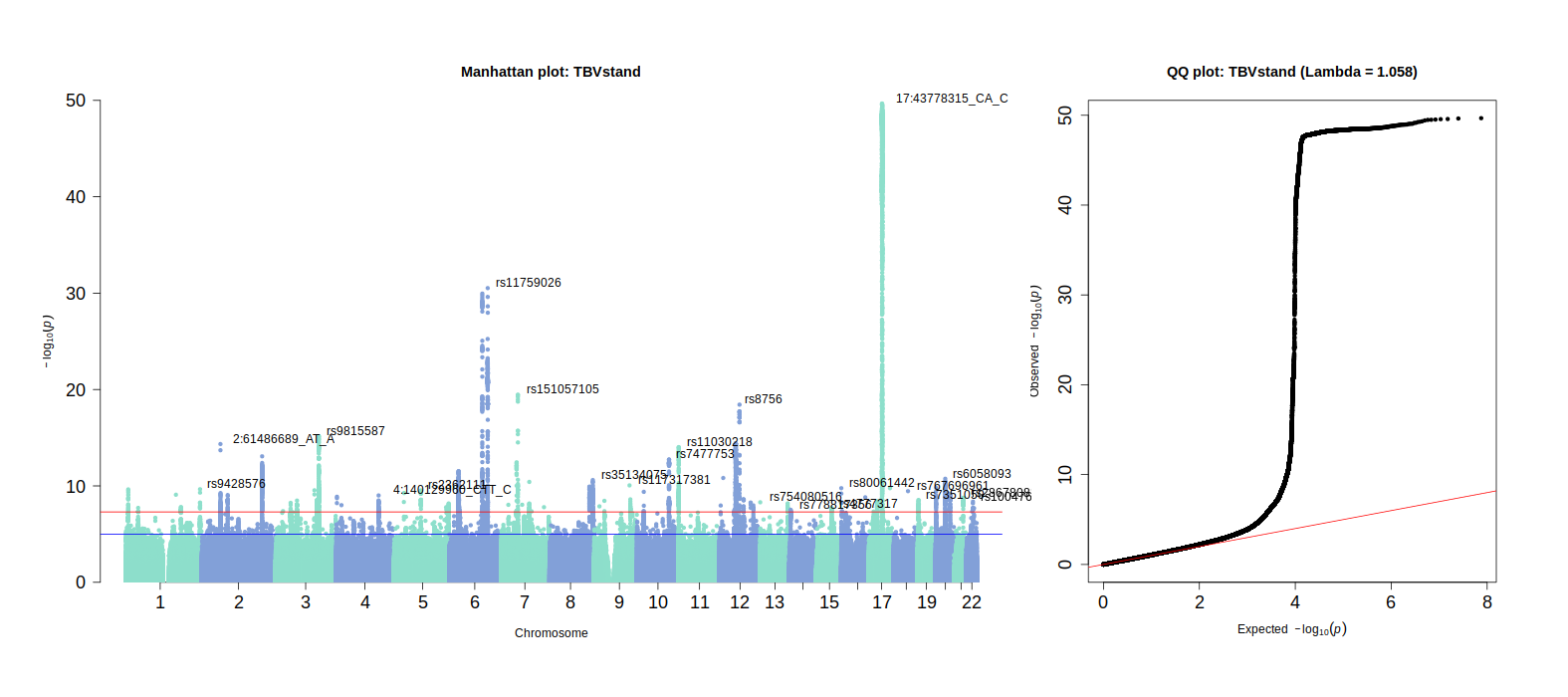

*SFig.18.* Manhattan plot for total brain volume (TBV) indicating top GWAS hits

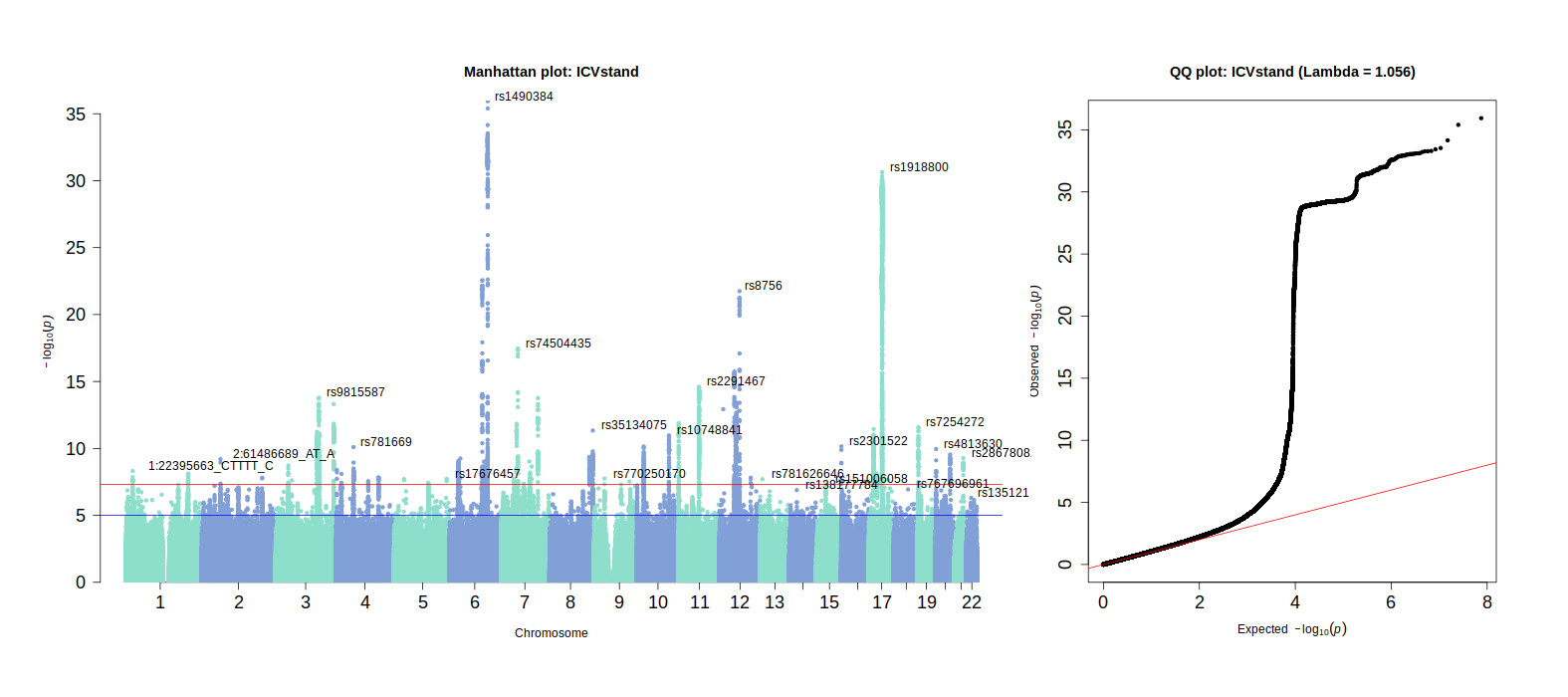

*SFig.19.* Manhattan plot for intracranial volume (ICV) indicating top GWAS hits

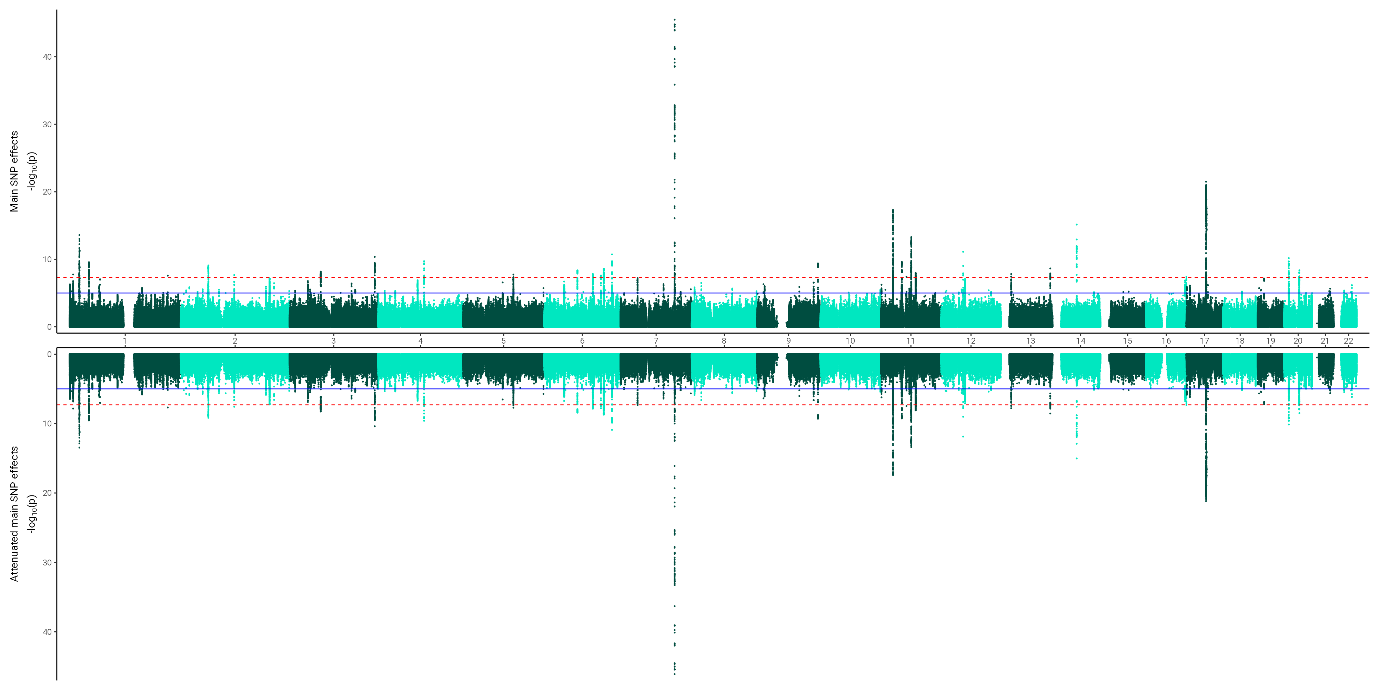

*SFig.20.* Miami plots for **residual score** contrasting main SNP effect *p*-values from original GWAS with main SNP effect *p*-values from GWAS analyses including an age interaction term (SNP filters: MAF > 0.01, INFO > 0.9, diallelic)

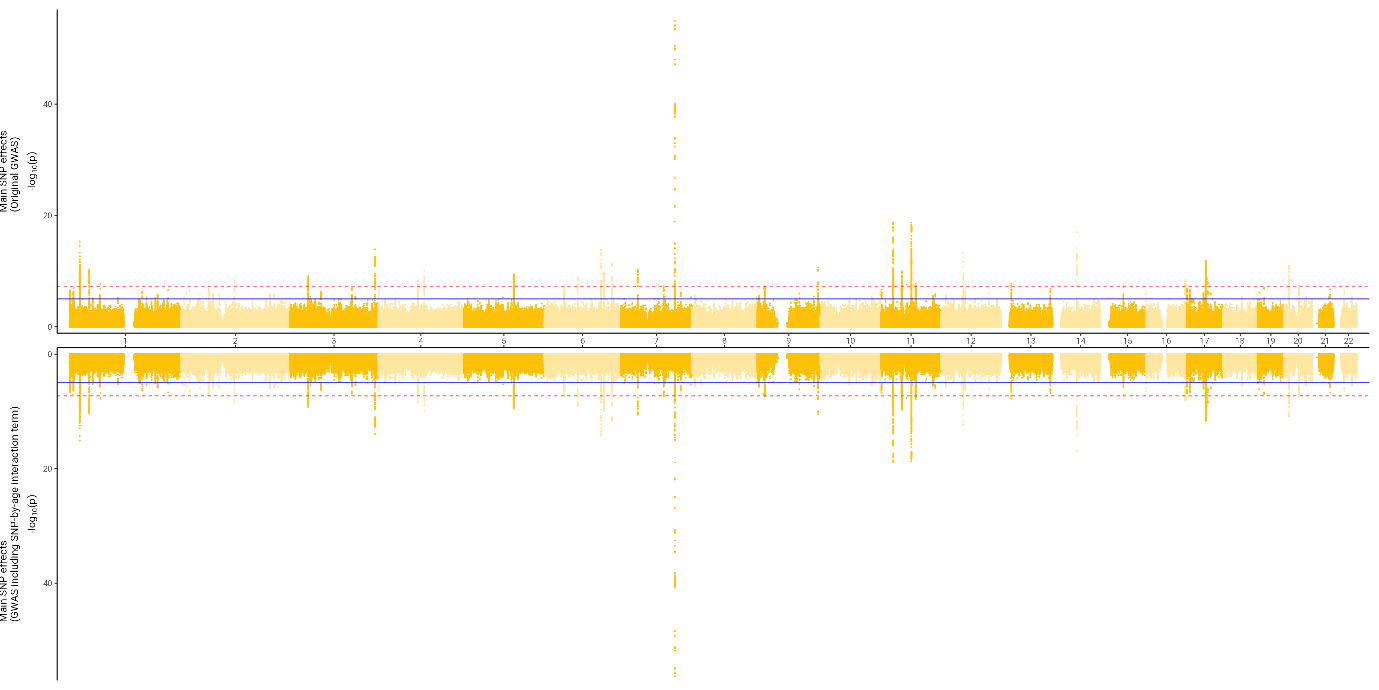

*SFig.21.* Miami plots for **ratio score** contrasting main SNP effect *p*-values from original GWAS with main SNP effect *p*-values from GWAS analyses including an age interaction term (SNP filters: MAF > 0.01, INFO > 0.9, diallelic)

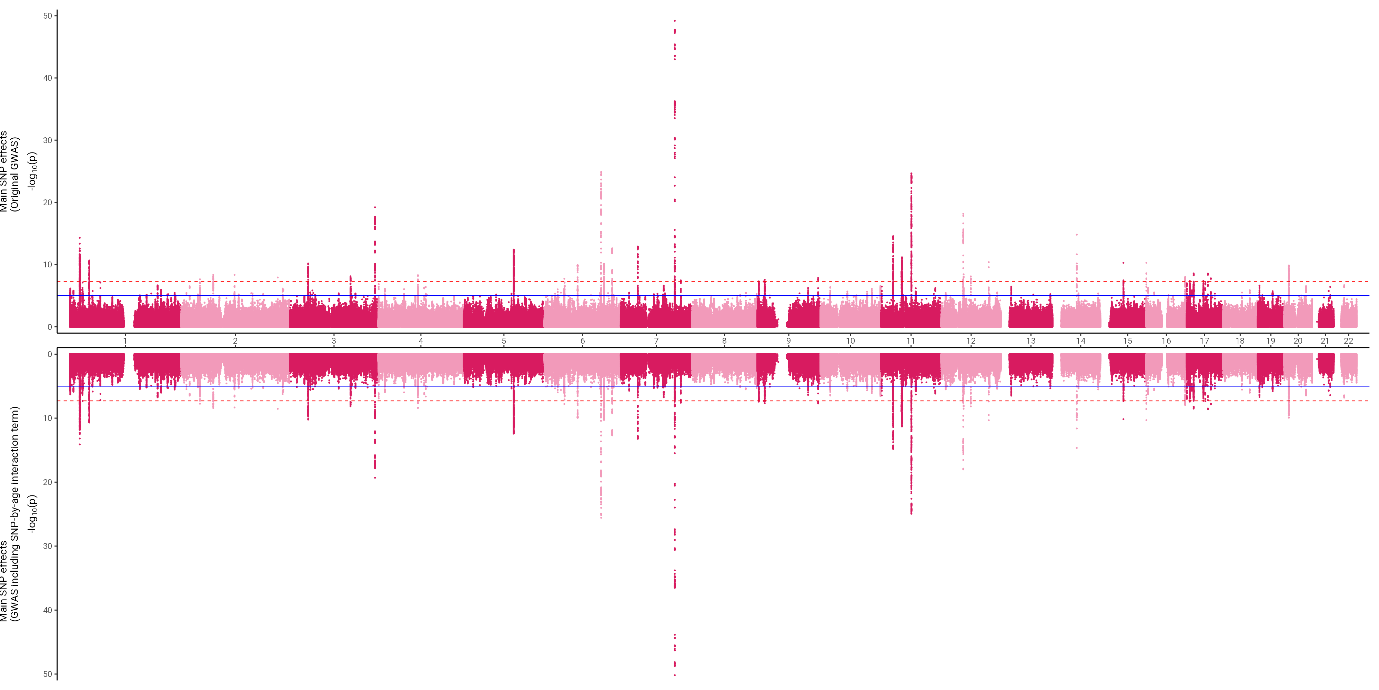

*SFig.22.* Miami plots for **difference score** contrasting main SNP effect *p*-values from original GWAS with main SNP effect *p*-values from GWAS analyses including an age interaction term (SNP filters: MAF > 0.01, INFO > 0.9, diallelic)

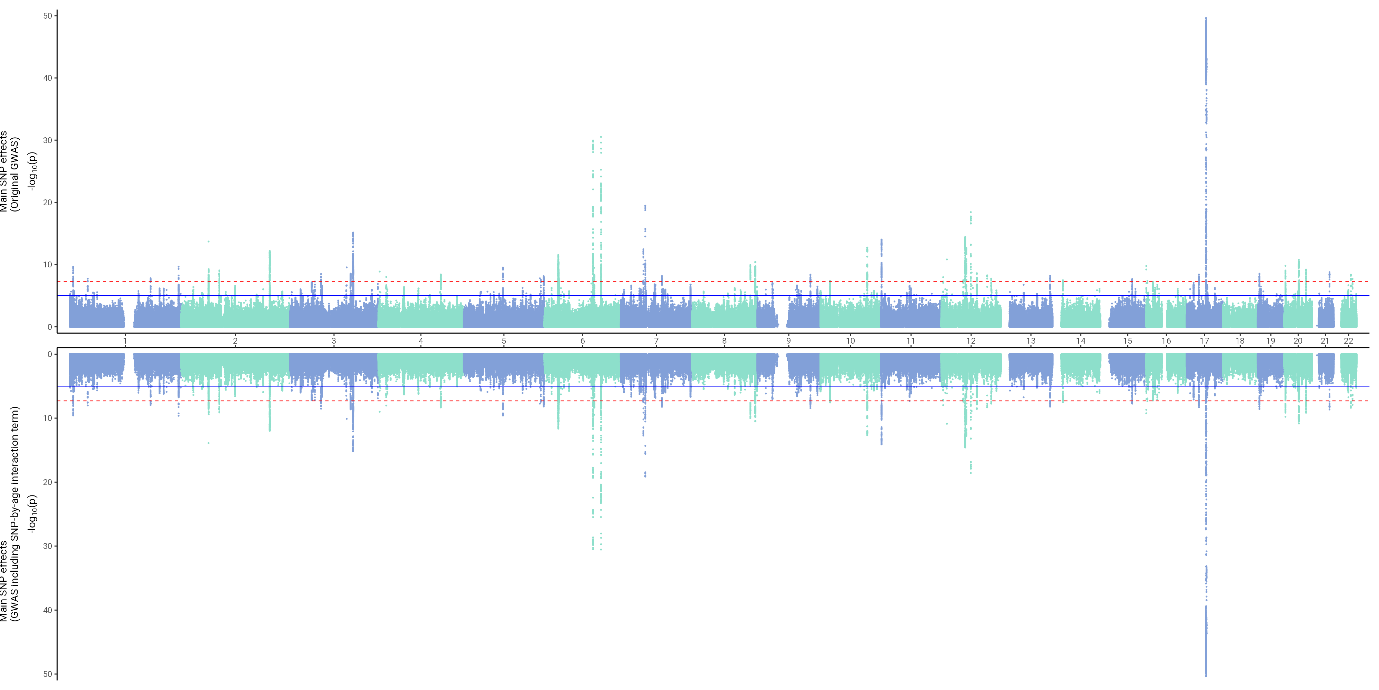

*SFig.23.* Miami plots for **TBV** contrasting main SNP effect *p*-values from original GWAS with main SNP effect *p*-values from GWAS analyses including an age interaction term (SNP filters: MAF > 0.01, INFO > 0.9, diallelic)

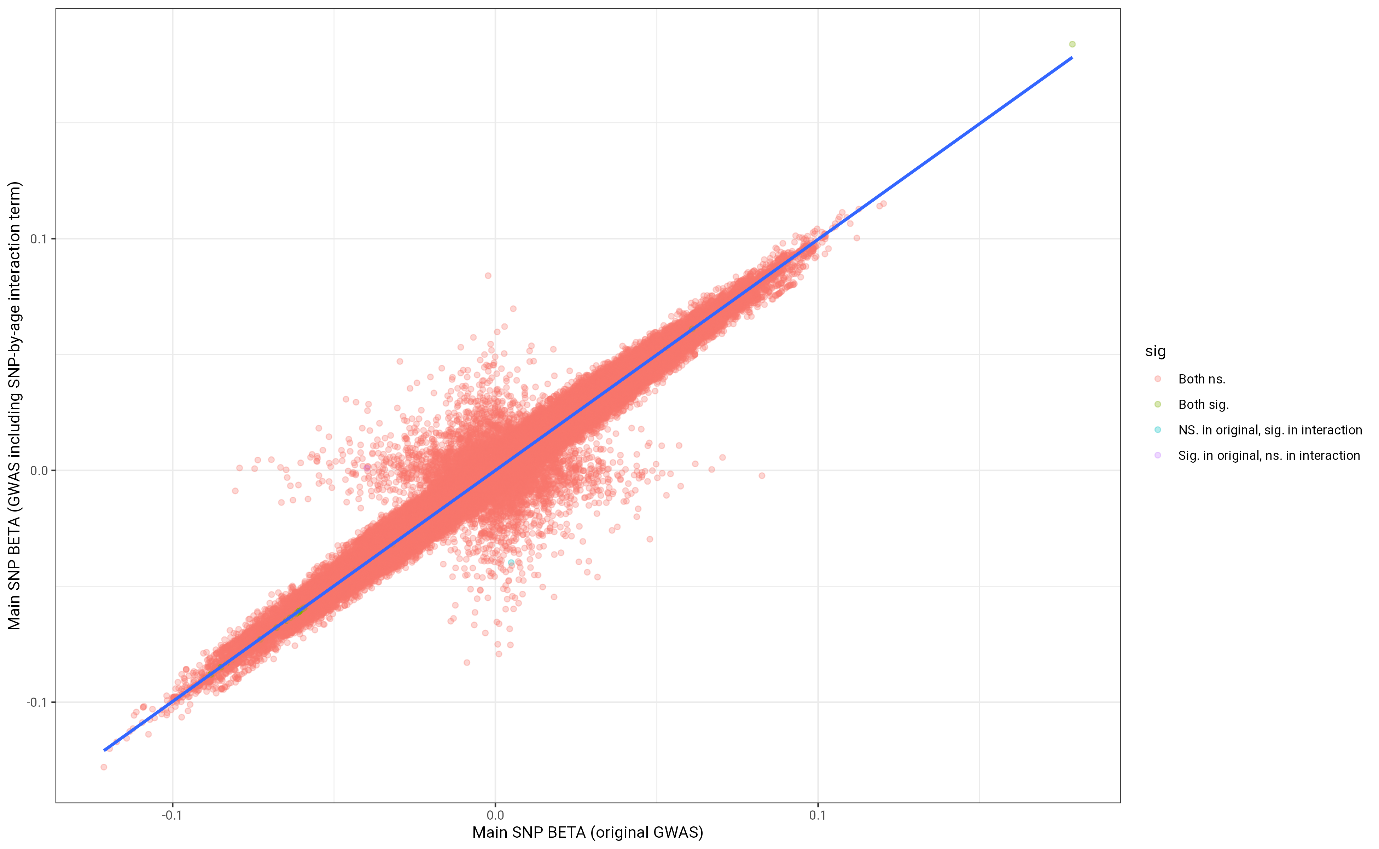

*SFig.24.* **Residual score**: Contrasting main SNP beta effect sizes from original GWAS with main SNP beta effect sizes from GWAS analyses including an age interaction term (SNP filters: MAF > 0.01, INFO > 0.9, diallelic)

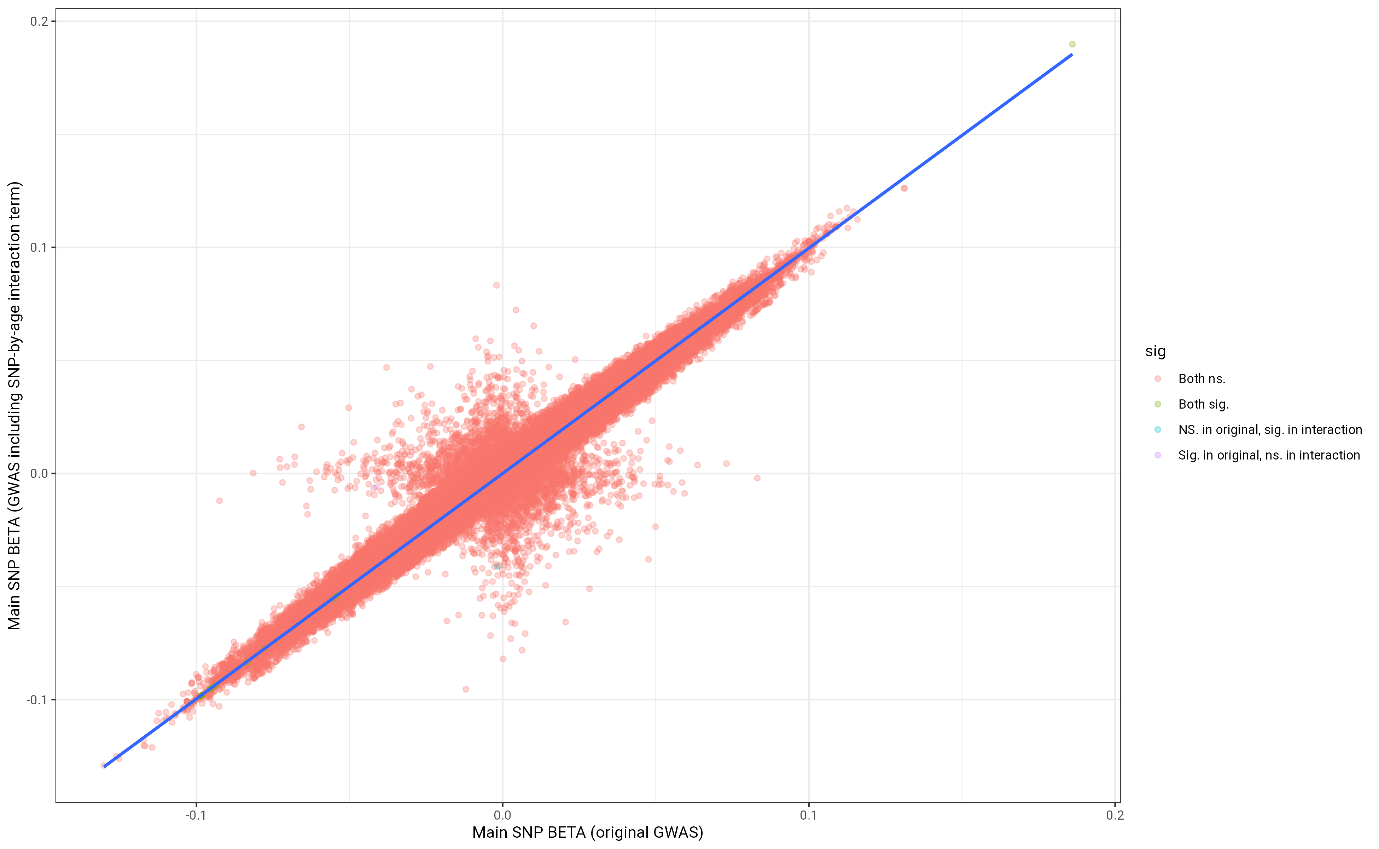

*SFig.25.* **Ratio score**: Contrasting main SNP beta effect sizes from original GWAS with main SNP beta effect sizes from GWAS analyses including an age interaction term (SNP filters: MAF > 0.01, INFO > 0.9, diallelic)

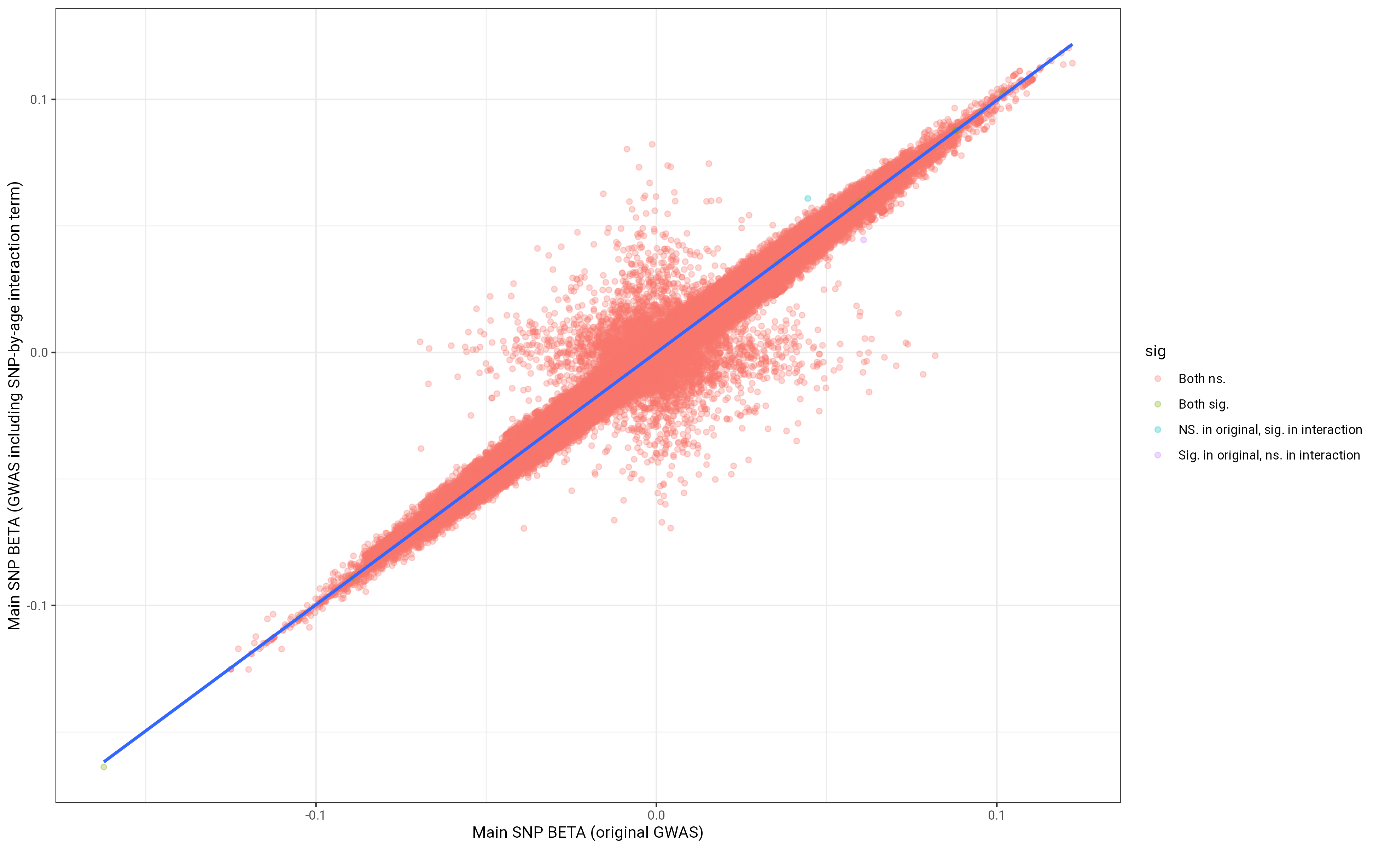

*SFig.26.* **Difference score**: Contrasting main SNP beta effect sizes from original GWAS with main SNP beta effect sizes from GWAS analyses including an age interaction term (SNP filters: MAF > 0.01, INFO > 0.9, diallelic)

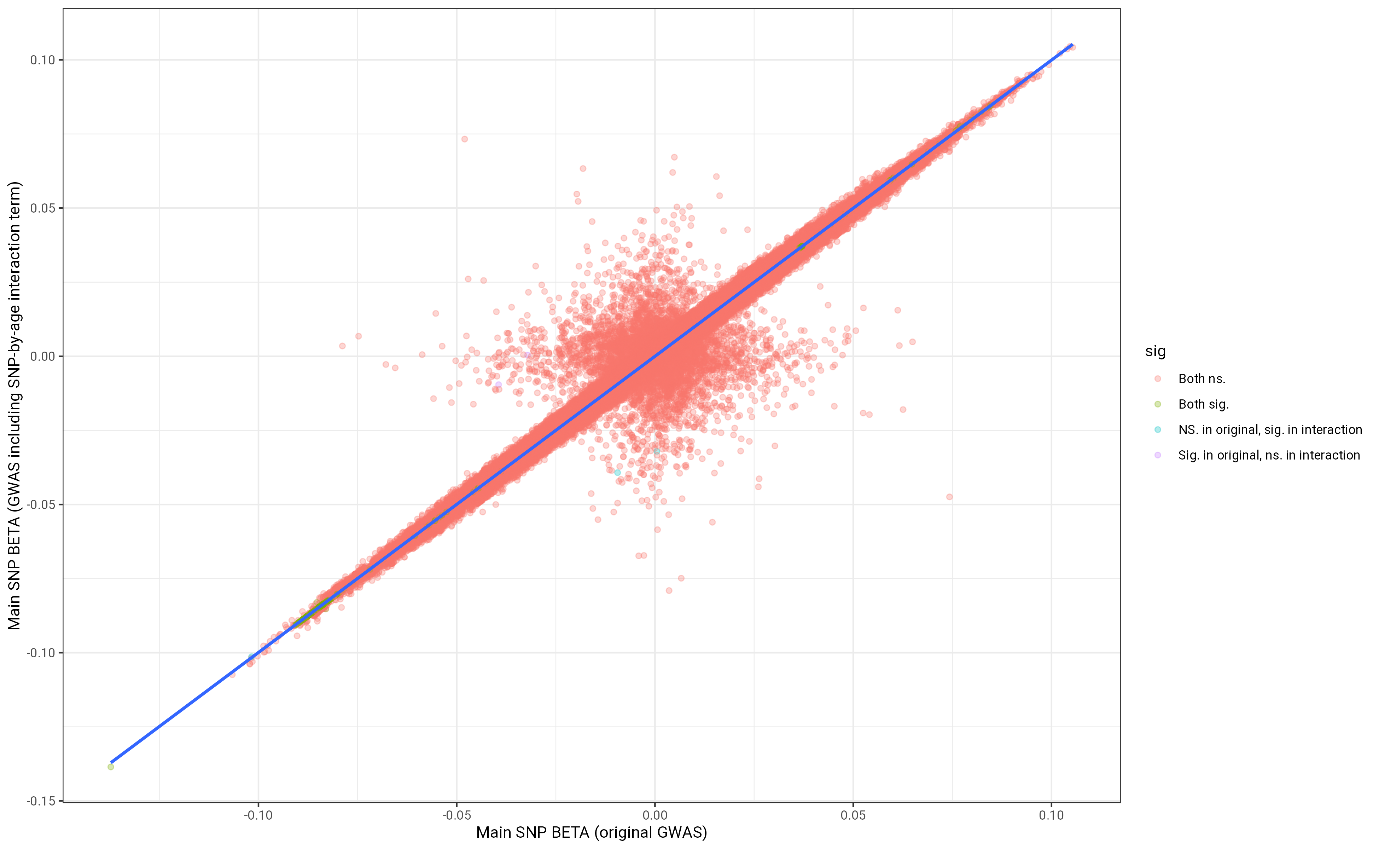

*SFig.27.* **TBV**: Contrasting main SNP beta effect sizes from original GWAS with main SNP beta effect sizes from GWAS analyses including an age interaction term (SNP filters: MAF > 0.01, INFO > 0.9, diallelic)

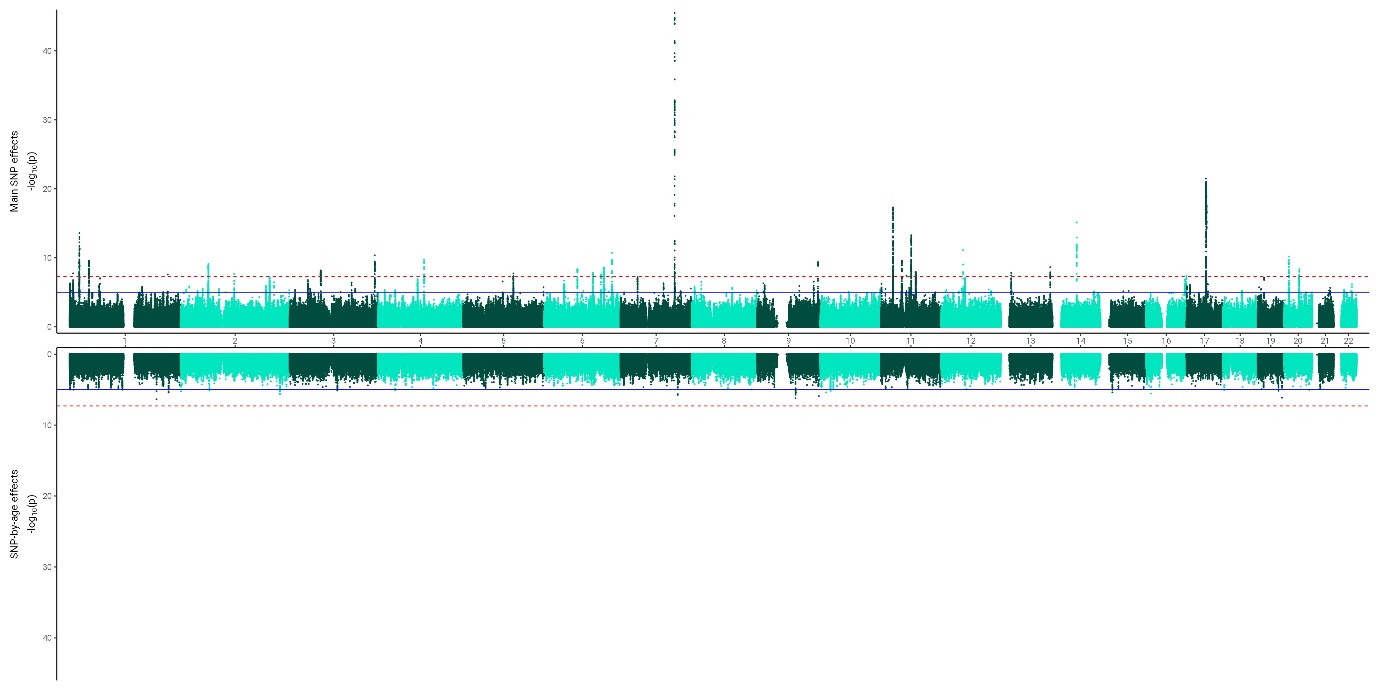

*SFig.28.* Miami plots for **residual score** contrasting main SNP effect *p*-values from original GWAS with SNP-by-age interaction *p*-values from GWAS analyses including an age interaction term (SNP filters: MAF > 0.01, INFO > 0.9, diallelic)

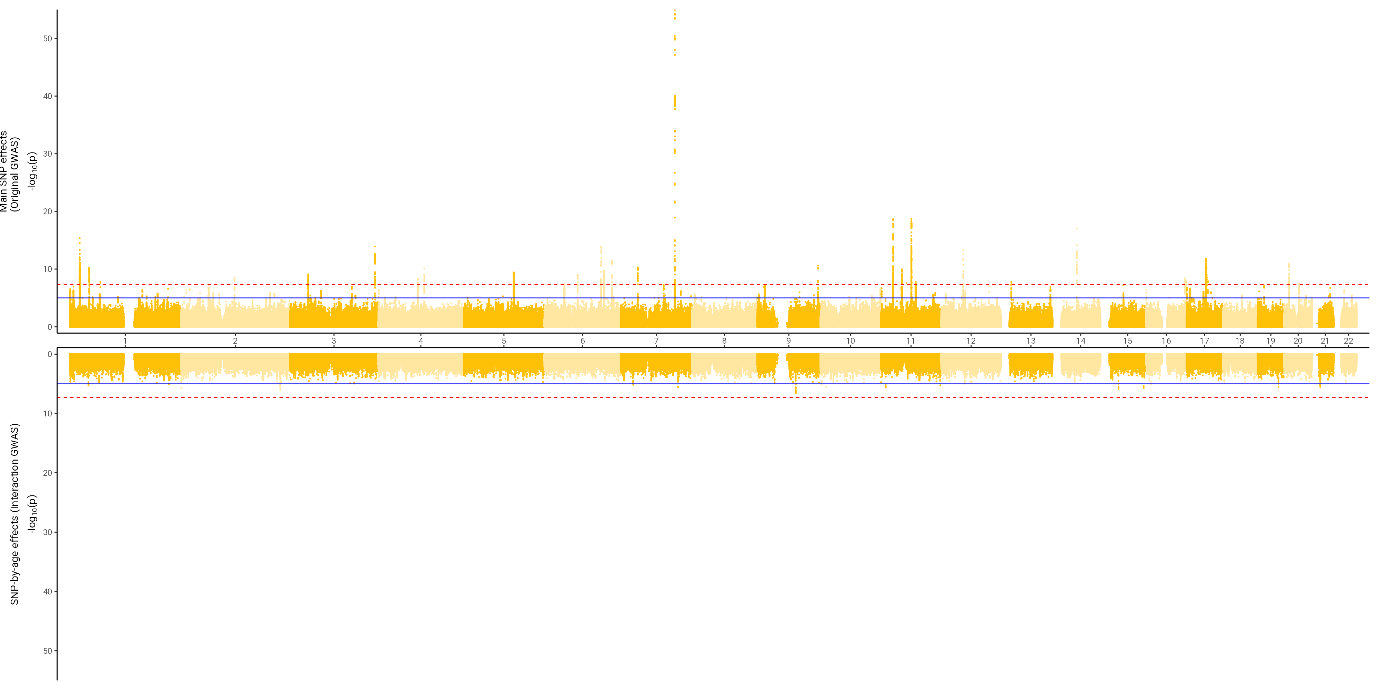

*SFig.29.* Miami plots for **ratio score** contrasting main SNP effect *p*-values from original GWAS with SNP-by-age interaction *p*-values from GWAS analyses including an age interaction term (SNP filters: MAF > 0.01, INFO > 0.9, diallelic)

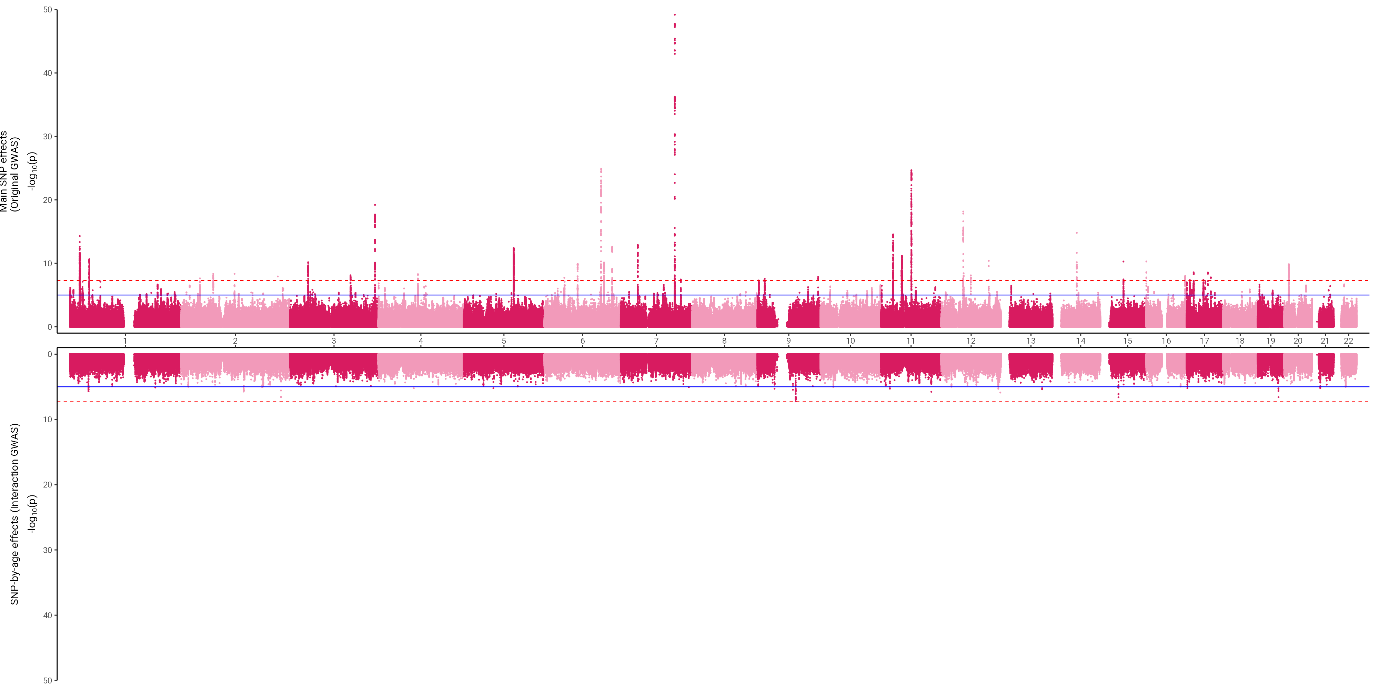

*SFig.30.* Miami plots for **difference score** contrasting main SNP effect *p*-values from original GWAS with SNP-by-age interaction *p*-values from GWAS analyses including an age interaction term (SNP filters: MAF > 0.01, INFO > 0.9, diallelic)

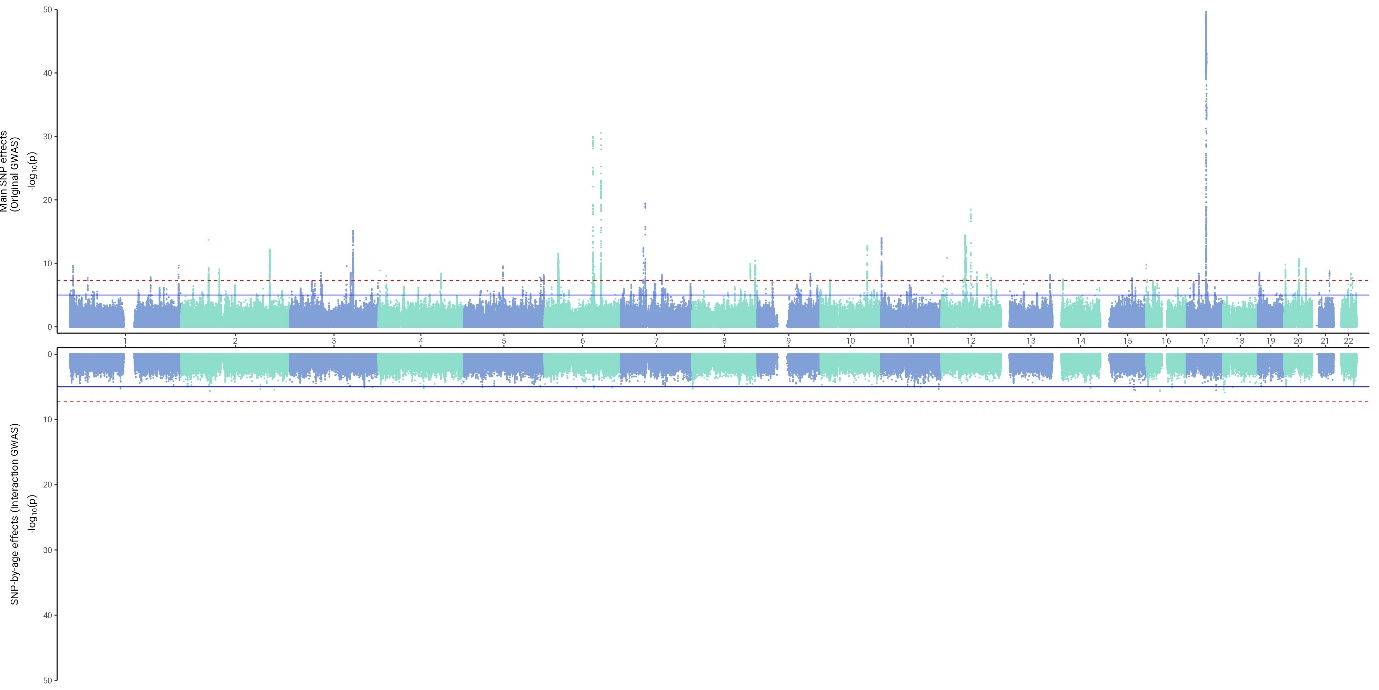

*SFig.32.* Miami plots for **TBV** contrasting main SNP effect *p*-values from original GWAS with SNP-by-age interaction *p*-values from GWAS analyses including an age interaction term (SNP filters: MAF > 0.01, INFO > 0.9, diallelic)

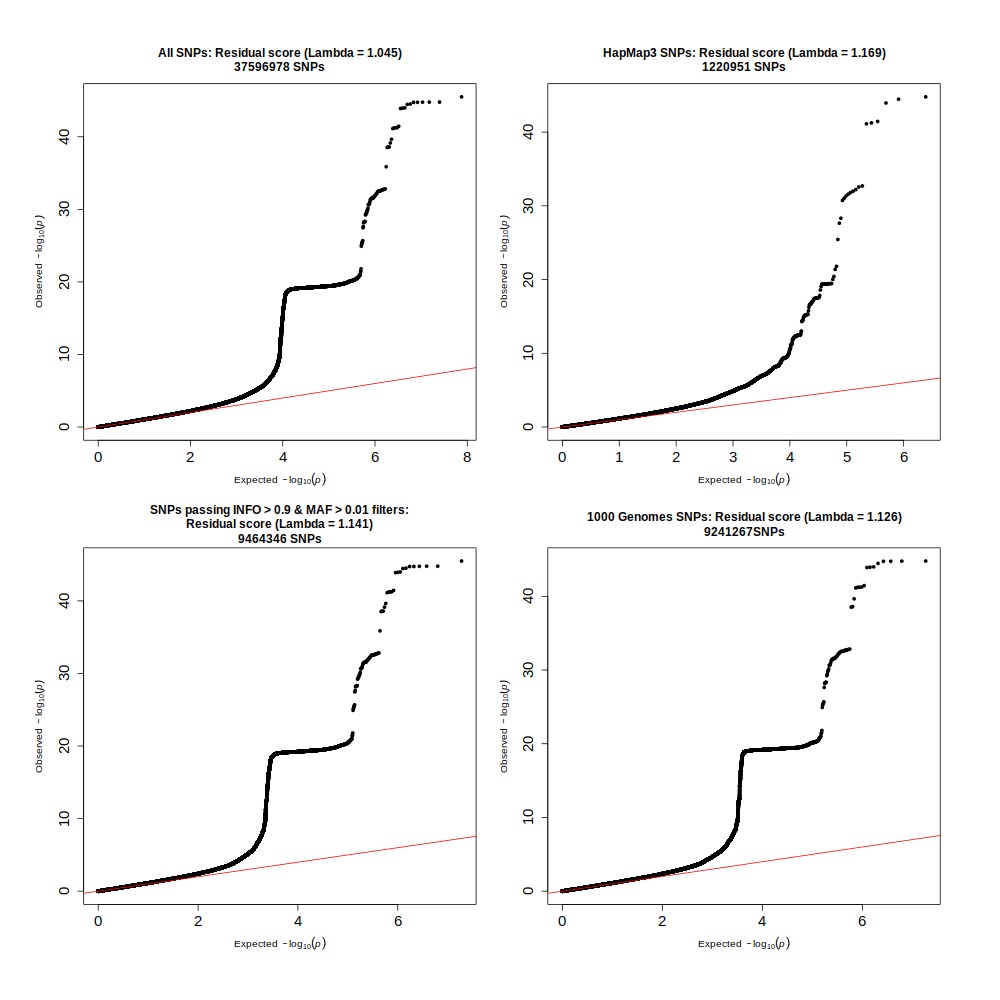

*SFig.33.* QQ plots for residual score including **top left:** all SNPs, **top right:** only HapMap3 SNPs, **bottom left:** SNPs that pass an INFO filter > 0.9, and **bottom right:** MAF filter > 0.1, and 1000 Genomes SNPs. The step in the QQ plot, often present in GWAS QQ plots of neuroimaging phenotypes, disappears when considering HapMap SNPs only

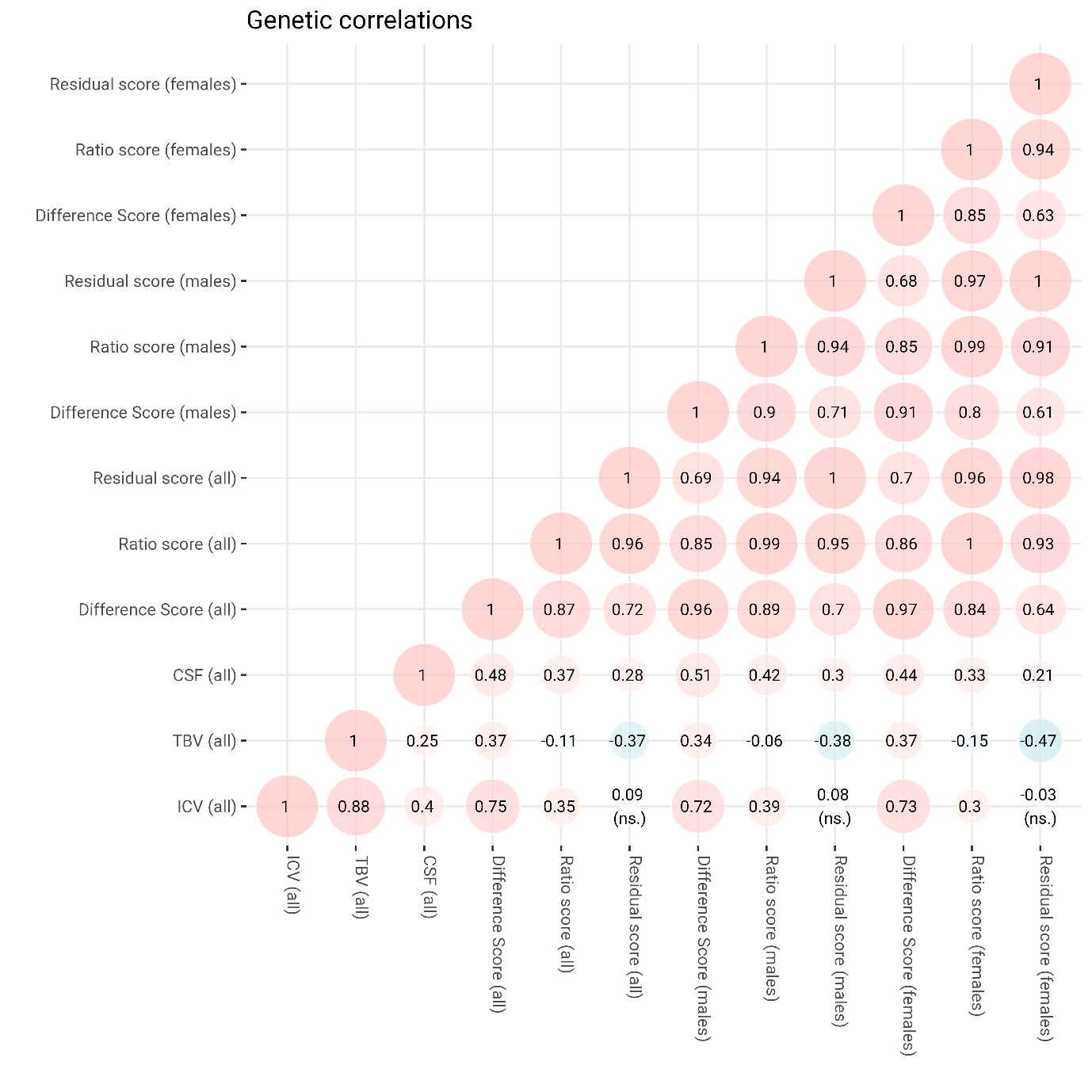

*SFig.34*. Genetic correlations calculated from GWAS summary statistics in the whole UKB sample (all; *N* = 43,110), males only (*N* = 20,453), and females only (*N* = 22,657). Dotted box highlights male-female genetic correlations. LDSC heritability (SE) were LBA _difference_ = 0.37 (0.03), LBA _ratio_ = 0.30 (0.03), LBA _residual_ = 0.27 (0.03) for males, and LBA _difference_ = 0.27 (0.03), LBA _ratio_ = 0.24 (0.03), LBA _residual_ = 0.25 (0.03) for females.
