## SupplementaryMaterials for "Lifetime brain atrophy estimated from a single MRI: measurement characteristics and genome-wide correlates"

**Supplementary Materials**

### **Supplementary information for the Introduction**

#### Description of visual rating scales

There are already-available approaches for quantifying lifetime brain atrophy (LBA) from a single MRI scan. For example, in neuroradiological settings, visual rating scales aid identifying atrophy based on the extent of visible features from a single scan. Well-validated visual scales rate medial temporal lobe atrophy (MTA; Scheltens et al., 1995; Scheltens et al., 1992), global cortical atrophy (GCA; Pasquier et al., 1996), and posterior atrophy (PA; Koedam et al., 2011). In research settings, computational approaches to measuring total brain atrophy from a single MRI scan are common (Bu et al., 2021; Good et al., 2002; Kilsdonk et al., 2015). Computational approaches concur with visual rating scales, but offer greater statistical power since they capture more interindividual variability (not being restricted to a smaller – and more practical – ordinal scale), and offer comparatively greater fidelity to index subtle adulthood atrophic changes in non-clinical samples (Velickaite et al., 2020).

#### *Box 1.* Computational approaches used to infer lifetime brain atrophy (LBA) from either a single cross-sectional MRI scan, or observed atrophic changes from two repeated longitudinal MRI scans

| **Method** | **Interpretation** | **LBA  (i**nferred from a single cross-sectional MRI scan**)** | **Observed atrophic changes (i**nferred from two repeated longitudinal MRI scans**)** | **Advantages** | **Disadvantages** |
| --- | --- | --- | --- | --- | --- |
| Difference score | Captures absolute brain matter losses that occurred since the brain maximally filled the skull  Larger values denote greater brain atrophy in *mm^3^* | **LBA _difference_** = *ICV* - *TBV* | **Observed atrophic changes _difference_ =**  *TBV_time1_* - *TBV_time2_* | Intuitive to interpret in *mm^3^*  Each participant’s value can be computed independently of the study sample | Difference scores where one value is subtracted from the other are correlated with the initial value by construction (e.g., Clifton & Clifton, 2019)  Difference scores conflate the unreliability of the two contributing measures (Cronbach & Furby, 1970; Lord, 1956) particularly in instances where baseline and subsequent measures are moderately-to-strongly correlated, such as TBV and ICV (Rogosa & Willett, 1983). Thus, we expect that *ICV minus TBV* may largely index baseline levels of brain size – as opposed to brain changes only – and any resultant GWAS of this measure may mainly reflect SNPs related to head size. |
| Ratio score | Captures proportional brain matter losses from when TBV maximally filled the skull  Sometimes referred to as the *parenchymal volume fraction*, or *brain parenchymal fraction* (Rudick et al., 1999)  Smaller values denote greater brain atrophy* | **LBA _ratio_** = *TBV* / *ICV* | **Observed atrophic changes _ratio_** =  *TBV_time2_* / *TBV_time1_* | Easy to interpret as proportional values between 0 and 1  Each participant’s value can be computed independently of the study sample | Ratio scores also suffer from a mathematical coupling issue (Archie, 1981). This means, just as with difference scores (above), the ratio score of atrophy is likely correlated with ICV, and hence captures baseline differences in brain size, in addition to proportional brain matter losses |
| Residual score | A residual score captures the difference between an individual’s observed TBV and their predicted TBV given their ICV size (i.e., a negative value means an individuals’ TBV is smaller than expected given their ICV)  Smaller values denote greater brain atrophy* | *TBV*-associated residuals of *ICV*  **LBA _residual_** =  lm(TBV ~ ICV) | *TBV_time2_* -associated residuals of *TBV_time1_*  **Observed atrophic changes _residual_ = lm(***TBV_time2_ ~ TBV_time1_***)** | Residual scores are uncorrelated with baseline differences in head size (i.e., ICV in single-occasion MRI estimation or TBV_time1_ in longitudinal estimation) | Residual scores could be categorised as less intuitive and less concrete than the two methods above  A regression-based approach means that an individual’s residuals will differ as a function of the overall sample composition which dictates the regression line from which the residuals deviate. This makes the resulting residual score susceptible to sources of bias such as overfitting (when computed in small samples) and variance inflation (where the covariate is highly correlated with the exposure when used in a multivariable setting) which can affect regression estimates |

* Throughout the paper, residual and ratio score values have been flipped (i.e., multiplied with -1) to enable a unified interpretation across all three methods whereby larger values denote greater brain atrophy.

#### *Supplementary Table 1.* Descriptive statistics of TBV, ICV and lifetime brain atrophy in younger (MRi-Share, HCP) and older adults (UKB, LBC)

| Statistic | HCP | MRi-Share | STRADL | UKB  (initial scan) | UKB (second scan) | LBC1936 (initial scan) | LBC1936 (fourth scan) |
| --- | --- | --- | --- | --- | --- | --- | --- |
| *N* | 800 | 1831 | 987 | 4674 | 4674 | 634 | 286 |
| Age in years:  Mean (range) | 27 (22-31) | 22 (18-35) | 60 (26-84) | 62 (46-81) | 65 (49-83) | 73 (71-74) | 82 (81-83) |
| TBV: Mean (SD) | 1173.81 (120) | 1131.88 (100) | 1101.77 (110 | 1186.03 (110) | 1172.09 (110) | 1011.23 (97) | 935.48 (92) |
| ICV: Mean (SD) | 1606.3 (170) | 1568.07 (140) | 1414.44 (210) | 1555.14 (150) | 1547.39 (150) | 1396.53 (140) | 1382.16 (150) |
| cor(ICV,TBV) | 0.92 ^[1]^ | 0.96 | 0.77 | 0.90 | 0.89 | 0.81 | 0.75 |
| Difference score |  |  |  |  |  |  |  |
| Mean (SD) | 432.49 (79.34) | 436.18 (48.29) | 312.67 (144.35) | 369.11 (69.41) | 375.6 (72.23) | 385.31 (86.88) | 446.68 (101.51) |
| Median | 437.57 | 431.42 | 353.16 | 363.11 | 369.36 | 382.43 | 444.7 |
| Range | 7.9 to 644.09 | 298.44 to 630.19 | 2.1 to 660.4 | 6.01 to 834.2 | 120.02 to 739.69 | 53.15 to 664.4 | 100.58 to 882.81 |
| Variance | 6300 | 2300 | 21000 | 4800 | 5200 | 7500 | 10000 |
| Ratio score |  |  |  |  |  |  |  |
| Mean (SD) | 0.73 (0.04) | 0.72 (0.02) | 0.79 (0.09) | 0.76 (0.03) | 0.76 (0.03) | 0.73 (0.05) | 0.68 (0.05) |
| Median | 0.73 | 0.72 | 0.76 | 0.76 | 0.76 | 0.73 | 0.68 |
| Range | 0.65 to 0.99 | 0.64 to 0.78 | 0.61 to 1 | 0.6 to 1 | 0.59 to 0.9 | 0.6 to 0.95 | 0.48 to 0.9 |
| Variance | 0.0012 | 0.00037 | 0.0074 | 0.0011 | 0.0011 | 0.0021 | 0.0027 |
| Residual score |  |  |  |  |  |  |  |
| Mean (SD) | 0 (48.09) | 0 (30.06) | 0 (71.1) | 0 (49.21) | 0.00 (50.64) | 0 (57.37) | 0 (61.08) |
| Median | -7.2 | 2.29 | -6.29 | 0.99 | 0.69 | 0.46 | 3.26 |
| Range | -128.64 to 238.91 | -145.08 to 90.54 | -192.39 to 252.4 | -301.01 to 317.80 | -274.31 to 157.87 | -175.03 to 178.61 | -259.78 to 151.3 |
| Variance | 2300 | 900 | 5100 | 2400 | 2600 | 3300 | 3700 |

### **Supplementary Results**

#### Repeated LBA measures increase with chronological age in the LBC1936

Repeated MRI measures in the age-homogeneous LBC1936 showed that LBA increases with advancing age, even when MRI measures at each visit were processed independently of each other (i.e., here MRI data was processed with the FS cross-sectional rather than the longitudinal processing stream; *SFig.12*). According to all three computational methods, participants showed significantly more LBA at age 82 than at age 73 years [difference score: mean at age 70 (SD) = 375 (82) *mm^3^*, mean at age 79 (SD) = 446 (102) *mm^3^*; ratio score: mean at age 70 (SD) = 0.73 (0.04); mean at age 79 = 0.68 (0.05); residual score ^[2]^: mean at age 70 (SD) = 40 (53); mean at age 79 = -37 (58); t-test *p-*values for the three scores < 5x10^-14^]. This section was conceived to compliment *Fig.3* in the main manuscript.

#### Supplementary moderation analysis

To better understand the relationship between lifetime brain atrophy and longitudinal atrophic changes, we conducted a moderation analysis in the UKB to investigate whether the association between observed atrophic changes (capturing change between time 1 and time 2) and lifetime brain atrophy (at time 2), is moderated by the amount of lifetime brain atrophy that occurred prior to a participants’ first MRI scan (at time 1). We hypothesised that the strongest correlation between those measures would occur when participants exhibit minimal atrophy at time 1, in which case atrophic changes and lifetime atrophy at time 2 would cover approximately the same timeline. Conversely, when substantial atrophy already occurred prior to time 1, observed atrophic changes would likely capture fewer changes compared with lifetime measures because the latter should reflect the entire span of atrophic effects ever experienced – assuming that brain atrophy declines continually and linearly.

To perform this analysis, we computed an interaction term by multiplying variables for lifetime brain atrophy at time 2 and longitudinal atrophic changes between time 1 and time 2. We then fitted a multiple regression with lifetime brain atrophy at time 2 as the outcome, and longitudinal atrophic changes between time 1 and time 2 and the interaction term as predictor variables. The interaction term was significant in a model considering atrophy scores estimated with the difference method (*b* = 0.08, SE = 0.01, *p* = 1.88x10^-7^), but they were non-significant in a model considering atrophy scores estimated with the ratio method (*b* = 0.02, SE = 0.01, *p* = 0.124) or the residual method (*b* = 0.01, SE = 0.01, *p* = 0.337). This suggests that those with greater LBA _difference score_ prior to the first instance of neuroimaging showed a weaker association between LBA _difference score_ and observed longitudinal atrophy.

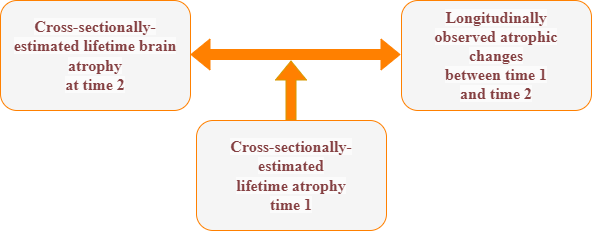

*Illustration of the moderation analysis.* Directed acyclic graph illustrating moderation analysis. This moderation suggests that the association between lifetime atrophy at time 2 and longitudinal atrophic changes between time 1 and time 2 is mediated by an individuals initial lifetime atrophy at time 1

#### Aim 3: Associations with ageing-related variables

Our pre-registered Aim 3 was designed to assess the external validity of cross-sectionally-estimated lifetime brain atrophy by quantifying the extent to which lifetime atrophy captures variance predictive of cognitive- and other ageing-related phenotypes (presented in Fig.2 in main manuscript for LBC1936 and SFig.7 for UKB). The pre-registered 3.1-3.4 analyses below aimed to further evaluate the nature of their relationship. We suspect that particularly the analyses in 3.2-3.3 suffer from variance inflation because we were including TBV, ICV or a variable derived from TBV and ICV in the same model. Results do not seem intuitive and it is possible that estimates were instable due to variance inflation, or other biases introduced by performing a regression that included both a derived measure (such as LBA _difference_, LBA _ratio_ or LBA _residual_) and a baseline measure on which basis the first measure was derived (e.g., Butler et al., 2021; Glymour et al., 2005). We have therefore not further interpreted the results displayed in the plots below. They are only included here for completeness as they had been pre-registered.

##### Aim 3.1: Amplifier effect

1. Health-related phenotypes ~ lifetime atrophy *time 2*
2. Health-related phenotypes ~ TBV *time 2*

Our lifetime brain atrophy definition using the residual score considers the residuals of TBV adjusted for ICV, and this analysis 3.1 demonstrated that this procedure resulted in an amplifier effect whereby the residuals explain more variance than the original/ unadjusted variable (i.e., TBV). We showed the existence of this amplifier effect in the LBC1936 whereby considering variance explained in an ageing-related trait we would sensibly expect to be associated with lifetime brain atrophy (e.g., visual rating scales, cognitive slope). We did not see evidence of an amplifier effect when considering variance in another trait that we would not expect to be associated with lifetime brain atrophy (e.g., cognitive intercept).

To illustrate the amplifier effect, we contrasted estimates of variances explained (*R^2^*) from two models: the first predicting health outcomes with the residuals of TBV adjusted for ICV, and the second predicting the same health outcomes with TBV alone. Those residuals were obtained from either a difference, ratio, or residual method. *R^2^* estimates are plotted in Figure below.

Three overarching results from the LBC1936 are discussed below by focusing on lifetime brain atrophy scores obtained from the residual method, because it is the one score (out of the three computational methods) that is fully uncorrelated with ICV (i.e., baseline differences in head size). First, lifetime atrophy (*R^2^* = 11-20%), but not TBV alone (*R^2^* = 0.01-0.35%) are associated with visually rated atrophy, confirming that the residuals indeed carry information beyond simple baseline differences in TBV, that is relevant to clinically measured brain atrophy. Second, lifetime atrophy calculated with the residual method explained more than double the variance in a slope of cognitive function (*R^2^* = 13%) than can be explained with TBV alone (*R^2^* = 5%). This illustrates that adjusting for ICV acts as an amplifier in predicting a variable that is expected to be associated with neurodegeneration, but less so with baseline differences in head size. It complements this observation that TBV alone explains more variance in baseline cognitive ability (*R^2^* = 12%) than lifetime atrophy (*R^2^* = 0.02-8%), suggesting that atrophy scores indeed index variance related to change, and less so baseline differences in cognitive function. With respect to other traits for which we had longitudinal measures available and modelled intercepts and slopes, at least numerically it appears that estimated atrophy tends to explain more variance in slopes (i.e., change), and TBV alone explains more variance in intercepts (i.e., baseline).

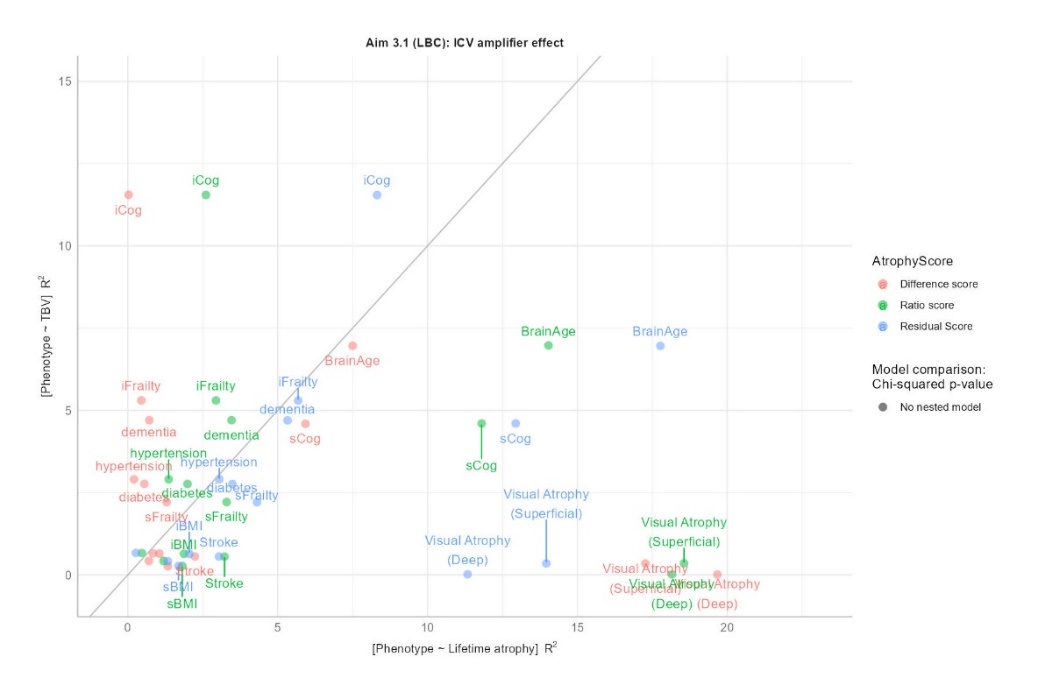

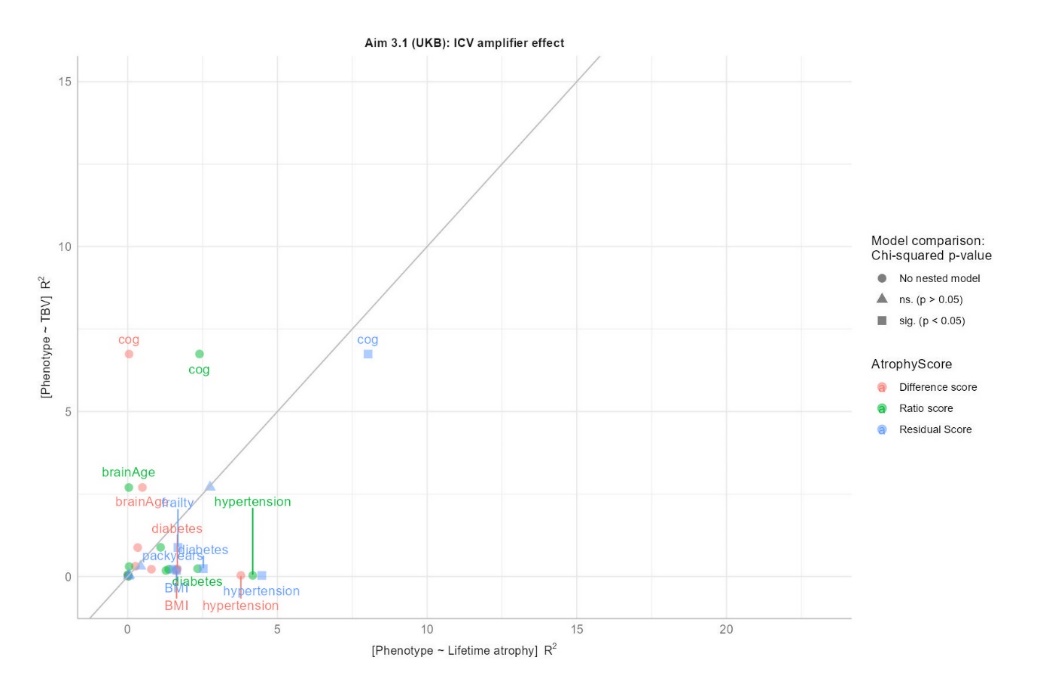

Linear regression x~y:

- UK Biobank: intercept = 1.17 (p = 0.003), slope = 0.29 (*p* = 0.078)
- LBC1936: intercept = 5.31 (*p* = 9.25x10^-5^), slope = 0.01 (*p* = 0.95)

##### Aim 3.2: Does lifetime atrophy (LBA) explain variance above and beyond baseline variables TBV and ICV?

1. Health-related phenotypes ~ *ICV time 2* (or *TBV time* 2) + *cross-sectional atrophy time 2*
2. Health-related phenotypes ~ *ICV time 2* (or *TBV time* 2)

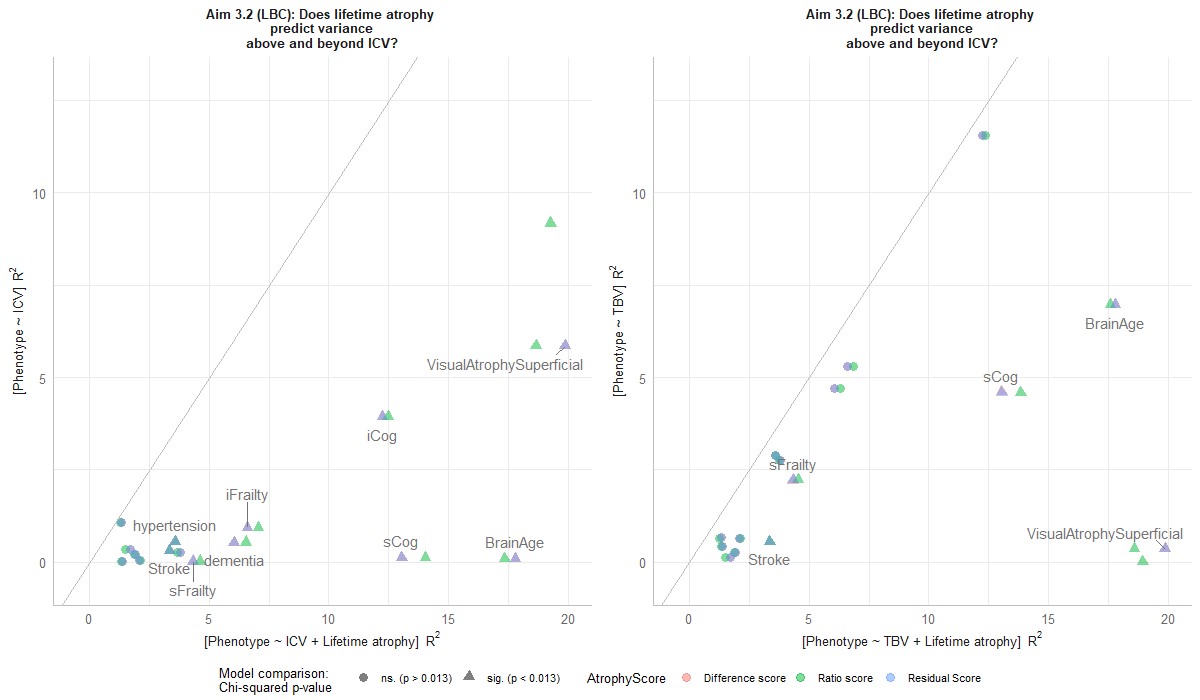

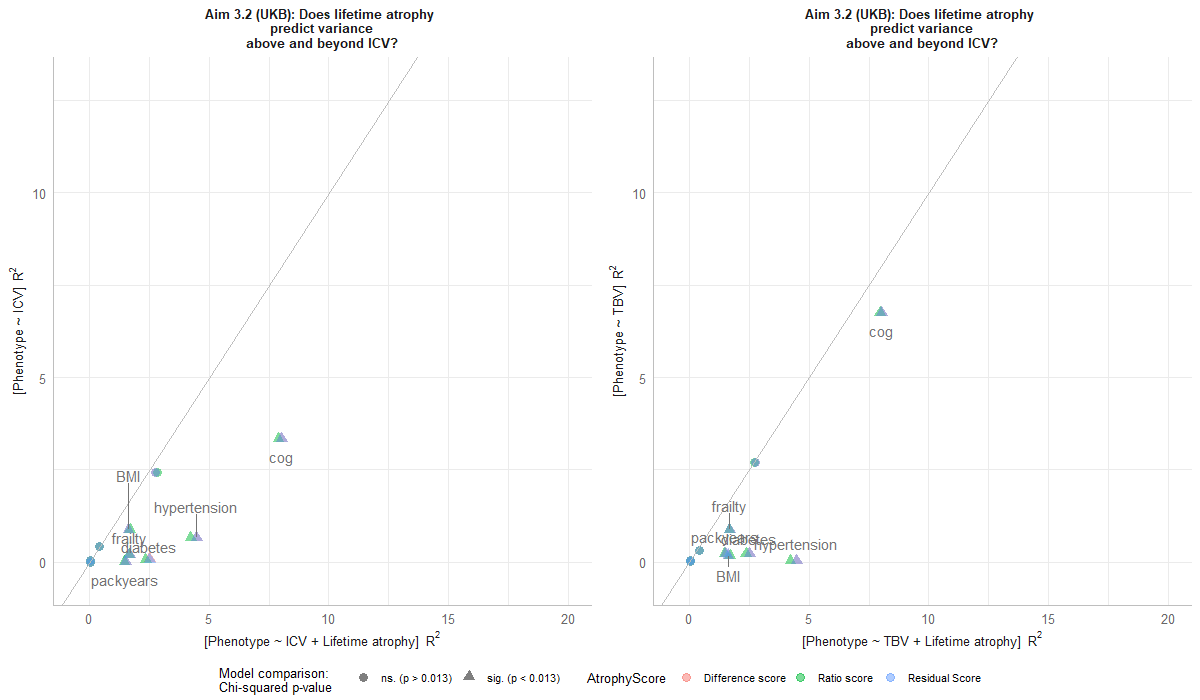

Linear regression x~y:

- UK Biobank (TBV analysis): intercept = 1.23 (*p* = 0.0001), slope = 0.94 (*p* = 1.7x10^-8^)
- UK Biobank (ICV analysis): intercept = 0.96 (*p* = 0.006), slope = 1.66 (*p* = 1.34x10^-7^)
- LBC1936 (TBV analysis): intercept = 5.55 (*p* = 4.14x10^-5^), slope = 0.69 (*p* = 0.024)
- LBC1936 (ICV analysis): intercept = 4.73 (*p* = 2.47x10^-7^), slope = 1.87 (*p* = 9.43x10^-9^)

##### Aim 3.3: Does lifetime brain atrophy explain health phenotype-associated variance above and beyond the variance explained by estimated atrophy?

1. Health-related phenotype ~ (*cross-sectionally) estimated atrophy time 2* + (*longitudinally) observed atrophy*
2. Health-related phenotype ~ (*cross-sectionally) estimated atrophy time 2*

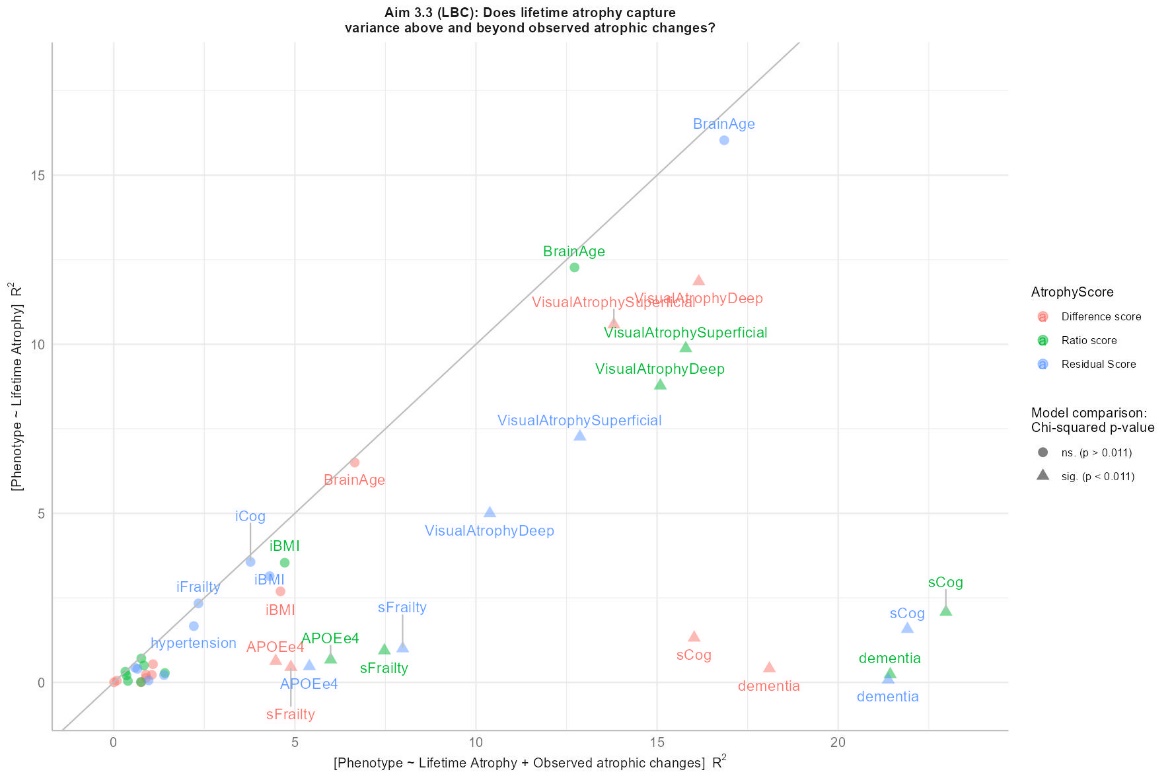

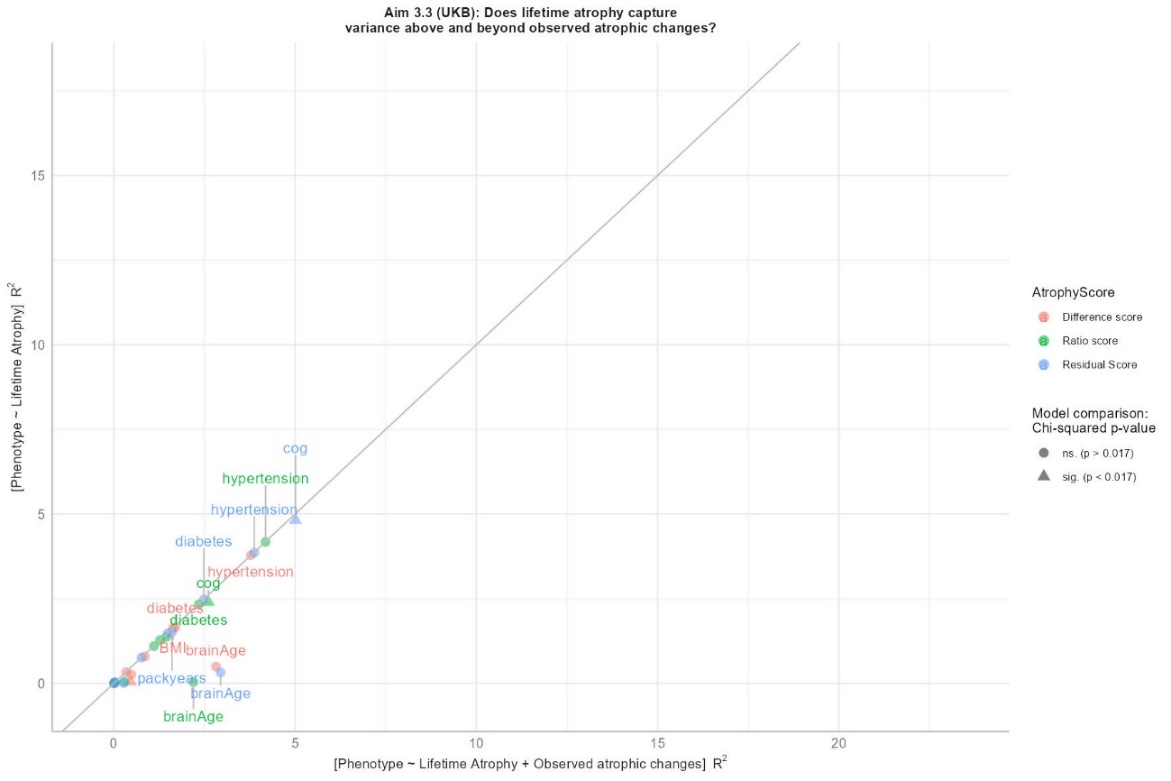

Linear regression x~y:

- UK Biobank: intercept = 0.45 (*p* = 0.0139), slope = 0.88 (*p* = 3.04x10^-10^)
- LBC1936: intercept = 4.62 (*p* = 0.0003), slope = 0.89 (*p* = 0.0007)

##### Aim 3.4: General trends of associations across aims 3.1-3.3

Simply from looking at the plots above, there do not seem to be obvious overarching trends. Linear regressions reporting intercepts and slopes do not deliver strong evidence for consistent trends across UKB and LBC1936. The regression results may be summarised as follows:

- Aim 3.1 (amplifier effect): Both samples yielded non-significant slopes between predicted *R^2^* by TBV alone vs. *R^2^* by LBA. Some outcome traits were more strongly associated with TBV and others were more strongly associated with LBA. For example, in LBC1936, the amplifier effect seemed to affect traits like cognitive slopes and visual atrophy scales but not cognitive intercepts.
- Aim 3.2: In general, LBA added explanatory variance above and beyond variance explained by baseline levels of TBV or ICV, which was indicated by a significant overarching slope in LBC1936 and UKB.
- Aim 3.3: Both cohorts produce different results with regard to whether longitudinally-observed atrophic changes add variance above and beyond variance explained by LBA alone. In UKB, observed atrophic changes do not tend to add explanatory variance above and beyond LBA (slope near 1, and intercept near 0), which may be driven by the limited variance captured in this longitudinal measure (*SFig.8*). In the LBC1936, observed atrophic changes tend to add explanatory variance above and beyond LBA suggesting LBA and observed atrophy changes capture meaningful variance in their own right. This is consistent with the interpretation in the main manuscript that LBA seems to capture longer-term changes of ageing that are not specifically disease-related and that longitudinally-observed atrophic changes capture more short-term brain shrinkage that is associated with dementia.

#### SNP-by-age interaction effects on LBA

Given the strong association between LBA and age, we aimed to explore how SNP effects might be moderated by age through a SNP-by-age interaction analysis. This analysis helps clarify whether the genetic associations we observe are consistently significant across different age groups or if their strength and significance change with advancing age. Interaction tests were calculated in REGENIE (Mbatchou et al., 2021) where we added the entry-wise product of SNP and age to the GWAS model. From this model we extracted the marginal SNP effects (REGENIE label: ADD-INT_SNP) and the SNP-by-age interaction effects (REGENIE label: ADD-INT_SNPxAge) for SNPs that were of primary interest under a polygenic model (minor allele frequency; MAF > 0.01, INFO > 0.9, biallelic). For LBA _residual_, the interaction model produced a very similar-appearing Manhattan plot as the original GWAS model (*SFig.20*). Marginal GWAS SNP effects on LBA remained very similar compared to SNP effects obtained from a model excluding the SNP-by-age interaction term: the main SNP betas from the original GWAS vs. the marginal SNP effects from the interaction GWAS were perfectly correlated (intercept = 0, slope = 1; *SFig.24*). Zero SNPs showed significant SNP-by-age interaction effects (*p* < 5 x10^-8^; *SFig.29*), which included both *APOE* SNPs rs7412 and rs429358 (*p* *_rs7412_* = 0.195; *p _rs429358_* = 0.660). This could be due to the typical lack of power to detect modest SNP-by-age interaction effect sizes. Those SNP-by-age interaction trends were very similar for both the difference and ratio scores, as well as TBV (*SFig.20-32*). The lack of SNP-by-age interaction effects may suggest that our measure of brain change (i.e., LBA) is largely independent of age meaning that age-associated variance has been absorbed and is represented by the LBA measure.

Additionally, we calculated the LBA variance explained by SNP-by-age interactions using GCTA (Yang et al., 2011) to obtain an estimate independent of individual SNP effects (*Table S2 below*). Although statistically non-significant in *N* = 38,624, point estimates for the gene-by-age interaction insinuated a sizable contribution to phenotypic variance (34%; *SE* = 0.83), but the large standard errors underlined the lack of power for GxE interaction models. Their reliable detection will require much larger samples.

##### *Supplementary Table 2.* SNP-by-age interaction results

| Traits | GCTA SNP-heritability (SE) | GCTA SNP-by-age † (SE) | GCTA SNP-by-age LRT value |
| --- | --- | --- | --- |
| LBA  (residual score) | 0.41 (0.01) | 0.34 (0.83)  *ns.* | 0.172 |
| LBA  (ratio score) | 0.42 (0.01) | 0.37 (0.82) *ns.* | 0.198 |
| LBA  (difference score) | 0.47 (0.01) | 0.17 (0.82) *ns.* | 0.041 |

Results statistically non-significantly different from zero are marked with *ns* where relevant.
† This GCTA SNP-by-age estimate indicates the phenotypic variance accounted for by the interaction between age and genome-wide common variants. This estimate was derived from a model with three variance components for common genetic variants (i.e., SNP-heritability), SNP-by-age interactions and error. In this three-variance-component model, SNP-heritability was estimated to be the same as the SNP-heritability from a model with only two variance components (common variants + error), but the SNP-heritability standard errors were slightly larger in the three-component model (i.e., increased from 0.01 to 0.02).

1. Same correlation (*r* = 0.93) when considering unrelated HCP individuals only (*n* = 326)
2. In order to illustrate that the residual score indicated greater LBA with advancing age in this age-homogeneous sample, the residual score presented here was calculated across all measurements at 4 time points so that atrophy estimates were derived relative to the same average value. This was required to illustrate time-dependent increases, because all time points would have had the same average value had we derived the residual score for each individual time point independently, as was done for the residual score across the manuscript where only one time point was available. The LBC1936 data in this section was processed with the FS cross-sectional stream (as opposed to the longitudinally processed MRI data) where ICV is *not* held constant across visits, which should mimic a cross-cohort comparison, but likely makes the residual score noisier. It speaks to the validity of LBA that increases in the residual score remain prominent here.
