## SupplementaryMethods for "Lifetime brain atrophy estimated from a single MRI: measurement characteristics and genome-wide correlates"

**Supplementary Methods**

### 2.1 Sample descriptions

#### 2.1.1 Young adults

**Human Connectome Project (HCP).** The HCP is a cohort of healthy young adults (*N* ~ 1,113; age range 22-35) that were recruited from ongoing studies as part of the Missouri Family Study (Van Essen et al., 2012). Exclusion criteria mainly included severe diseases (e.g., epilepsy, multiple sclerosis, cerebral palsy), as well as criteria that would have prohibited safe MRI scanning (e.g., metal or devices in the body, claustrophobia). The HCP analysis team performed MRI data processing (Glasser et al., 2013; Marcus et al., 2013) in FreeSurfer (v5.2; Fischl et al., 2002; Glasser et al., 2013; Marcus et al., 2013), from which we downloaded ICV (FS label: *eTIV ^[1]^*) and TBV (FS label: *BrainSegNotVent*) estimates. To maximise sample size, our analyses included related individuals, but were also repeated in unrelated individuals (randomly sampled from *Family_ID* variable). We removed six participants from the sample because their TBV estimate was larger than their ICV estimate (the brain cannot be larger than the skull). Refer to an explanation for why only below 31-year-olds were included in the analysis in the ‘Deviations from the pre-registration’ section below. Final distributions of TBV, ICV, and the LBA scores are displayed in *SFig.3*.

**MRi-Share.** The MRi-Share cohort consists of 1,831 university students in Bordeaux, France, who were recruited as volunteers from the larger online i-Share study ([www.i-share.fr](http://www.i-share.fr)). Participants were aged between 18 and 35 years, and exclusion criteria were pregnancy, and characteristics that would have prohibited safe MRI scanning (e.g., claustrophobia). The MRI acquisition protocol was designed to match MRI acquisition in the UK Biobank sample (<https://www.ukbiobank.ac.uk/>). ICV (FS label: *eTIV*) was extracted using FreeSurfer v6.0 (<http://surfer.nmr.mgh.harvard.edu/>). Structural T1 and FLAIR images were processed by Tsuchida et al. (2021), where white and grey matter volume were extracted using SPM12 (<https://www.fil.ion.ucl.ac.uk/spm/>). We calculated TBV by summing white and grey matter volume. Imaging-derived phenotypes were freely available for download online ([URL](https://datadryad.org/stash/dataset/doi:10.5061/dryad.q573n5tj2)). Final distributions of TBV, ICV, and the LBA scores are displayed in *SFig.3*.

#### 2.1.2. Middle- and older-aged adults

**UK Biobank (UKB).** The UKB is a prospective population-based cohort including half a million participants in the United Kingdom (<https://www.ukbiobank.ac.uk/>). Participants were recruited as volunteers to collect health-related information from physical measurements, biological samples, and questionnaires during a planned 20 year follow-up (Sudlow et al., 2015). Brain MRI data was collected on a subsample of ~50,000 participants ('time 1'; Miller et al., 2016), which has so far been repeated for ~4,500 of those participants, about 4 years after the initial imaging visit ('time 2'; Littlejohns et al., 2020). T1 and FLAIR images were collected across three sites with identical hardware and software, and they were processed cross-sectionally on behalf of the UKB (FS v6.0; Alfaro-Almagro et al., 2018). TBV and ICV phenotypes are available for download (field IDs: TBV = 26515, i.e., FS label = *BrainSegNotVent*; ICV = 26521, i.e., FS label = *eTIV*; T1 volumetric scaling factor = 25000; CSF = 26527) (Smith et al., 2020). Data access was granted though application 10279. We removed participants with larger TBV estimates than ICV estimates (*n* = 17), and one participant as their TBV estimate was nearly five times larger than the average sample TBV. Two extreme participants (outside of 10SDs) were removed as their TBV values were only half their ICV values, and another two participants were removed because their CSF value was larger than their ICV value. Final distributions of TBV, ICV, and the LBA scores are displayed in *SFig.3*.

We had pre-registered to re-process MRI data where two time points were available using the FreeSurfer longitudinal processing stream (Reuter & Fischl, 2011). This, however, was not possible because files provided field ID 20263, were incomplete for most initial neuroimaging visits (available in ~800 participants), which would have been necessary to run the FS longitudinal processing stream. We were unable to obtain complete data from field ID 20252 – which would have been necessary to run the longitudinal FS processing stream (Reuter & Fischl, 2011) – because the UKB recently changed data download permissions (as part of moving all analyses to their Research Analysis Platform). Instead, we inferred longitudinal changes from TBV estimates produced by the FS cross-sectional processing stream as was previously done in Di Biase et al. (2023). In parallel with how we treated the cross-sectional data, we also applied a 10SDs cut-off to these repeated measures which removed 6 participants (*N* _total_ = 4674). Final distributions of the atrophy scores derived from longitudinal data (‘longitudinally-observed atrophic changes’) are displayed in *SFig.9*.

**Generation Scotland; Stratifying Resilience and Depression Longitudinally (STRADL).** STRADL is a population-based cohort, originating from the Generation Scotland Scottish Family Health Study. It was collected with the aim to subtype depression using clinical, cognitive, genetic, and brain imaging assessments (Habota et al., 2021). MRI data was processed using the cross-sectional FS stream for 1,043 participants (v5.3; eTIV for ICV estimate). We removed participants with estimates of zero *mm^3^* for TBV or ICV (*n _removed_* = 11), and where ICV estimates were smaller than TBV estimates (*n _removed_* = 45), resulting in a sample of 987 participants (ages = 26 to 84 years). Final distributions of TBV, ICV, and the LBA scores are displayed in *SFig.3* and their interpretation should consider that STRADL includes both depression cases and controls.

#### 2.1.3. Older age adults

**Lothian Birth Cohort 1936 (LBC1936)**. The LBC1936 is a cohort of community-dwelling older adults, born in 1936, who have been prospectively phenotyped over the past 20 years (Deary et al., 2007; Taylor et al., 2018). Most participants had taken part in the Scottish Mental Survey in 1947. They were recruited from the Edinburgh City and surrounding Lothians’ area. LBC1936 data collection included cognitive, psychosocial and biological measures where we used assessments from 5 waves each acquired 3 years apart during an approximate duration of 9 years. Brain MRI data was acquired across 4 time points (ages 70-87 years wave 2 *N* = 629, wave 3 *N* = 428, wave 4 *N* = 319, wave 5 *N* = 304) (Taylor et al., 2018), but to match the UKB with only two available time points, we consider wave 2 (first neuroimaging visit) as time 1, and wave 5 (fourth neuroimaging visit) as time 2. MRI data was processed and cleaned by the LBC analysis team (FS v5.1.0; Wardlaw et al., 2011), and includes variables for ICV (FS label = *eTIV*), TBV (FS label: *BrainSegNotVent*), and CSF (field ID: csf_mm3_wX). Estimates of lifetime atrophy were inferred from MRI data processed with the FS cross-sectional stream, and estimates of atrophic changes were inferred from MRI data processed with the FS longitudinal stream (Reuter & Fischl, 2011). Two participants were removed because their TBV estimate was larger than their ICV estimate. Final distributions of TBV, ICV, and the LBA scores are displayed in *SFig.3*.

Details on MRI acquisition and processing software are listed per sample in STable 1.

*STable 1.* MRI scanner and acquisition information for each of the considered cohorts

| *Sample* | *MRI scanner & acquisition info* | *MRI processing software* |
| --- | --- | --- |
| Human Connectome Project (HCP) – Young adults ^1^ | 3T Skyra MRI, customized gradient (UMinn -> Wash U) | FreeSurfer ^6^  Cross-sectional FS processing stream performed |
| MRi-Share (university students; T1 & FLAIR) ^2^ | Designed to match acquisition protocol in UKB  Siemens 3T Prisma scanner with a 64-channels head coil (gradients: 80 mT/m–200 T/m/sec), in the 2-year period between November 2015 and November 2017, voxel (matrix) size: 1.0 × 1.0 × 1.0 mm^3^ (192 × 256 × 256); Key parameters: 3D MPRAGE, sagittal, *R* = 2, TR/TE/TI = 2000/2.0/880 msec  Download global estimates: [Dryad \| Data -- The MRi-Share database: Brain imaging in a cross-sectional cohort of 1,870 university students (datadryad.org)](https://datadryad.org/stash/dataset/doi:10.5061/dryad.q573n5tj2) | ICV = FreeSurfer ^6^  White and grey matter = SPM12 ^7^  Cross-sectional FS processing stream performed |
| UK Biobank (UKB) ^3^ | Data acquired across three sites (Manchester, Newcastle, and Reading) with identical hardware and software  SIEMENS MAGNETOM Skyra syngo MR D13  Voxel size:1.6×1.6×1.6 mm | FreeSurfer ^6^  Cross-sectional FS processing stream performed; longitudinal processing impossible due to incomplete data |
| Lothian Birth Cohort 1936 (LBC1936) ^5^ | GE Signa LX 1.5T Horizon HDx clinical scanner (General Electric, Milwaukee, WI) with a manufacturer supplied 8-channel phased array head coil. For T1-weighted images (3D IR-Prep FSPGR), 160 coronal slices were acquired, with a field of view of 256 mm and a matrix size of 192 x 192 pixels giving a resolution of 1 x 1 x 1.3 mm3. The repetition time was 10 ms, echo time was 4 ms and inversion time was 500 ms. | FreeSurfer ^6^  Cross-sectional and longitudinal FS processing stream performed |
| STRADL | Two testing sites: Aberdeen and Dundee **Aberdeen**: 3T Philips Achieva TX-series MRI system scanner (Philips Healthcare, Best, Netherlands) with a 32-channel phased-array head coil and a back facing mirror (software version 5.1.7; gradients with maximum amplitude 80 mT/m and maximum slew rate 100 T/m/s)  **Dundee**: Siemens 3T Prisma-FIT scanner (Siemens, Erlangen, Germany) with 20 channel head and neck phased array coil and a back facing mirror (Syngo E11, gradient with max amplitude 80 mT/m and maximum slew rate 200 T/m/s). | FreeSurfer ^6^  Cross-sectional FS processing stream |

References: ^1^ Van Essen et al. (2012); ^2^ Tsuchida et al. (2021); ^3^ [URL](https://www.fmrib.ox.ac.uk/ukbiobank/protocol/V4_23092014.pdf), Miller et al. (2016); ^5^ [URL](https://www.ed.ac.uk/lothian-birth-cohorts/data-access-collaboration), Wardlaw et al. (2011); ^6^ FreeSurfer reference: Fischl et al. (2002), ^7^ [URL](https://www.fil.ion.ucl.ac.uk/spm/)

#### 2.1.4. Phenotype definition for health-related traits in UKB and LBC

To characterise lifetime brain atrophy (LBA) and to compare it to longitudinally-observed atrophic changes, we calculated associations with ageing-related traits. They were derived in the UKB and LBC1936 based on the field IDs listed in STable 2.

*STable 2*. Health outcome definitions considered in this study

| Trait | UK Biobank item (field ID) | LBC (variable name) |
| --- | --- | --- |
|  | - | Brain atrophy visual rating scales (Farrell et al., 2009)   - **Superficial brain atrophy:** width of the cortical sulci, the width and shape of the cortical gyri and the space between the brain and the skull - **Deep brain atrophy:** width of the ventricles |
| General factor of cognitive ability modelled using confirmatory factor analysis in the lavaan R package (Rosseel, 2012) | We modelled a latent factor underlying the listed cognitive tests where we consider assessments at the initial imaging visit   - Reaction Time (20023) - Number span (4282) - Fluid intelligence (20016) - Trail making B (6350) - Matrix pattern (6373) - Tower task (21003) - Digit-symbol substitution (23324) - Pairs matching (399) - Prospective memory (20018) - Paired associates (20197) - Picture vocabulary (26302) | **Waves 1- 5:** We modelled individual-level *intercepts* and *slopes* in a latent growth curve model based on the listed cognitive tests, to test baseline levels of cognitive ability (intercept) as well as rates of change in cognitive ability (slope) for their association with brain atrophy measures ([URL](https://www.ed.ac.uk/sites/default/files/atoms/files/r_factor_of_curves_g_txt.txt))   - WMS III - Logical Memory - WMS III - Spatial Span - WMS III - Verbal Paired Associates - WMS III - Symbol Search - WMS III - Digit Symbol Coding - Simple and 4-choice reaction time - Inspection Time - WAIS III - Matrix Reasoning - Verbal Fluency C,F,L - WAIS III - Digit Span Backwards - WAIS III - Block Design - WTAR (Wechsler Test of Adult Reading) - NART (National Adult Reading Test |
| Dementia status | Lifetime all-cause dementia proxy phenotype   - ICD-10 diagnoses: F00, F01, and F03, G30 (41270) - Self-reported illnesses: dementia/alzheimers/cognitive impairment (20002; code = 1263) - Source of all cause dementia report (42019) - Contributing causes of death: F00, F01, and F03, G30 (40001 & 40002) - Illnesses of father and mother: Alzheimer's disease/dementia (20107 & 20110, code = 10) - Date F00 first reported (130836) | - Lifetime clinical dementia diagnosis (*dement*), incidence across all waves |
| Apolipoprotein (APOE) status | Allele present in genotype data for SNPs *rs7412* & *rs429358* (Kuo et al., 2020) | - APOE e4 allele present (*APOEe4*) |
| Fried frailty phenotype (Fried et al., 2001) | Fried frailty definition adopted to UKB sample according to Jiang et al. (2023) at the initial imaging visit   - Weight loss (2306) - Exhaustion (2080) - Walking speed (924) - Weakness (46, 47) - Physical activity (6164, 1011) | **Waves 1-5:** We modelled individual-level *intercepts* and *slopes* of Fried frailty (as implemented in Welstead et al., 2020) in a latent growth curve model.   - Weight loss (*weight*; *height*) - Exhaustion: ‘I feel as if I'm slowed down’ (*HADSD4*) - Physical activity (phyactiv) - Walking speed (sixmwk) - Weakness: grip strength (griprh) |
| Type 2 diabetes | Lifetime Type 2 diabetes (as implemented in Fürtjes et al., 2022)   - Self-reported illness (20002) - ICD9 diagnoses (41203 & 41205) - ICD10 diagnoses (41202 & 41204) - Cause of death (40001) - Diagnosis by doctor (2443) | Lifetime diagnosis of diabetes (*diab*), incidence across all waves |
| Hypertension | Lifetime hypertension diagnosis (as implemented in Cox et al., 2019)   - Self-reported illness (20002) - ICD9 diagnoses (41203 & 41205) - ICD10 diagnoses (41202 & 41204) - Cause of death (40001) - Diagnosis by doctor (2443) - Source of report I10 (121387) | Lifetime diagnosis of hypertension, incidence across all waves   - Self-reported GP diagnosis: *hibp* |
| Smoking (packyears) | *Packyears* at initial brain imaging visit   - Pack years of smoking (20161) | - *Packyears* of smoking at wave 5 (*smokpackyrs*) |
| Body mass index | Initial imaging visit: Body mass index calculated as weight/height^2^ (kg/m^2^)  Calculated mean of   - Body composition by impedance (23104) - Body mass index (21001) | **Waves 1-5**: We modelled individual-level intercepts and slopes of body mass index in a latent growth curve model (*bmi*) |
| Brain age gap | Initial imaging visit: We modelled brain age as implemented by Cole et al. (2018) using brainAgeR [URL](https://github.com/james-cole/brainageR) ^[2]^ Brain age gap = chronological age – brain predicted age (x-1) | **Wave 2:** We modelled brain age as implemented by Cole et al. (2018) [URL](https://github.com/james-cole/brainageR)  Brain age gap = chronological age – brain predicted age (x-1) |
| Stroke | Lifetime stroke diagnosis (as implemented in Fürtjes et al., 2022)   - Self-reported illness (20002) - ICD9 diagnoses (41203 & 41205) - ICD10 diagnoses (41202 & 41204) - Cause of death (40001) - Diagnosis by doctor (2443) - Source of stroke report (42007) | Lifetime stroke lesions: any silent stroke lesions identified from neuroimage by a neuroradiologist (Muñoz Maniega et al., 2019) (*stroke_mask*) |

### 2.2 Statistical analysis

#### 2.2.1 Computational approaches to estimate brain atrophy

We investigated different computational classes of brain atrophy calculated from either one MRI scan or two repeated MRI scans acquired some years apart. Approaches using one scan leveraged ICV to approximate premorbid brain size. Hereafter, we will refer to atrophy inferred from one cross-sectional MRI scans as LBA, and atrophy inferred from two longitudinal MRI scans as (within-person) *observed atrophic changes*. We will report descriptive statistics for raw measures (*STable 1*), but associations were calculated for standardised measures (mean = 0, SD = 1).

##### 2.2.1.1 LBA: Computing brain atrophy cross-sectionally using a single MRI scan

**Difference score.** The cross-sectional difference score is computed on an individual-level from a cross-sectional MRI scan as the difference between ICV and TBV (*ICV-TBV*).

**Ratio score.** The cross-sectional ratio (or proportional) score is computed on an individual-level from a cross-sectional MRI scan as the ratio between TBV and ICV (*TBV/ICV*).

**Residual score.** The cross-sectional residual score is computed as the TBV-associated residuals from the regression between TBV and ICV (*TBV~ICV*). Raw residual scores are interpreted as the difference between an individuals’ observed TBV and their predicted TBV given their ICV (i.e., a negative value means an individuals’ TBV is smaller than expected given their ICV).

**Cerebrospinal fluid (CSF).** A raw measure of CSF may capture inter-individual variance strongly related to variance indexed by the difference, ratio, and residual scores, considering TBV volume equals ICV volume minus CSF volume. To evaluate the potential of CSF to index brain atrophy, phenotypic and genetic correlations between LBA and TBV and ICV were also repeated for a measure of CSF (phenotypic results section 3).

##### 2.2.1.2 Observed atrophic changes: Computing brain atrophy longitudinally using two repeated MRI scans

**Difference score.** The longitudinal difference score is computed on an individual-level from the difference between two estimates of TBV that were obtained from two time-shifted MRI scans (*TBV _time 1_ – TBV _time 2_*).

**Ratio score.** The longitudinal ratio (or proportional) score is computed on an individual-level from the ratio between two estimates of TBV that were obtained from two time-shifted MRI scans (*TBV _time 2_/TBV _time 1_*).

**Residual score.** The longitudinal residual score is computed as the TBV *_time 2_* associated residuals from the regression between TBV *_time 2_* and TBV *_time 1_* (*TBV _time 2_ ~ TBV _time 1_*).

#### 2.2.2 Phenotypic analyses

**Phenotypic correlations and associations.** Anytime the manuscript reports correlations in the first phenotypic part of the manuscript, they were obtained via simple Pearson’s correlations. Associations between brain atrophy and ageing-related traits (results presented in *Fig.2* in main manuscript) were obtained with linear regressions when the outcome trait was continuous, with logistic regression when the outcome trait was binary, and with hurdle regression when the outcome trait was zero-inflated count data. Variance explained (*R^2^*) in the outcome trait was calculated as the linear regression coefficient squared, *R^2^* was obtained with Nagelkerke’s *R^2^* for logistic regression using the fmsb package in R ([URL](https://doi.org/10.32614/CRAN.package.fmsb)), and a maximum likelihood pseudo *R^2^* for the hurdle regression using the pscl package in R ([URL](https://rdocumentation.org/packages/pscl/versions/1.5.9)). If *beta* values are reported (as opposed to correlation values), any predictor and outcome variables were standardised to a mean of zero and a standard deviation of one. All analyses were performed in R v4.2.2, and all plots were produced in ggplot2.

#### 2.2.3 Genetic analyses

**Genetic data cleaning.** Participants were excluded when labelled outliers in heterozygosity and missingness by the UKB analysis team (het_missing_outlier), and when they self-reported non-European ancestry or were missing this self-report information. European ancestry was also determined based on 4-means clustering of 40 genetic principal components. Relatedness was identified in ukbtools (Hanscombe et al., 2019) using the default cut-off > 0.0884 King coefficient corresponding to 3rd degree relatedness. The sex check was performed in PLINK based on the observed number of heterozygote variants from that expected under Hardy-Weinberg equilibrium. Autosomal SNPs were filtered for missing genotype data at a rate of 0.02, minor allele frequency > 0.01 and Hardy-Weinberg equilibrium exact test at 0.00000001. Genotypes were imputed by the UKB analysis team with reference to the Haplotype Reference Consortium (HRC) and UK10K haplotype resource ([: Category 100319 (ox.ac.uk)](https://biobank.ndph.ox.ac.uk/showcase/label.cgi?id=100319)).

*STable 3.* Cleaning of 45,598 participants with available genetic and neuroimaging data

| **Data cleaning step** | **Removed** | **Remaining** |
| --- | --- | --- |
| Remove outliers in heterozygosity and missingness | 83 | 45,515 |
| Determine European ancestry by 4-mean clustering | 1,332 | 44,183 |
| Filter for self-reported Europeans | 147 | 44,045 |
| Remove participants with missing self-reported ancestry | 9 | 44,036 |
| Remove related individuals | 622 | 43,414 |
| Sex check | 22 | 43,132 |
| Missing covariate info removed by REGENIE | 282 | 43,110 |

*STable 4.* Cleaning of 805,161 SNPs available in UKB genotype data (incl. sex chromosome)

| **Data cleaning step** | **Removed** | **Remaining** |
| --- | --- | --- |
| Missing genotype data (0.02) | 104,462 | 700,699 |
| Minor allele frequency (0.01) | 103,158 | 597,541 |
| Hardy-Weinberg exact tests | 11,483 | 586,058 |
| Autosomal SNPs | 16,231 | 569,827 |

**SNP-heritability.** We calculated SNP-based heritability (*h^2^*) for TBV, ICV, CSF and all LBA measures using Genome-wide Complex Trait Analysis (GCTA v1.94.1; Yang et al., 2011) in UKB genotype data with a cryptic relatedness cut-off at 0.025 (*N* = 38,624). The following covariates were included age, sex, assessment month, assessment site, x, y, and z coordinates, genotyping array, genotyping batch, and 40 genetic PCs. We did not include age-squared or product-wise age-by-sex as additional covariates because they were nearly perfectly correlated with age (*r* = 0.99) and sex (*r* = 0.89), respectively. Our reasoning was that including near identical variables would cause issues of multicollinearity which would produce unreliable estimates. Estimates of SNP-heritability were derived from a model with two variance components (common variants, and error), and estimates of SNP-by-age were derived from a model with three variance components (common variants, SNP-by-age interaction, and error).

**GWAS analysis.** GWAS were performed for TBV, ICV, CSF, and lifetime atrophy inferred with the residual, ratio and difference method. Genome-wide SNP-based associations were calculated in a mixed linear model using REGENIEv3.5 (Mbatchou et al., 2021), including the following nuisance covariates: sex (field ID: 31), acquisition site (field ID: 54), acquisition time (field ID: 53), scan positions x,y,z (field IDs: 25756, 25757, 25758), 40 genetic PCs, genotyping array and genotyping batch. Given the wide age range in UKB, age (field ID: 21022) was also included as a covariate to ensure that atrophy levels were comparable between older and younger participants. Age-adjusted analyses should capture whether individuals show larger or smaller levels of atrophy in comparison with other individuals of similar ages and circumstances. We did not include age-squared or product-wise age-by-sex as additional covariates because they were nearly perfectly correlated with age (*r* = 0.99) and sex (*r* = 0.89), respectively. Our reasoning was that including near identical variables would cause issues of multicollinearity which would produce unreliable estimates. Additional GWAS were performed for males (*n* = 20,453) and females (*n* = 22,657) separately. Age-by-SNP interactions were also tested in REGENIEv3.5 using the --interaction flag.

**Genetic correlations.** Using bivariate linkage disequilibrium score regression (LDSC; Bulik-Sullivan et al., 2015), we used GWAS summary statistics calculated in the steps above to quantify genetic correlations with other structural neuroimaging traits. SNP effects associated with the residual and ratio score were flipped (i.e., multiplied by -1) so that a positive association can be interpreted as increasing the risk for LBA. We assessed the genetic correlations of these traits with other MRI-based phenotypes: TBV (Zhao et al., 2019), TBV (BrainSegNotVentSurf, ID: 0167; Smith et al., 2021), ICV (Adams et al., 2016), brain age (Kaufmann et al., 2019), general dimensions of brain morphometry shared across 83 brain-wide volumes (Fürtjes et al., 2023), brain ventricular volume (Vojinovic et al., 2018). Our pre-registration also included longitudinal changes in brain structure (Δ total brain; Brouwer et al., 2022), but this set of summary statistics produced a negative heritability estimate and was not suitable for LDSC. Finally, we used GWAS summary statistics of four neurodegenerative disorders associated with aging that were well powered enough to perform LDSC (i.e., *h^2^* Z-score > 4). The traits were Alzheimer’s disease (Kunkle et al., 2019), Alzheimer’s disease and related dementias (Bellenguez et al., 2022), Parkinson’s disease (Nalls et al., 2019), Amyotrophic lateral sclerosis (van Rheenen et al., 2021). We included them post-hoc to provide a commentary on the current relevance of these genetic information to clinical outcomes.

**GWAS-by-subtraction model.** To obtain the TBV-associated residuals of ICV, we fitted a GWAS-by-subtraction model in GenomicSEM (Grotzinger et al., 2019). The lavaan syntax is printed below. In the main manuscript, we report the correlation between those TBV-associated residuals (‘*Atrophy*’) and the LBA residual score (‘*resid*’) described in the main manuscript.

BaseICV=~NA*TBV + ICV
Atrophy=~NA*TBV
Atrophy ~~ 1*Atrophy
BaseICV ~~ 1*BaseICV
Atrophy~~0*BaseICV
ICV ~~ 0*TBV
ICV ~~ 0*ICV
TBV ~~ 0*TBV
Atrophy ~~ resid
BaseICV ~~ resid

**Functional mapping and annotation (FUMA).** FUMA v.1.5.2 with default settings (Watanabe et al., 2017; https://fuma.ctglab.nl) was utilised to identify the genomic risk loci captured by our GWAS summary statistics. Independent significant SNPs were identified based on a genome-wide significance threshold (*p* < 5x10^-8^) and independence of other variants at *r^2^* < 0.6. LD structure was calculated from the UKB release2b 10k European reference panel population. From the set of independent significant SNPs, FUMA identified lead SNPs independent of one another at *r^2^* < 0.1. If identified SNPs were located close to another (<250kb), they were merged into one genomic risk locus, which means each locus can contain multiple independent significant SNPs and multiple lead SNPs. Gene prioritisation was performed based on both positional mapping (max. distance 10 kb), and expression quantitative trait loci (*eQTL*) mapping to identify whether independent significant SNPs from the GWAS (and their SNPs in LD) are known *eQTL*s in specific tissue types (refs. PsychENCODE, BRAINEAC, GTEx v8 Brain).

### 3.1 Deviations from the pre-registration

#### 3.1.1 Restricting the HCP cohort to below 31-year-olds

We had pre-registered (<https://osf.io/gydmw/>) that young cohorts such as the HCP and MRi-Share should demonstrate no, or very weak associations between age and both ICV and LBA. In this project, we understand ICV to be constant across the lifetime, which is why we use it in this project to approximate premorbid brain size, i.e., the size of a participant’s brain prior to any neurodegeneration. Indeed, there is evidence that ICV should be largely stable across the lifespan, but minor age-related changes can occur due to skull and meninges thickening (Caspi et al., 2020; Nerland et al., 2022; Royle et al., 2013). Previous studies have also delivered evidence for effects of ICV measurement method and study population, whereby older cohorts tend to have somewhat smaller skulls than younger cohorts (Caspi et al., 2020; Ma et al., 2019; Nerland et al., 2022).

Contrary to our expectation of no or negligible ICV-age correlations (*r* < |0.1|), the HCP data, as downloaded, showed a moderate ICV-age correlation of *r* = -0.2 (*p* = 4.2x10^-11^). This ICV-age correlation was even stronger than the TBV-age correlation in the same sample (*r* = 0.16; *p* = 4.9x10^-8^). Implausibly, age was negatively associated with more LBA in this young sample where atrophy was derived from ICV and TBV measures using three computational methods. That is, older HCP participants had less brain atrophy (i.e., larger values in the difference score, smaller values in the ratio score) than younger participants which is not sensible, and we suggest must be a sample artifact rather than an interpretable finding. Note, however, that age was uncorrelated with brain atrophy derived using the residual method where the atrophy score is independent of ICV by design (*Methods Figure 1* below).

In the process of making sense of these unexpected correlational patterns, we excluded related individuals (n = 445), and adjusted for batch effects, none of which reduced, or substantially altered either the ICV-age correlation, or the LBA-age correlation. We had anticipated in our pre-registration that there may be unexpected correlations between age and ICV (as well as the atrophy measures), which could be driven by few older-age outliers. To test for this, we successively reduced the sample age (equivalent procedure to analyses presented in *Fig.3D-G*) by applying maximum-age cut-offs to obtain a subsample of individuals in which age and ICV are negligibly correlated below our pre-registered but arbitrary cut-off of *r* < |0.1|. Indeed, the strength of the ICV-age correlation was moderated by sample age, and excluding all participants > 31-years reduced the ICV-age correlation to *r* < 0.1 (*Methods Figure 2 below*). This indicates a cohort effect whereby older participants in the HCP have systematically smaller skulls than younger participants. *Methods Figure 3* (below) illustrates that the HCP restricted to below 31-year-olds produces a negligible age-ICV correlation (*r* < |0.1|), as well as non-significant lifetime atrophy-age correlations, which is in keeping with our expectations for young adults (outlined in the pre-registration). Hence, all analyses presented in the main manuscript were calculated in the HCP participants below the age of 31 years.

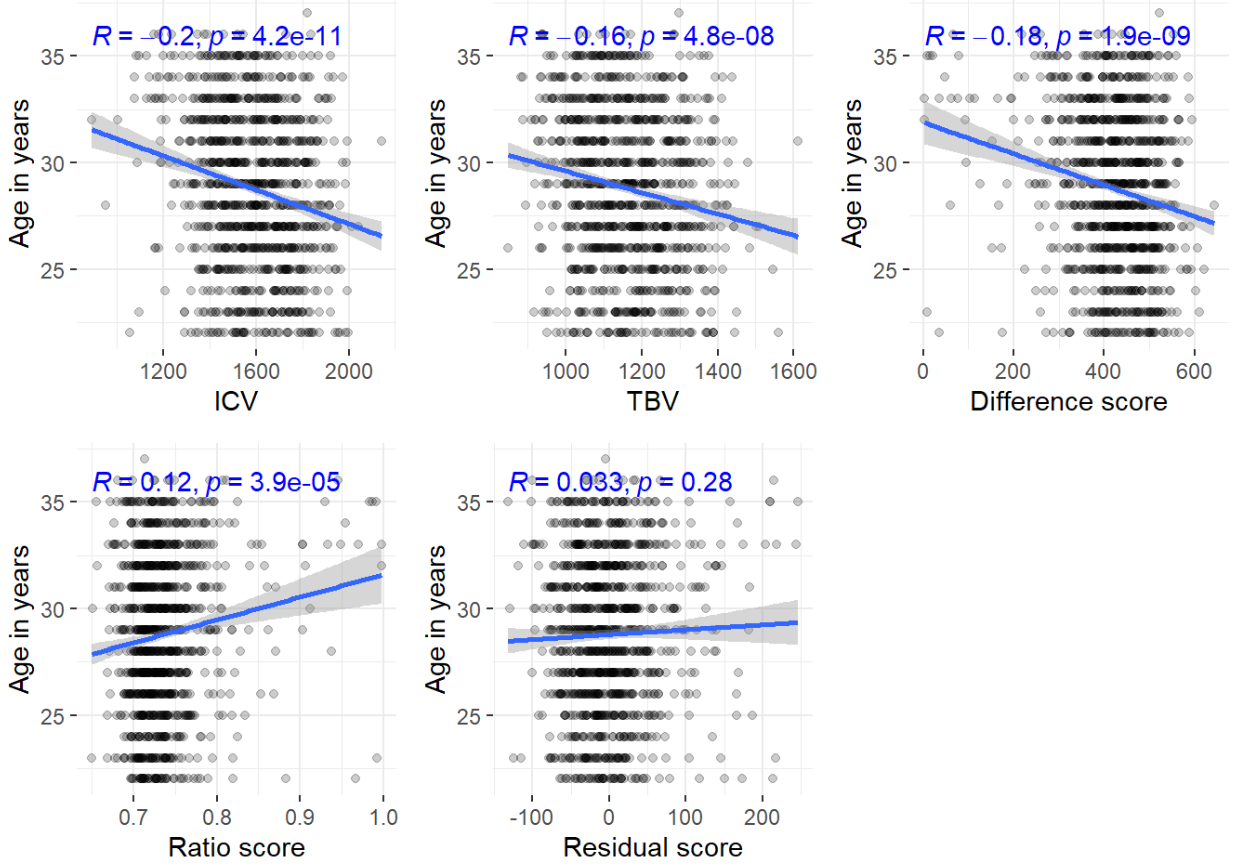

*Methods Figure 1.* Age correlations with ICV, TBV and the three atrophy scores including full HCP sample as downloaded.

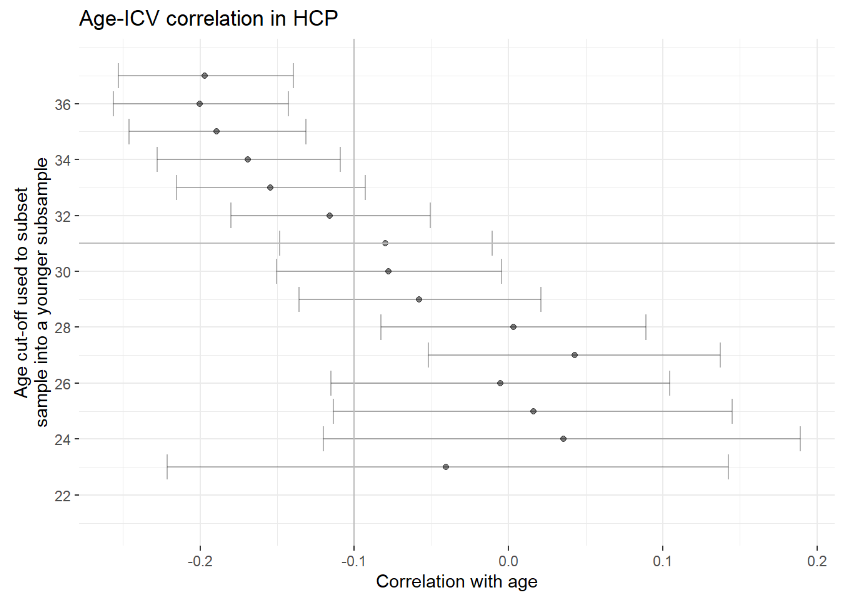

*Methods Figure 2.* Age ICV correlations in the HCP. *Y*-axis indicates the maximum age of each considered subsample

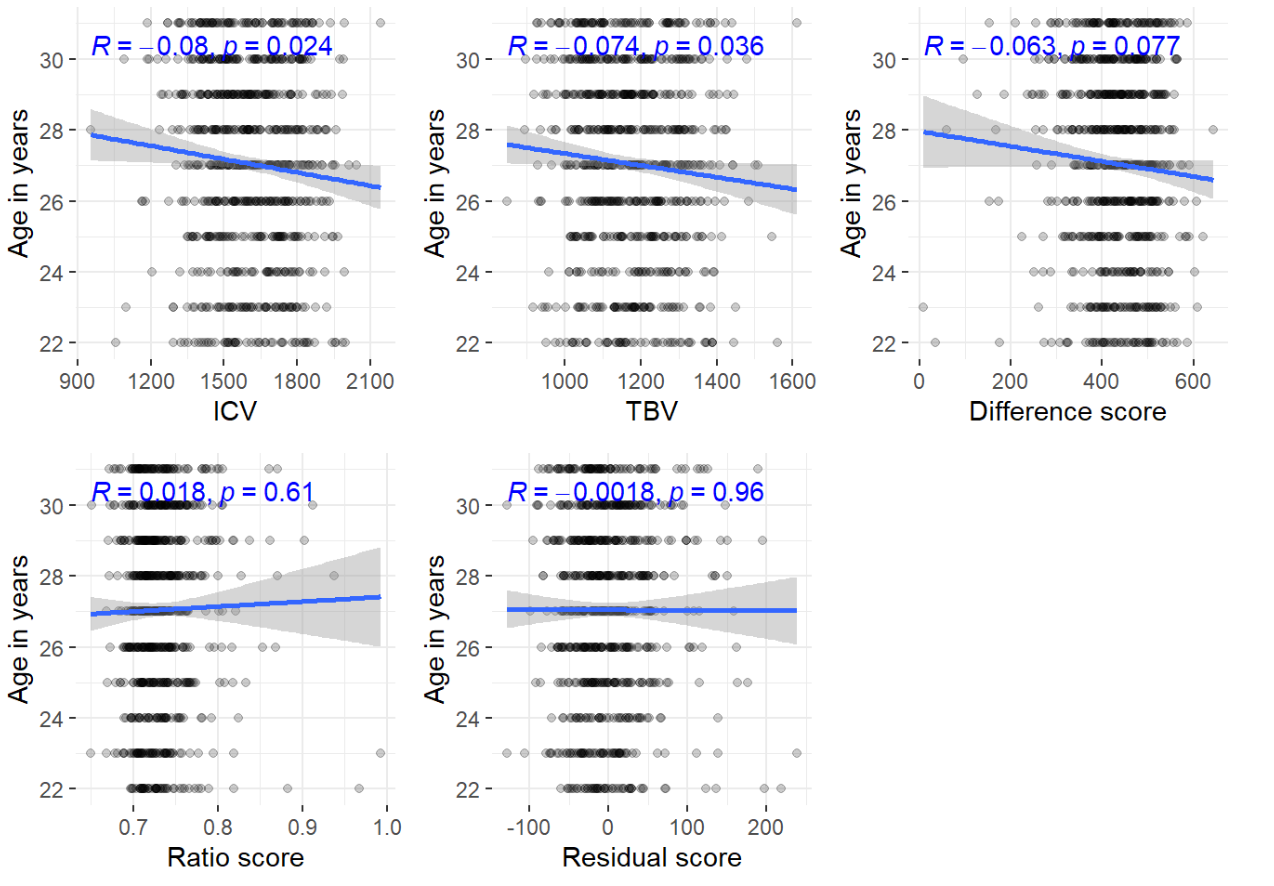

*Methods Figure 3.* Age correlations with ICV, TBV and the three atrophy scores including sample of HCP participants below the age of 31 years

#### 3.1.2 Pre-registered Aim 2; sample specificity of LBA norms and the role of ICV

Our pre-registered Aim 2 of this study had been designed to ensure fair comparisons between cross-sectionally- ‘estimated’ lifetime atrophy and longitudinally- ‘observed’ atrophic changes, where the former will have occurred across an entire *lifetime* and the latter can only span a time window between the two measured time points. We outlined the following analysis plan in order to approximately equate those two timelines: We planned to identify a pre-neurodegenerative subsample of the UKB that appeared to have no brain atrophy at the initial neuroimaging visit (according to one single MRI scan at initial neuroimaging visit). We had theorised that this sample may then demonstrate some loss in TBV between the first and the second neuroimaging visit, which would mean that timelines for lifetime atrophy and longitudinal atrophic changes should be more similar than if comparing estimated LBA with change in people who have already atrophied to some extent. As there is no objective criterion to determine whether a participant may fall into this pre-neurodegenerative category, we pre-registered to derive cut-off thresholds from independent, young cohorts (i.e., HCP and MRi-Share), where it is reasonable to assume that participants have not yet had meaningful neurodegeneration (mean age in HCP = 27 years, mean age in MRi-Share = 22 years).

Specifically, we assumed that young participants’ brains within 2 standard deviations in their difference, ratio, and residual scores to be representative of healthy-looking brains. We aimed to carry those raw 2SD cut-off values over to the UKB, to identify a subsample of participants with pre-neurodegenerative brains. Applying cut-offs derived from young cohorts to the older UKB cohort was intended to capture UKB participants whose brain atrophy scores implied similarly healthy-looking brains to those in the younger cohorts HCP and MRi-Share.

However, efforts to conduct this analysis failed because raw lifetime atrophy scores were highly sample-specific, and therefore not sensibly transferable between samples. This was likely driven by the fact that ICV measures were strongly discordant between samples [ICV means (SDs) in *mm^3^*: MRi-Share = 1568.07 (140), HCP = 1606.3 (170), UKB = 1555.18 (150), LBC = 1396.53 (140)]. A cross-sample evaluation of raw atrophy scores would have misleadingly implied that older participants’ brains scanned in UKB and LBC looked just as healthy as young participants’ brains scanned in HCP and MRi-Share. That is, the raw difference between TBV and ICV was approximately the same for younger [Mean (SD) difference in *mm^3^*: HCP = 432.49 (79.34); MRi-Share = 436.18 (48.29)] as for older samples [Mean (SD) in *mm^3^*: UKB = 369.24 (69.44); LBC = 385.31 (86.88); *p* = 1]. This was the same for the raw ratio of ICV to TBV [Mean (SD) ratio: HCP = 0.73 (0.04); MRi-Share = 0.72 (0.02); UKB = 0.76 (0.03); LBC = 0.73 (0.05)] (*SFig.1-2*). This strong sample mismatch meant that our pre-registered Aim 2 analyses were not feasible or interpretable.

#### 3.1.3. Moderation analysis

To better understand the relationship between LBA and longitudinal atrophic changes, we conducted a moderation analysis to investigate whether the association between observed atrophic changes (capturing change between time 1 and time 2) and LBA (at time 2), is moderated by the amount of LBA that occurred prior to a participants’ first MRI scan (at time 1). We hypothesised that the strongest correlation between those measures would occur when participants exhibit minimal atrophy at time 1, in which case atrophic changes and lifetime atrophy at time 2 would cover approximately the same timeline. Conversely, when the brain had already substantially atrophied prior to time 1, observed atrophic changes would likely capture fewer changes compared with lifetime measures because the latter should reflect the entire span of atrophic effects ever experienced – assuming that brain atrophy declines continually and linearly. This analysis was conceived and pre-registered as a sensitivity analysis to the failed Aim 2 analyses reported above, and was therefore not reported in the main manuscript.

1. *eTIV*, estimated Total Intracranial Volume, is computed from the extent of scaling of the brain image performed by FreeSurfer in the talairach.xfm file. It is *not* the volume of all voxels in the aparc+aseg.mgz file, which would be stored in the *BrainSeg* variable. Investigations in our team have shown that eTIV overestimates intracranial volume compared with BrainSeg, but eTIV and BrainSeg tend to correlate relatively strongly (*r* = 0.88). The MRi-Share resource provides eTIV but not BrainSeg, and because this project requires consistent use of measures across samples, we used eTIV instead of BrainSeg in all samples.
2. We obtained brain age estimates that were on average ~15 years older than the participants chronological ages. To still obtain a brain age gap estimate centred around zero, we deducted the mean value of the brain age gap (M = 14.96) from each participants’ value. For more intuitive interpretation, brain age gap estimates were flipped (x-1) so larger values represent older-appearing brains.
